# Mechanism of phospholipid transport to the bacterial outer membrane by TAM

**DOI:** 10.64898/2026.03.22.713439

**Authors:** Alanah G Eisenhuth, Lachlan S R Adamson, Catherine Zhang, Ghaeath S K Abbas, Rachel A North, Denisse L Leyton, Harris D Bernstein, Simon H J Brown, Alastair G Stewart, Constance B Bailey, Anthony S Don, Aidan B Grosas, Matthew Thomas Doyle

## Abstract

Gram-negative bacteria transport phospholipids from the inner membrane (IM) to the outer membrane (OM) via poorly understood processes. These processes are essential for cell growth and the establishment of an antibiotic-resistant barrier. Here, we conducted single-particle cryo-electron microscopy, *in vivo* functional assays, and lipidomics to investigate the role of the “translocation and assembly module” (TAM) in lipid transport. We found that the OM-embedded subunit TamA anchors the IM-embedded bridge-like subunit TamB to the OM by forming a functional stable hybrid-barrel structure with the highly conserved C-terminal domain of unknown function 490 (DUF490). Using *in vivo* disulfide-tethering experiments we found that a highly conserved amphipathic helix within TamB DUF490 is important for TAM to function in OM maintenance. We also found that TamB DUF490 forms a β-taco channel containing lipid-like densities and that the lipophilic property of the channel is important for TAM to maintain the levels of cardiolipin in the OM. Not only do our data support a novel model in which TAM acts to direct specific lipid classes into the OM, but it also supports the notion that TamB is a bacterial evolutionary prototype of a structurally homologous superfamily of eukaryotic bridge-like lipid transfer proteins.

## INTRODUCTION

The translocation of lipids between intracellular compartments is required for life but the mechanisms by which this occurs are poorly understood. For instance, how of Gram-negative bacteria transport lipids post-synthesis from the inner membrane (IM) to the outer membrane (OM) has remained a prevailing mystery. In eukaryotes, a surge of recent studies has begun to reveal the widespread importance of bridge-like lipid transfer (BLT) proteins that facilitate bulk lipid transport reactions^1–7^. BLT proteins appear to be ubiquitous and possess N-terminal Chorein domains that extend to form elongated β-taco structures with lipophilic channels critical for lipid transfer between organelles (e.g. ER to plasma membrane, mitochondria, peroxisomes, autophagosomes, etc.) as observed in organisms such as yeast, fruit fly, and implicated in several human diseases^2–4,8^. Remote homology searches have recently identified the presence of BLT proteins in bacteria (also known as the AsmA superfamily) and the importance of these β-taco proteins in lipid transport has subsequently emerged^9–17^. *E. coli* encodes several BLT proteins including TamB, YhdP, and YdbH that bridge the periplasmic space between the inner and OMs and together are essential for cell viability^10,13,18^. Strains that have one or two of these genes deleted become sensitive to detergents, bile salts, and a range of antibiotics, indicative of a severe defect in the OM permeability barrier^10,11,13^. Deletion/depletion of TamB, YhdP, and YdbH results in a reduction in OM phospholipids, arrest of cell elongation, and cell death^10,13^. The mechanism by which these proteins direct lipid transport, and whether specific lipid substrates are preferentially transported by specific bridge proteins, remains to be determined.

Given the presence TamB in all Gram-negative bacteria, and the recent identification of TamB homologs in plants, the TamB family might represent the most evolutionarily ancient form of BLT β-taco proteins^5,19–22^. TamB family proteins have an IM-embedded N-terminal helix and Chorein domain, a lipophilic β-taco that spans the periplasm, and are easily identified based on the presence of a C-terminal domain of unknown function 490 (DUF490) that possesses unusually high sequence conservation^17,19,23,24^ (**Fig. 1a**). An early solved fragment of the *E. coli* DUF490 showed the presence of detergent-like densities within the lipophilic β-taco and a pull-down experiment using a larger fragment of TamB recently reported that the protein can bind to phosphatidylethanolamine (PE) and phosphatidylglycerol (PG) lipid classes^23,25^. In *E. coli,* the TamB DUF490 binds to an outer membrane protein (OMP) called TamA to form a complex known as the translocation and assembly module (TAM)^24,25^. TamA possesses three N-terminal periplasmic domains (POTRA1-3) and a C-terminal OM-embedded β-barrel domain^26^ (**Fig. 1a**). TamA is homologous to BamA, the major subunit of the β-barrel assembly machinery (BAM) which folds and inserts OMPs into the OM by forming hybrid-barrel intermediates with OMP substrates^27–30^. Both TamA and BamA belong to the widespread Omp85 protein superfamily that is conserved across bacteria and eukaryotes^27^. Prior to its implication in lipid transport, extensive studies have shown that the expression of TAM is required for a subset of OMPs to adopt stable folds in the OM and that this has critical impacts on bacterial virulence^24,31–34^. TamA has been proposed to function similarly to BamA for OMP folding^26^. This model is complicated by recent cryo-electron microscopy (cryo-EM) structures that suggest that the TAM complex exists as two potentially unstable populations; one in which the TamA β-barrel forms a hybrid-barrel with the C-terminus of the TamB DUF490, and another where the TamA β-barrel is free^25^. However, because this study isolated TAM in the absence of lipids, the mechanism of lipid transport and the relevance of these conformations to the function of TAM in the OM remain ambiguous.

**Figure 1.**
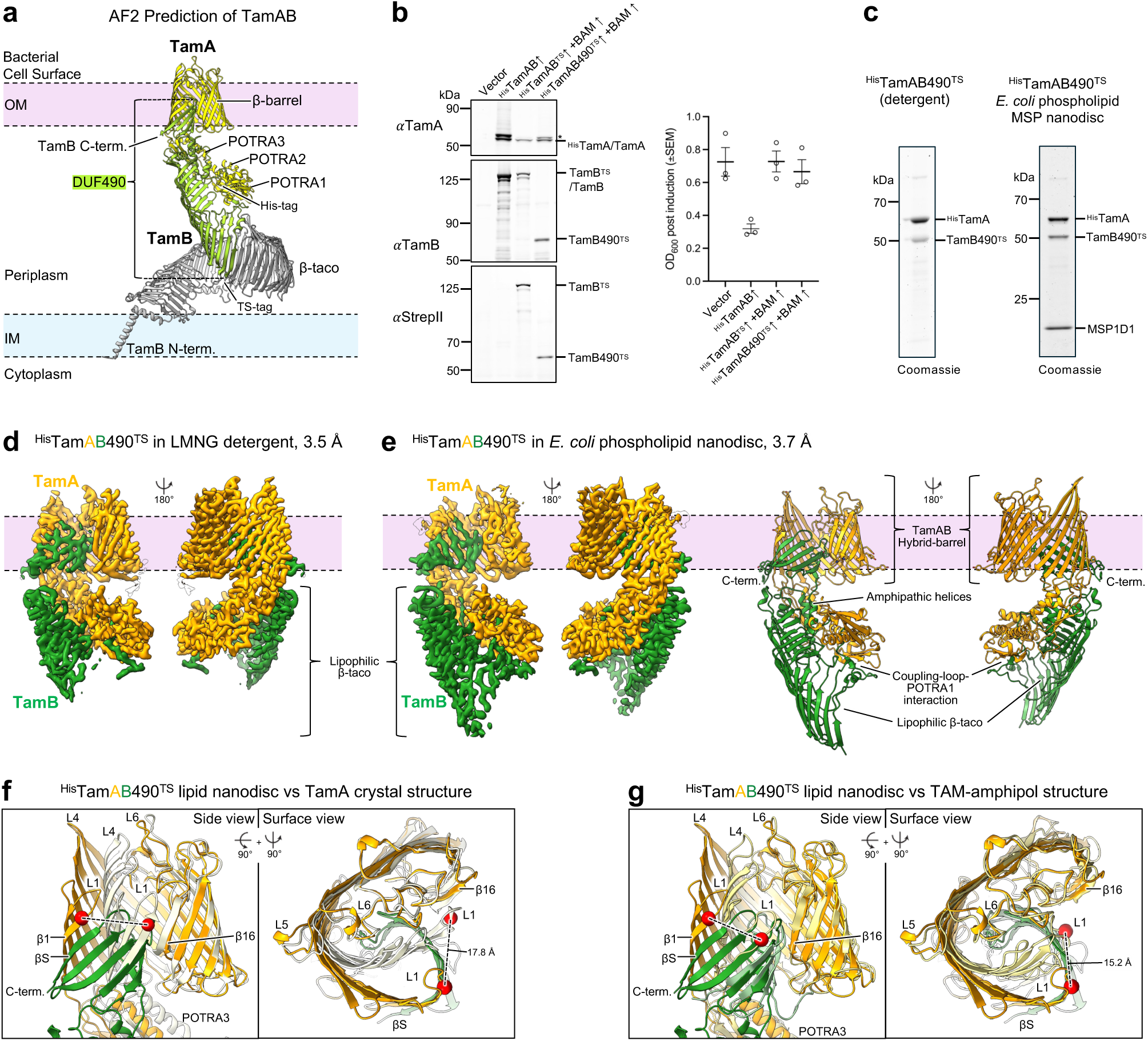
Cryo-EM structures of TamAB490 in detergent and phospholipid nanodiscs. (**a**) Structure of TAM predicted using AlphaFold2 and depicted embedded into both the inner membrane (IM) and outer membrane (OM). POTRA and β-barrel domains of TamA (yellow), and the location of an N-terminal His-tag, are indicated. The conserved TamB domain of unknown function 490 (DUF490) is colored light-green and the remaining N-terminal region of TamB is grey. The location of a Twin-StrepII (TS)-tag introduced into TamB is indicated. (**b**) *E. coli* BL21 cultures expressing ^His^TamAB, ^His^TamAB^TS^ and BamABCDE, ^His^TamAB490^TS^ and BamABCDE, or possessing an empty pTrc99a plasmid were induced with 0.4 mM IPTG at 25 ℃ for 1h. Total cell protein was probed by western immunoblotting using αTamA, αTamB, or αStrepII and culture density (OD_600_) was recorded that this time point (n = 3). TAM subunit expression in uninduced samples was also probed by immunoblot (Fig. S1). Asterisk, putative TamA prior to signal peptide cleavage. Statistical tests are in Table S2. (**c**) ^His^TamAB490^TS^ purified in LMNG detergent (left) or reconstituted into *E. coli* phospholipid membrane nanodiscs with membrane scaffold protein 1D1 (MSP1D1) (right) was resolved by SDS-PAGE. (**d**) Cryo-EM map of ^His^TamAB490^TS^ in LMNG detergent (3.5 Å average resolution). ^His^TamA and TamB490^TS^ are colored orange and dark-green respectively. (**e**) Cryo-EM map of ^His^TamAB490^TS^ in phospholipid nanodiscs (left, 3.7 Å average resolution) and model (right). (**f**) Comparison of ^His^TamAB490^TS^-nanodisc (orange, dark-green) to the crystal structure of TamA alone (white, PDB ID: 4C00^26^). Models were aligned on TamA β-barrel α-carbons Y440-L577. The difference in the position of α-carbon Y274 (red sphere, TamA β1) between each model was measured. (**g**) As in **f** except ^His^TamAB490^TS^-nanodisc and TamAB^His^-amphipol (TamA, cream; TamB, teal-green; PDB ID: 9XDC^25^). Additional comparisons in Fig. S8 and S9.

Here, we conducted cryo-EM to examine the OM portion of the TAM complex isolated within *E. coli* phospholipid membrane nanodiscs and probed the structure-function relationship of TamB DUF490 *in vivo* by disulfide-formation assays, antibiotic susceptibility, microscopy, and quantitative lipidomic analyses. We observed a single stable hybrid-barrel conformation that was significantly different from previously reported structures of TAM, and found that this state was stable and functional *in vivo*. Furthermore, we produced evidence that the positioning of highly conserved amphipathic helices between the hybrid-barrel and the β-taco is required for TAM function. Remarkably, we also observed lipid-like densities in the β-taco of TamB DUF490 reminiscent of lipids observed in eukaryotic BLT proteins^2^, and that the lipophilicity of the β-taco channel was required for maintenance of cardiolipins in the OM which strongly suggests a substrate bias for TAM. These data allowed us to propose a novel mechanistic model for the function of TamB DUF490 in lipid transport that suggests generalizable themes for the function of BLT proteins in both bacteria and eukaryotes.

## RESULTS

### Structure of TamAB in phospholipid membrane nanodiscs

We first identified DUF490-containing proteins in the UniProt database as a marker for the TamB family and found that those in bacteria can be subcategorized into three major length classes; short (1250-1300aa), medium (1450-1500aa), and long (1650-1700aa), with the *E. coli* TamB being a model representative of the short class (**Table S1**). As an aside, we found that eukaryotic DUF490-containing proteins have a much greater mean length (2150-2200aa) that might reflect the larger scales of intermembrane distances within these cells (**Table S1**). To investigate the role of *E. coli* TamAB in phospholipid transport we next isolated the complex within membrane nanodiscs for structural determination. Initially we transformed *E. coli* BL21(DE3) with p^His^TamAB for IPTG-inducible production of ^His^TamAB (N-terminal His-tag on the TamA subunit). However, we observed a significant growth defect upon induction of ^His^TamAB in liquid culture (**Fig. 1b**) and that overexpression of the complex in an *E. coli ΔtamABΔyhdP* strain is lethal (**Fig. S1**, *for genetics see below*). Western immunoblotting of total protein extracts from the BL21(DE3) cultures showed that upon IPTG-induction TamB was the target of significant endogenous degradation and that ^His^TamA accumulated as two distinct bands (**Fig. 1b**). When we probed uninduced culture samples for leaky expression, however, we did not detect the slower migrating ^His^TamA species (**Fig. S1**). Together these results might explain why endogenous TamA expression is maintained at levels 20-fold lower than BamA^35^ and suggested that ^His^TamAB overexpression can quickly overwhelm the BAM complex at the OM resulting in a backlog of unassembled TamA and TamB subunits. Consistent with our hypothesis, co-expression of both BAM and TAM from the same plasmid fully restored bacterial growth, resulted in homogenous TamA production, and significantly reduced endogenous TamB degradation (**Fig. 1b, S1**). We therefore maintained BAM co-expression in all experiments involving plasmid expression of TamAB. We also inserted a TwinStrepII (TS)-tag into a predicted turn in the TamB β-taco between residues 836/837 for detection and purification purposes. We further engineered a derivative plasmid to produce ^His^TamAB490^TS^, a version of the complex that possesses a truncated TamB subunit possessing only the DUF490-containing C-terminal half of the protein. This decision was based on (1) the incredibly high sequence conservation of DUF490 that suggested that it conducts critical steps in the mechanism of phospholipid transport (**Fig. S2**), (2) our use of AlphaFold2 (AF2) to predict the TamAB complexes of a broad range of species which suggested that the TamA-TamB(DUF490) interactions constitute the most structurally conserved and stable portion of the complex (**Fig. S3**), and (3) that the N-terminal portion of TamB was unobservable in the recently reported TamAB^His^-amphipol cryo-EM maps (only DUF490 observed) which strongly suggested that this region is unstable upon extraction of the complex from the bacterial cell envelope^25^. Finally, we purified ^His^TamAB490^TS^ in detergent micelles and reconstituted the complex into membrane nanodiscs containing *E. coli* phospholipids (**Fig. 1c**).

High-resolution structures of both ^His^TamAB490^TS^ in detergent and ^His^TamAB490^TS^-nanodiscs were solved by single-particle cryo-EM to global resolutions of 3.5 Å and 3.7 Å, respectively (**Fig. 1d, 1e, S4, S5**). Although both of our ^His^TamAB490^TS^ structures are virtually identical to each other (RMSD of 1.664 Å, α-carbons), they possess significantly different conformations to those of previously solved structures of TAM. They reveal a novel hybrid-barrel structure wherein β-strand 1 (β1) of the TamA β-barrel is bound to the C-terminal strand of TamB DUF490 that contains a “β-signal” (βS) motif (similar to the OMP β-signal for substrate recognition by BamA^29^) to accommodate six membrane-embedded β-strands of TamB (**Fig. 1e, 1f**). This hybrid-barrel conformation appears to be stabilized by electrostatic interactions observed between highly conserved residue pairs within the lumen; (1) TamA_K283_-TamB_D1252_ (TamA β-strand 2 with TamB βS) and (2) TamA_R368_-TamB_D1200_ (TamA β-strand 7 with TamB loop1) (**Fig. S2, S6, S7**). Comparison of the ^His^TamAB490^TS^-nanodisc structure to the crystal structure of TamA alone in which the β-barrel is closed through hydrogen bonding between β1-β16^26^, shows that the hybrid-barrel structure forms by a significant flattening of the N-terminal portion of the TamA β-barrel through strands β1-β8 which causes β1 to separate from β16 by ∼17.8 Å (**Fig. 1f**). Strikingly, the opening of TamA in our ^His^TamAB490^TS^-nanodisc hybrid-barrel structure is also highly similar to the open conformation of BamA observed during the folding of OMP β-barrel substrates in which a transient BamA-OMP hybrid-barrel structure is formed^29,30^ (**Fig. S8a**). Interestingly, compared to our hybrid-barrel structure in membrane nanodiscs, the reported TamAB^His^-amphipol structure^25^ possesses a highly compressed hybrid-barrel conformation in which the TamA β-barrel domain is similar to that of TamA-alone, the β1 remains close to β16 (β1 is ∼15.2 Å away from its position in our nanodisc structure) (**Fig. 1g, S8b**), and TamA_R368_ and TamB_D1200_ are separated. The difference in the position of TamA β1 between ^His^TamAB490^TS^-nanodisc and TamAB^His^-amphipol structures results in a significant displacement in the positions of POTRA1-3 and, consequently, the position of the TamB β-taco which is linked to the POTRA domains via the coupling-loop in both structures (**Fig. S9**).

Given the differences in our structures to those previously reported, we further assessed the potential for dynamic regions in ^His^TamAB490^TS^ by conducting 3D variability analysis on our particles in either detergent (**Video S1-S9**) or nanodiscs (**Video S10-S18**). In both cases we observed moderate variability in the region of the hybrid-barrel that involved the embedded TamB β-strands near TamA β16 (**Video S1-3, S10-12**) which corresponded with the distribution of local resolution observed in our maps (**Fig. S4 and S5**). We also observed an apparent tilting of the TamA β-barrel in the membrane plane in the ^His^TamAB490^TS^-nanodisc particles (**Video S10-12**). To model the apparent flexibility of the hybrid-barrel we further analyzed our ^His^TamAB490^TS^-nanodisc particles using the 3DFlex neural network^36^ which identified a series of motions wherein the C-terminal portion of the TamA β-barrel moves up and down across the z-axis of the membrane while the overall hybrid-barrel structure remains stable (**Video S19-20**). These motions, however, do not appear to correspond with a trajectory towards a compressed hybrid-barrel conformation nor the release of the TamB C-terminus from the TamA β-barrel which was observed in ∼50% of the particles for amphipol reconstituted TamAB^His^ ^25^. Instead, our results indicate that TAM forms a long-lived (stable) hybrid-barrel structure in the OM with a potential role in local membrane dynamics.

### The stable TamA-TamB hybrid-barrel is the functional state in vivo

We next obtained evidence that TAM forms a stable TamA-TamB hybrid-barrel for its function *in vivo*. To test whether TamA β1 forms a hybridization interface with TamB βS in the OM, we used our ^His^TamAB490^TS^ structures to guide the substitution of register-paired lumen-facing residues for cysteine within these strands (**Fig. 2a**). We then expressed wild-type (WT) full-length ^His^TamAB^TS^ and derivatives in *E. coli* and assessed the intermolecular proximity of cysteine-pairs by monitoring the maximal formation of disulfide-bonds *in vivo* after the addition of the thiol-specific disulfide-oxidation catalyst 4-DPS (Ox +). In these experiments we included single-cysteine controls, reduction controls, and identified intermolecular disulfide-bonds by duplex fluorescent immunoblotting. As an aside, spontaneous disulfide formation between TamA_T267C_/TamB_E1258C_ and TamA_E269C_/TamB_Q1256C_ was recently reported^25^ but the experiment did not unambiguously determine the existence of the interaction, nor the proportion of TAM complexes in a hybrid-barrel conformation. Consistent with our hypothesis, we observed remarkable levels of disulfide-bond formation between register-paired lumen-facing cysteines in ^His^TamA β1 and TamB^TS^ βS. After oxidation the majority of cellular ^His^TamA_T267C_, ^His^TamA_E269C_, and ^His^TamA_G273C_, formed high molecular weight adducts with TamB_E1258C_^TS^, TamB_Q1256C_^TS^, and TamB_D1252C_^TS^, that were detected by both anti-TamA and anti-StrepII antibodies (**Fig. 2b**, lanes 4, 6, 12). As expected, we did not observe any high molecular weight bands on immunoblots on reduced samples, nor samples from assays repeated on single-cystine controls, which together shows that the detected crosslinks are the result of site-specific disulfide-bond formation (**Fig. S10, S11)**. Although only low levels of ^His^TamA_G271C•_TamB_L1254C_^TS^ adducts were observed, this is explained by the observation that a significant fraction of TamB_L1254C_^TS^ was detected as a lower molecular weight degradation fragment (**Fig. 2b**, lanes 9, 10, **S10**). Degradation of TamB_D1252C_^TS^ was also detected but to a much lower level (**Fig. 2b**, lanes 11, 12, **S10**). We also tested an opposite-oriented ^His^TamA_T270C_/TamB_Q1256C_^TS^ pair but observed no disulfide-bond formation (**Fig. 2b**, lanes 7, 8). Together, the results strongly suggest that ^His^TamA β1 and TamB^TS^ βS hybridize to form a rigid and stable β-sheet secondary structure in the bacterial OM.

**Figure 2.**
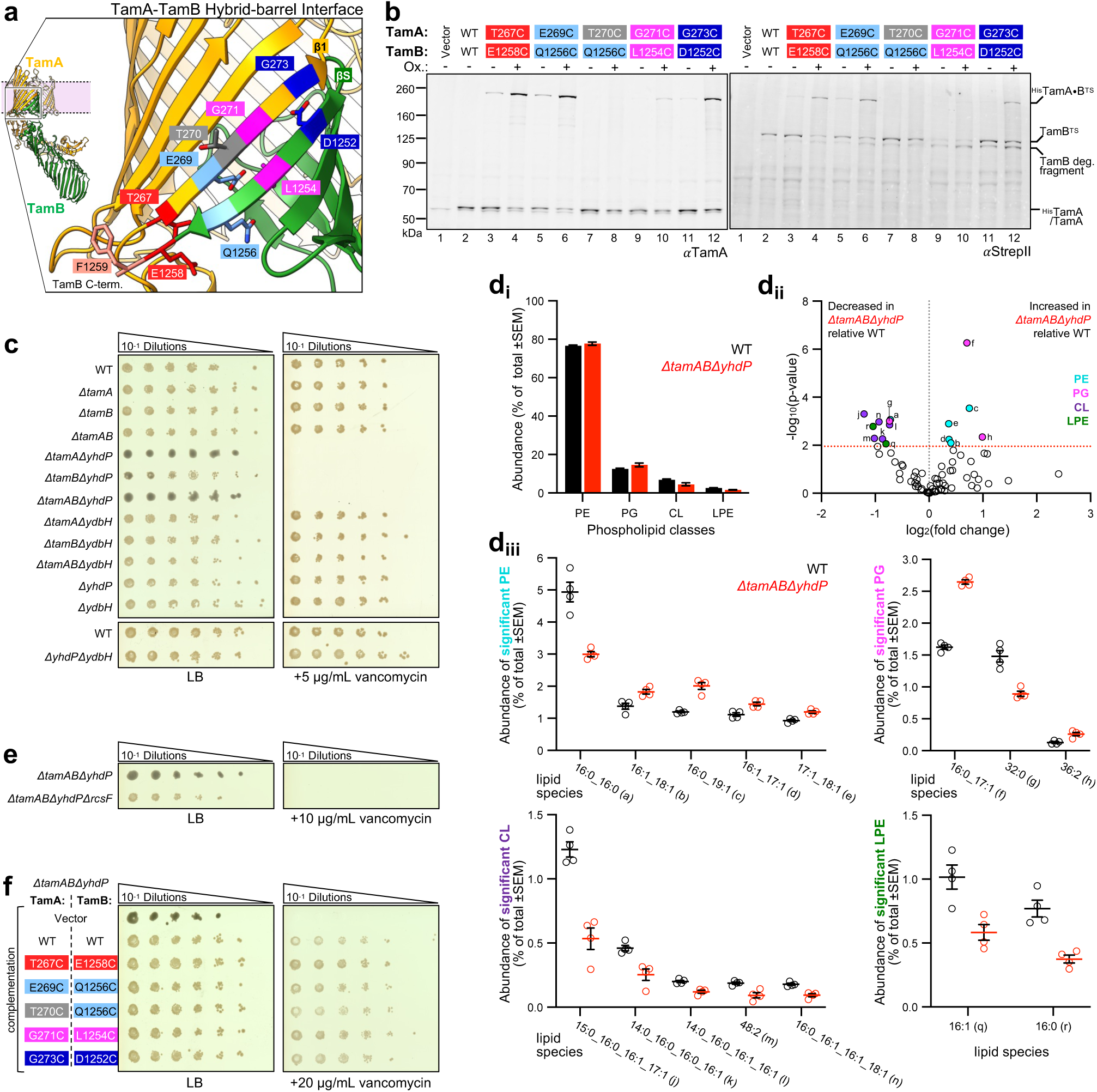
A stable TamA-TamB hybrid-barrel suppresses lipid-transport defects. (**a**) Magnified ^His^TamAB490^TS^-nanodisc structure showing hybridization interface between TamA β-strand 1 (β1) and TamB DUF490 C-terminal β-signal (βS) strand. Register-paired lumen-facing residues that are substituted for cysteine in experiments in **b** and **f** are indicated by matching colors, except for T270 (grey) which is membrane-facing. (**b**) *E. coli* expressing wild-type (WT) ^His^TamAB^TS^, or derivatives with cysteine-pair substitutions at the indicated residues, were mock treated (Ox -) or treated with 200 µM 4-DPS (Ox +), n = 2. Empty pTrc99a as vector control. Intermolecular disulfide-bonds (•) in total cell protein extracts were detected by double-immunoblotting with antibodies against TamA (αTamA) and the TS-tag in TamB (αStrepII). See Fig. S10 and S11 for reduced sample and single-cysteine substitution controls, respectively. (**c**) Efficiency of plating assay. Serial dilutions of WT or mutant derivatives of *E. coli* K-12 W3110 were spotted onto plain LB agar plates or plates containing 5 µg/mL vancomycin, n = 3. See Fig. S12 for results with 10 µg/mL vancomycin or with 0.2% deoxycholate. (**d**) Phospholipidomic analysis of W3110 or W3110Δ*tamAB*Δ*yhdP* outer membranes, (n = 4). **d_i_**, Abundance of phospholipid classes phosphatidylethanolamine (PE), phosphatidylglycerol (PG), cardiolipin (CL), and lysophosphatidylethanolamine (LPE) possessing species with significant differences as in **d_ii_**. **d_ii_**, Volcano plot showing significant fold-changes in abundance of specific phospholipid species between strains (above the red dashed line is significant with false discovery rate correction). Letters correspond to the same species in **d_ii_** and Fig 5c. **d_ii_**, Abundances of all significantly different phospholipid species. See Table S3 for all identified lipid species (**e**) Experiment as in **c** except with W3110 Δ*tamAB*Δ*yhdP* or Δ*tamAB*Δ*yhdP*Δ*rcsF* strains spotted onto plain LB agar plates or plates containing 10 µg/mL vancomycin, n = 3. (**f**) Experiment as in **c** except with W3110 Δ*tamAB*Δ*yhdP* complemented with empty pTrc99a or harboring genes for expression of ^His^TamAB^TS^, or derivatives with cysteine-pair substitutions at positions as in **a**, and spotted onto plates containing 50 µM 4-DPS and the absence or presence of 20 µg/mL vancomycin, n = 3. See Fig. S16 for different treatment concentrations of vancomycin, or 0.2% deoxycholate, and no-4-DPS controls.

To further investigate the mechanisms by which TAM functions in OM homeostasis, we next created a series of *E. coli* strains in which the genes for the major AsmA-like family members were singly or doubly deleted as in previous reports^10,13^. However, to characterize the role of all TAM subunits in phospholipid transport we also created strains in which *tamA* was deleted in combination with the AsmA-like null mutations. Strikingly, we observed that *ΔtamAΔyhdP* and *ΔtamABΔyhdP* strains possessed extreme mucoid phenotypes and fused colonies upon growth on solid media (**Fig. 2c, S12**). The *ΔtamBΔyhdP* strain also possessed a mild mucoid phenotype (**Fig. 2c, S12**) as observed previously in a study in which the phenotype was linked to RcsF, an OM lipoprotein that senses OM defects to trigger the Rcs-stress response pathway and upregulates colanic acid surface capsule production^13^. We therefore constructed a *ΔtamABΔyhdPΔrcsF* quadruple mutant that grew as non-mucoid colonies and strongly suggested that the extreme mucoidy of its parent strain is the result of RcsF sensing significant OM defects (**Fig. 2e**). In line with this model, exposure of the *ΔtamAΔyhdP*, *ΔtamBΔyhdP*, *ΔtamABΔyhdP*, and *ΔtamABΔyhdPΔrcsF* strains to low concentrations of vancomycin, an antibiotic too large to cross the OM of WT *E. coli* (typical MIC > 256 μg/mL), was lethal (**Fig. 2c, 2e, S12**). This indicates that the OMs of these strains are much more permeable. Although the mechanism of OM stress sensing by RcsF remains unknown, one model suggests that it might sense defects in OMP folding by BamA^37,38^. However, we did not observe changes in BamA expression or folding into the OM in any of the knock-out strains (**Fig. S13**). Finally, microscopy has shown that *ΔtamBΔyhdP* strains have altered cell shapes^10,13^. We therefore stained the OM of *ΔtamAΔyhdP*, *ΔtamBΔyhdP*, *ΔtamABΔyhdP*, and *ΔtamABΔyhdPΔrcsF*, imaged the bacteria by fluorescence microscopy, and measured the cell dimensions. All strains were significantly shorter than WT (∼0.4 μm) (**Fig. S14a, b**) which suggested that TAM and/or YhdP assists in a process important for cell elongation during growth and that elongation defects correlate with vancomycin sensitivity more than with mucoidy. The *ΔtamAΔyhdP*, *ΔtamBΔyhdP*, and *ΔtamABΔyhdP* strains were also wider than WT but the *ΔtamABΔyhdPΔrcsF* mutant was not, suggesting that the Rcs stress response has a role in the wide-cell phenotype (**Fig. S14c**).

Our results suggested that TAM (not only TamB) and YhdP have vital functions in the biogenesis of the OM. To determine whether the *ΔtamABΔyhdP* strain possesses an OM with an altered phospholipid content, we isolated the OM from WT *E. coli* or the *ΔtamABΔyhdP* mutant and quantified the relative abundance of phospholipids within each OM by LC-MS/MS (**Fig. 2d, S15, Table S3**). Despite major OM and growth defects, the ratio of the proportions of bulk phospholipid classes in the OM of the *ΔtamABΔyhdP* strain was largely similar to WT (**Fig. 2d_i_**) suggesting that YdbH and/or other transporters can largely maintain this ratio, even if not maintaining the transport rates presumably required for normal elongation. However, significant differences in the abundance of specific phospholipid species in the OM were observed between WT and *ΔtamABΔyhdP* strains (**Fig. 2d_ii_**). The *Δ**tamA**Δ**yhdP*** OM had higher levels of several minor PE and PG species and lower levels of other species including PG(32:0), PE(16:0_16:0), and LysoPE 16:1 and 16:0 (**Fig. 2d_ii_, d_iii_**). More consistent, however, was the reduction in cardiolipin (CL) species in the *ΔtamABΔyhdP* OM including CL(15:0_16:0_16:1_17:1) which is otherwise the most abundant cardiolipin species in the WT OM (**Fig. 2d_ii_, d_iii_, Table S3**).

Finally, having established a correlation between growth defects, vancomycin sensitivity, and changes in phospholipid species in the OM of the *ΔtamABΔyhdP* mutant, we used this strain to investigate the functional relevance of the TAM hybrid-barrel. Complementation of the *ΔtamABΔyhdP* strain with the plasmid for WT ^His^TamAB^TS^ production restored vancomycin resistance and returned colonies to a non-mucoid phenotype (**Fig. 2f, S16**). Transformation of the strain with the empty plasmid vector had no effect on these phenotypes, as expected (**Fig. 2f, S16**). We also quantified the abundances of OM phospholipids of ^His^TamAB^TS^*-*complemented *ΔtamABΔyhdP* and found that it had significantly higher levels of many cardiolipin species in the OM (similar to WT levels) relative to the vector control (*full description in subsection below,* ***Fig. 2d, 4c, Tables S3, S4***). Interestingly, expression of the ^His^TamAB^TS^ hybrid-barrel interface cysteine-pair derivatives in the *ΔtamABΔyhdP* strain also restored non-mucoid growth and vancomycin resistance under oxidizing conditions (**Fig. 2f, S16**). Some even grew better in the presence of vancomycin than cells that expressed WT ^His^TamAB^TS^ (e.g. ^His^TamA_T267C_/TamB_E1258C_^TS^). Together, the results strongly suggest that TAM conducts OM maintenance functions while possessing a stable hybrid-barrel conformation.

### TamB possesses a conserved amphipathic helix that is critical for TAM function

To investigate the function of periplasmic segments of the TAM complex we examined two key interactions between the TamA and TamB subunits. First, our ^His^TamAB490^TS^ structures show that three very highly conserved TamB α-helices (αH1-3) are positioned immediately above the terminus of the lipophilic β-taco in proximity to the membrane interface (**Fig. 3a, S2**). Although TamB αH2 clearly interacts with TamA POTRA3, the positioning of the amphipathic αH3 was ambiguous in our consensus cryo-EM maps (**Fig. 3a, S17**). AF2 also consistently predicts αH3 with high variability and low confidence regardless of the species of origin (**Fig. S3, S19**). For *E. coli* TAM, AF2 predicted αH3 in an alternative position 7-8 Å closer to POTRA2 than in our ^His^TamAB490^TS^ structures (**Fig. 3a**). Notably, αH3 was not observed in the TamAB^His^-amphipol structure^25^. The variable density for αH3 in our cryo-EM maps became more apparent upon conducting variability analysis on our particles which suggested that the helix can reposition within the space between the TamB β-taco terminus and TamA POTRA2 (**Video S4-S6, S13-S15**). The second key inter-subunit interaction is between the coupling-loop that extends from the lipophilic TamB β-taco to bind with POTRA1/2 (**Fig. 3b, S17**). The POTRA2 binding imparts a β-strand secondary structure on the coupling-loop through β-augmentation (**Fig. 3b**). The same β-strand structure is also confidently predicted by AF2 across species and strongly suggests that the TamB-POTRA2 interaction is highly stable (**Fig. S3, S20**).

**Figure 3.**
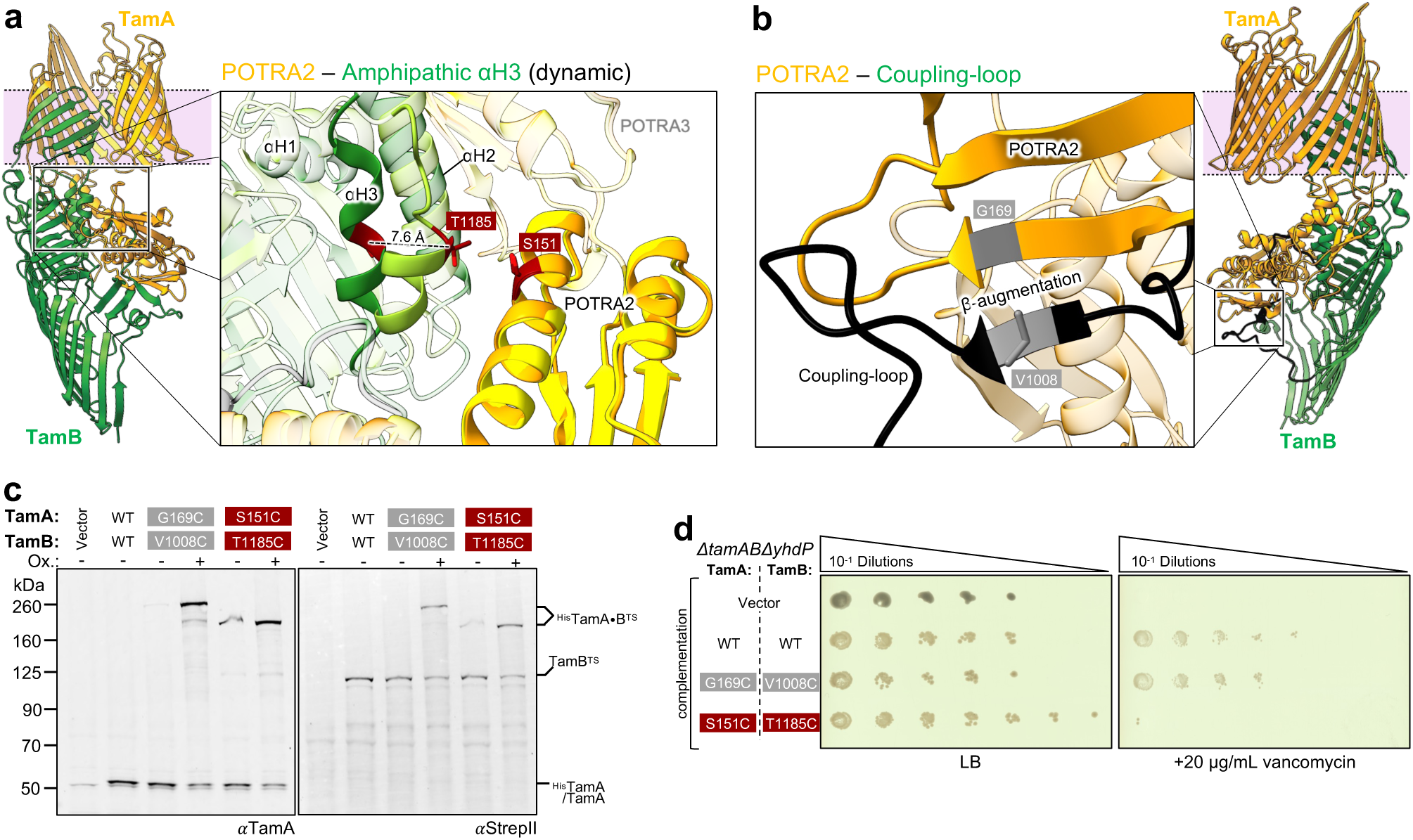
A highly conserved TamB amphipathic α-helix is critical for TAM function. (**a, b**) Magnified views of ^His^TamAB490^TS^-nanodisc structure. **a**, view of TamA POTRA domains and TamB DUF490 conserved α-helices (see Fig. S2 for conservation). The AF2 predicted structure of TamAB from Fig. 1a is also overlaid (TamA, yellow; TamB light-green). Differences in the AF2 prediction, cryo-EM nanodisc structure and 3D-variability analyses (3DVA, see Videos S4-6, 13-15) suggest that αH3 is highly dynamic. **b**, view of TamA POTRA2 interacting with TamB coupling-loop (black) via β-augmentation. Residue pairs substituted for cysteine in experiments in **c** and **d** are indicated by matching colors. (**c**) Experiment conducted as in Fig. 2b except that cysteine-pairs ^His^TamA_G169C_B_V1008C_^TS^ and ^His^TamA_S151C_B_T1185C_^TS^ were tested. See Fig. S10 and S11 for reduced sample and single-cysteine substitution controls, respectively. (**d**) Experiment conducted as in Fig. 2f except that cysteine-pairs ^His^TamA_G169C_B_V1008C_^TS^ and ^His^TamA_S151C_B_T1185C_^TS^ were tested. See Fig. S18 for different treatment concentrations of vancomycin, or 0.2% deoxycholate, and no-4-DPS controls.

To determine the importance of the periplasmic interactions in the function of TamB DUF490, we immobilized either αH3 or the coupling-loop to POTRA2 by expression of cysteine-pairs ^His^TamA_S151C_/TamB_T1185C_^TS^ or ^His^TamA_G169C_/TamB_V1008C_^TS^, respectively (**Fig. 3a, 3b**) and monitored the formation of disulfide-bonds between these positions. Consistent with our structural analyses, we observed very high levels of disulfide-bond formation within both tested interfaces (**Fig. 3c**) which strongly suggests that these structures form *in vivo*. As expected, we did not observe any high molecular weight bands upon immunoblotting reduced samples, nor samples from assays repeated on single-cysteine controls (**Fig. S10, S11**). We next grew derivatives of the *ΔtamABΔyhdP* strain expressing the cysteine-pairs to lock the periplasmic interfaces under oxidizing conditions. The coupling-loop pair ^His^TamA_G169C_/TamB_V1008C_^TS^ restored both normal colony growth and vancomycin resistance to *ΔtamABΔyhdP* (**Fig. 3d, S18**). However, the αH3 immobilization pair ^His^TamA_S151C_/TamB_T1185C_^TS^ did not restore vancomycin resistance which suggests that the bacteria retained significant OM defects (**Fig. 3d, S18**). Together the results strongly suggest that the TamB coupling-loop functions as a stable binder of TamA POTRA2 whereas TAM is unable to function to maintain the OM if the TamB αH3 is stabilized in one position.

### The TAM lipophilic channel transports phospholipids to the bacterial OM

Finally, we tested whether the β-taco of TamB DUF490 functions to maintain OM phospholipid homeostasis. Inspection of the ^His^TamAB490^TS^ cryo-EM maps revealed the presence of additional densities within the β-taco that cannot be attributed to either TamA or TamB subunits (**Fig. 4a, S21**). Striking lipid-like densities that contained projections into the TamB β-taco cavity reminiscent of lipid acyl chains, and terminated at the amphipathic αH3, were observed in the map of ^His^TamAB490^TS^ in nanodiscs reconstituted with *E. coli* phospholipids (**Fig. 4a, S21**). In the detergent ^His^TamAB490^TS^ map amorphous micelle-like density was also present within the β-taco and terminated at αH3 (**Fig. S21**). In both maps the densities are continuously variable over the β-taco region of DUF490 that corresponds to a completely hydrophobic and electrostatically neutral channel that narrows to a terminus under αH3 (**Fig. 4a, Video S7-9, S16-18**).

**Figure 4.**
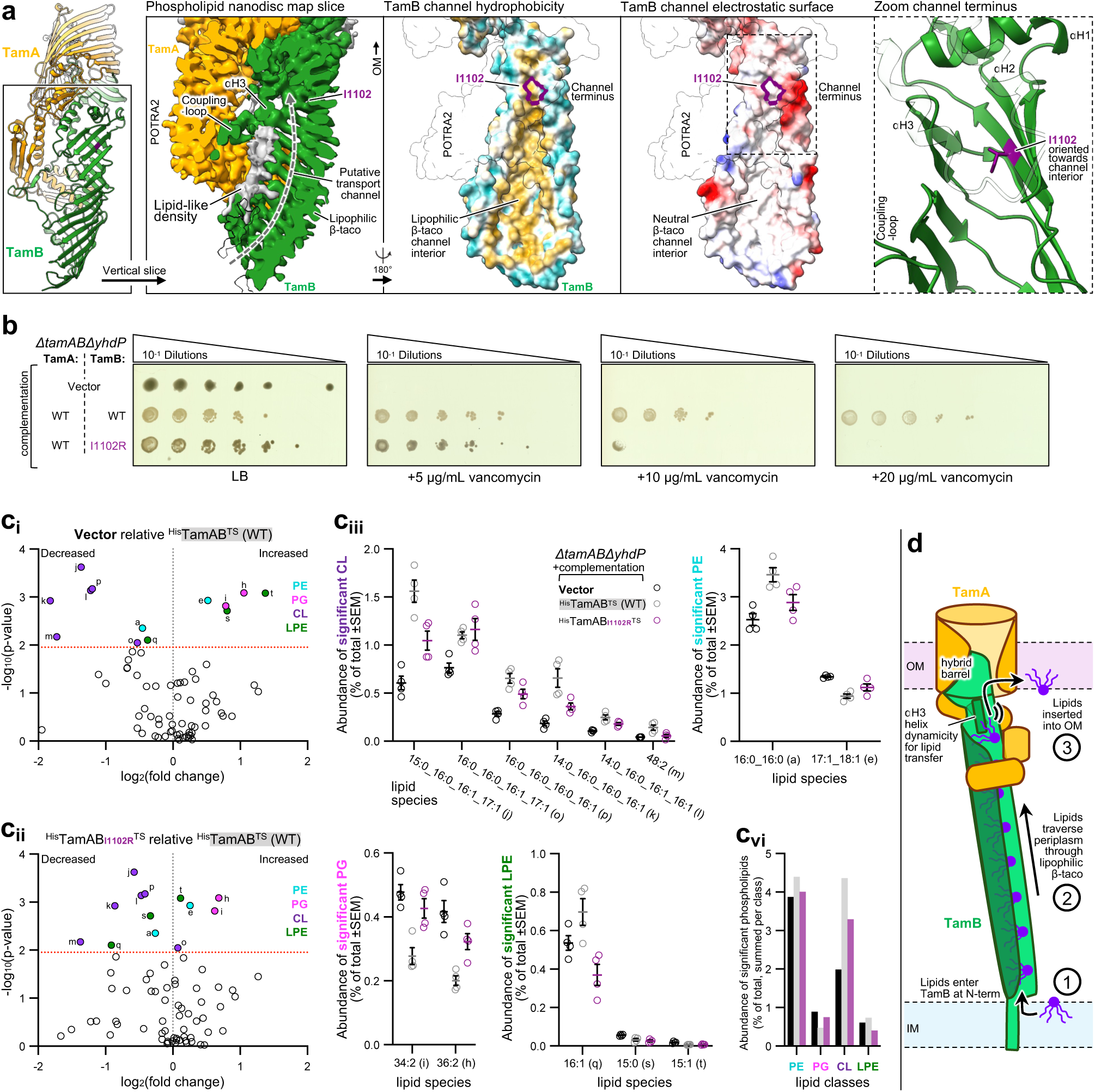
TAM functions as a channel to transport cardiolipin to the bacterial OM. (**a**) Far left, ^His^TamAB490^TS^-nanodisc structure with views aligned to the TamB lipophilic β-taco. Left, ^His^TamAB490^TS^-nanodisc cryo-EM map with vertical slice across the TamB lipophilic β-taco revealing potential lipid densities (grey). Additional obstructing densities are transparent. Map was sharpened with LocScale2. See Fig. S21 and Videos S16-18 for cryo-EM map comparisons of sharpening methods and 3DVA analysis of rivulet of additional dynamic densities in the β-taco. Middle, model showing TamB β -taco surface hydrophobicity (blue, hydrophilic; brown, hydrophobic). TamA is transparent. Right, model showing TamB β -taco surface electrostatics (red, negative; blue, positive; white, neutral). Dashed box indicates channel terminus and the position of TamB_I1102_ (purple outline). Far right, magnified view of channel terminus, αH1-3 locations (2 and 3 transparent), and orientation of I1102. (**b**) Efficiency of plating experiment as in Fig. 2f except Δ*tamAB*Δ*yhdP* strain complemented with empty pTrc99a or harboring genes for expression of ^His^TamAB^TS^, or ^His^TamAB_I1102R_^TS^. See Fig. S22 for 0.2% deoxycholate treatment condition. (**c**) Phospholipidomic experiment as in Fig. 2d except that outer membrane phospholipids from plasmid complemented strains in **b** were analyzed, (n = 4). **c_i_**, Volcano plot showing significant fold-changes in abundance of specific phospholipid species in the absence of TamAB (vector) relative to the presence of WT ^His^TamAB^TS^. Letters correspond to the same species in **c_ii_** and Fig 2d. **c_ii_**, as in **c_i_** except comparing ^His^TamAB_I1102R_^TS^ to ^His^TamAB^TS^. **c_iii_**, Abundances of all significantly different phospholipid species. See Table S4 for all identified lipid species. **c_vi_**, Sum of abundances of all significantly different phospholipid species into classes. (**d**) Molecular mechanism of TAM-mediated phospholipid transport to the bacterial OM. Phospholipids enter the N-terminus of the TamB β-taco, move towards the OM through DUF490, and are released into the outer membrane through a reaction that requires the dynamicity of the conserved amphipathic α-helices.

The neutral lipophilic properties of TamB, and the observed densities in our maps, strongly suggested that the function of the β-taco is to act as a channel to transport phospholipids to the OM. To test this hypothesis, we disrupted the presumptive phospholipid transport path by substituting a channel-facing residue near the TamB channel terminus for a positively charged arginine (**Fig. 4a**, I1102R). This substitution does not affect the expression or assembly of TAM *in vivo*^23^ (**Fig. S22a**). The *ΔtamABΔyhdP* OM dysbiosis reporter strain was transformed with plasmids to produce either WT ^His^TamAB^TS^ or ^His^TamAB_I1102R_^TS^, or an empty plasmid vector, and subsequently grown on solid media. Interestingly, ^His^TamAB_I1102R_^TS^ expression resulted in intermediate-mucoid colonies suggesting that the mutant is only partially functional relative to the activity of WT ^His^TamAB^TS^ (**Fig. 4b, S22b**). Indeed, ^His^TamAB_I1102R_^TS^ complementation only conferred vancomycin resistance at very low concentrations (<10 μg/mL) and partial resistance to secondary bile salts (**Fig. 4b, S22b**). Fluorescence microscopy also showed that ^His^TamAB_I1102R_^TS^ complementation only restored the ability of the bacterial cells to elongate to an intermediate level between that of the WT and vector controls (**Fig. S22c**). Finally, to understand the effect of ^His^TamAB^TS^, ^His^TamAB_I1102R_^TS^, or vector complementation on the phospholipid content of the *ΔtamABΔyhdP* OM, we isolated the OM of these derivatives and subjected them to phospholipidomic analysis (**Fig. 4c, S16**). Remarkably, an array of the otherwise most abundant cardiolipin species were significantly decreased in the vector control relative to complementation with WT ^His^TamAB^TS^ (by ∼3-4-fold depending on species) (**Fig. 4c_i_**, purple symbols, **Table S4**). Furthermore, the same cardiolipin species were reduced in the OM upon ^His^TamAB_I1102R_^TS^-complementation, but to an intermediate magnitude (**Fig. 4c_ii_-c_vi_, Table S4**). There were also variable changes in the levels of minor PE and PG species and rare lysoPE species in the OM of cells transformed with the vector or ^His^TamAB_I1102R_^TS^ plasmid relative to those transformed with the WT ^His^TamAB^TS^ plasmid (**Fig. 4c, Table S4**). The results suggest that the changes in the OM and cell elongation are a direct effect of the activity of the TamB lipophilic β-taco. Given that WT OM levels of cardiolipin are around 6-7% of total phophospholipids under our experimental conditions (**Fig. 2d_i_**), there was a clear disproportionate effect on the recovery of the general abundance of cardiolipin in the *ΔtamABΔyhdP* OM upon ^His^TamAB^TS^ complementation (or partial recovery for ^His^TamAB_I1102R_^TS^) (**Fig. 4c_vi_**). Together these results show that the expression of TAM is necessary to maintain levels of cardiolipin in the OM. Furthermore, the data strongly suggest that the function of the TamB lipophilic β-taco is to transport phospholipids to the OM.

## DISCUSSION

Although data amassed over the last 20 years show the critical importance of TAM for bacterial OM maintenance and virulence^10,11,13,24,31,32,34,39^, how the complex interacts with the membrane, the mechanism of nascent substrate translocation, and the identity of the substrates themselves, has remained poorly understood. In this work, we combine structural, biochemical, genetic, and lipidomic evidence to address each of these critical aspects of TAM function. With respect to how TAM interacts with the OM, our structural data show that the C-terminus of TamB DUF490 embeds into the membrane to form a hybrid-barrel with the β-barrel domain of TamA. Our disulfide-formation assays also show that the hybrid-barrel conformation is stable *in vivo* and suggest that it is the major functional state of the TAM complex in the OM. We therefore propose that a major function of the DUF490 is to act as an Omp85-docking domain as a mechanism to anchor a BLT protein to an OM. Indeed, whether we isolated TAM in detergent or reconstituted it into membrane nanodiscs, we only observed particles in a single hybrid-barrel conformation during cryo-EM. However, our data significantly conflicts with a recent report in which amphipol-solvated TAM particles were observed in either a non-hybrid TamA-closed conformation (C-terminus of TamB disengaged), or a compressed hybrid-barrel conformation^25^. Although differences in construct design (e.g. location of affinity tags) could account for the differential hybrid-barrel stability between the two cryo-EM studies, we speculate that the replacement of the detergent micelle surrounding TAM with either amphipol polymer^25^ or a membrane nanodisc composed of *E. coli* phospholipids differentially retains the native hybrid-barrel conformation. This notion is consistent with the observation that TamB does not co-purify with TamA when the β-barrel domain is tethered closed^40^, indicating that the TAM-amphipol non-hybrid conformation was adopted during a later stage of the reconstitution process. Furthermore, the conformation of the TamA β-barrel in our structures is strikingly similar to the “outward-open” BamA-substrate hybrid-barrels that are formed as transient intermediate states of OMP β-barrel folding reactions^29,30^ which strongly suggests that this is a native conformation widely conserved between these two Omp85 families. On this point, further investigation will be required regarding the involvement of TAM in the assembly of OMPs because it is unlikely that the TAM hybrid-barrel can participate in β-barrel substrate folding reactions in a similar way to BamA when the postulated active site in TamA is otherwise occluded by TamB.

Our data allow us to propose a mechanistic model for phospholipid transport by TAM that is consistent with recent reports on prokaryotic and eukaryotic BLT proteins. First, phospholipids enter the N-terminus of the TamB lipophilic β-taco at the inner membrane (**Fig. 4d**, step 1). This step is supported by recent molecular dynamics simulations on TamB, AsmA-like homologs YhdP and YdbH, and the yeast homolog Cfs1 which each possess a structurally homologous N-terminal chorein domain. In these simulations phospholipids have been observed to spontaneously desorb from the membrane bilayer and enter the β-taco channel to form a lipid rivulet^1,25,41^. The lipid-like density observed in our maps suggest that a rivulet of phospholipids diffuse towards the OM through the TamB β-taco channel at least as far as αH3 (**Fig. 4d**, step 2). Recent cryo-EM structures also observed similar lipid densities along the length of the structurally homologous β-taco domains of both human VPS13A and the *C. elegans* Cfs1 homolog LPD-3^2,42^. Finally, we propose that the conserved amphipathic helices of TamB DUF490 are involved in a critical step in which phospholipids are transferred to the OM (**Fig. 4d**, step 3). In this model the function of DUF490 is to anchor to the OM to act as a dynamic molecular valve whereby αH3 desorbs lipids from the β-taco to translocate them to the inner leaflet of the OM. Not only is such a scenario consistent with the OM permeability defects that we observed upon disulfide tethering αH3, but it is also in line with the defects reported for deletions of the analogous C-terminal helices of YhdP^12^, and with simulations on VPS13A that suggested that the C-terminal amphipathic helices reposition to facilitate unidirectional translocation of lipids to a membrane bilayer^42^. For this step it is plausible that the role of local membrane thinning imparted by TamA^43^ is coordinated with the hybrid-barrel rocking motion observed in our structures to accelerate lipid translocation in an otherwise energetically austere environment at the OM. TAM might therefore function in an analogous way to VPS13A which complexes with the scramblase XKR1, a helical-bundle membrane protein that is proposed to thin the membrane near the lipid entry site^42^.

Our data also show that TAM is required to maintain the level of cardiolipin in the OM. Reduction in cardiolipin species in the absence of TAM correlated with cell elongation and antibiotic permeability defects, and all these phenotypes were restored upon complementation with TAM. TAM can likely transport all phospholipid classes given the viability of *ΔyhdPΔydbH* strains and the finding that multiple lipid substrates bind TamB in both simulations and *in vitro* lipid binding assays^10,13,25,41^. However, our examination of the TamB_I1102R_ mutant in which a positive charge disrupts the lipophilicity of the β-taco strongly suggests that TAM has undergone a specific adaptation to bias the transport of cardiolipin *in vivo*. Although further investigation will be required, these data might unexpectedly link BAM activity in OMP folding with phospholipid transport by TAM. It is known that BAM is tightly bound by cardiolipin, and we have previously observed that BAM activity is affected by changes in OM tension which may be affected by the relative proportion of cardiolipin^29,44–47^. It is therefore possible that the defective folding and stability observed for a subset of OMPs upon deletion of *tam* genes or expression of TamB_I1102R_ (such as the fimbrial-ushers^23,32^), are related to the changes in the phospholipid content of the OM rather than a direct action on the OMP by TAM. Finally, our data are in remarkable alignment with recent live-cell microscopy analyses that show that TamB strictly assembles at cell poles immediately after cell division^48^. Given that the conical shape of cardiolipins cause their passive accumulation to negatively curved cell poles^49,50^, it is therefore tempting to speculate that the negative curvature of TamB and the positioning of the protein at poles are mechanisms to establish the substrate bias of TAM and thereby allow the complex to maintain the OM during a critical stage of bacterial cell growth.

## METHODS

### Bacterial strains and growth conditions

All strains used in this work are listed in Table S5. *E. coli* K-12 NEB5α, XL1-Blue, or XL10-Gold were routinely used for plasmid cloning and mutagenesis. *E. coli* K-12 W3110 and chromosomal gene deletion mutant derivatives, or *E. coli* B-strains BL21(DE3) or BL21(DE3)Δ*dsbA*, were used for all experiments as indicated. Chromosomal gene deletion mutants were constructed by generalized transduction from the *E. coli* Keio collection^51^, FRT-flippase recombination to remove antibiotic cassettes^52^, and confirmed by sequencing. All Strains were cultured in Lysogeny broth (LB) Lennox formulation at 37 ℃ with orbital shaking unless otherwise indicated. Culture media were supplemented with ampicillin (100 ug/mL, 50 ug/mL) as required and or other additives as indicated.

### Plasmid construction and substitution mutagenesis

Plasmids, dsDNA, and ssDNA oligonucleotides used in this study are listed in Tables S6, S7, and S8, respectively. To construct a plasmid for expression of TAM and BAM, pJH114^53^ and pXW47^40^ were amplified with primers 375bamF/375bamR and 377tamF/378tamR, respectively and the resulting fragments assembled by Gibson assembly yielding pMTD1590. To insert a TS-tag into TamB, pMTD1590 was amplified with primers mtd34/35 and the resulting fragment assembled with synthetic dsDNA fragment mtd37 yielding pMTDS80. To construct a plasmid for expression of TAM with a TamB subunit truncated to contain only DUF490, pMTD1590 was amplified with primers mtd33/34 and the resulting fragment assembled with synthetic dsDNA fragment mtd36 yielding pMTDS93. To create amino acid substitution mutations, pMTDS80 was amplified by primer pairs as in Tables S8 and ligated to circularize as appropriate. All plasmids were confirmed by whole-plasmid nanopore sequencing.

### Purification of ^His^TamAB490^TS^ in LMNG detergent

*E. coli* BL21(DE3) harboring pMTDS93 for the expression of ^His^TamAB490^TS^ were inoculated at a starting OD_600_ of 0.05 into 16 L Thomson Ultra Yield flasks, each containing 1 L of LB, and grown at 25 ℃ with shaking at 220 rpm. At an OD_600_ of 0.5 (∼4 h), bacteria were induced with 400 µM IPTG, and grown overnight (∼16 h). Bacteria were pelleted (5,000 x *g*, 15 min, 4 ℃), resuspended in 100 mL ice-cold phosphate buffered saline (PBS; 9 g/L NaCl, 0.144 g/L KH_2_PO_4_, 0.795 g/L Na_2_HPO_4_, pH 7.4), supplemented with SigmaFast EDTA-free protease inhibitor (PI), and frozen in liquid N_2_. Bacteria were thawed on ice, lysed with a Constant Systems Cell Disruptor (30,000 psi, 5 ℃), and the lysate centrifuged (20,000 x *g*, 20 min, 4 ℃) to remove debris. Clarified lysate was then ultracentrifuged (186,000 x *g*, 2 h, 4 ℃) to obtain the membrane pellet. The following steps were conducted with the aid of a Dounce homogenizer: membranes were washed by homogenization into 280 mL ice-cold PBS/PI, pelleted again (186,000 x *g*, 2 h, 4 ℃), homogenized into 140 mL solubilization buffer (50 mM Tris-HCl, 500 mM NaCl, 1% (w/v) n-dodecyl-β-D-maltoside (DDM), pH 8) plus PI, and incubated at 4 ℃ overnight with constant inversion. The solution was ultracentrifuged (186,000 x *g*, 2 h, 4 ℃) and the supernatant was incubated with 12.5 mL StrepTactin XT resin at 4 ℃ overnight with constant inversion. The resin-sample suspension was then transferred to a gravity column (all column steps mentioned hereafter were conducted at 4 °C). The resin was washed with 250 mL TN-LMNG buffer (50 mM Tris-HCl, 500 mM NaCl, 0.003% Lauryl Maltose Neopentyl Glycol (LMNG), pH 8), and protein eluted with 75 mL TN-LMNG-biotin buffer (TN-LMNG also with 50 mM biotin). The eluted protein was then incubated with 2 mL Ni-NTA resin at 4 ℃ overnight with constant inversion. The resin-sample suspension was then transferred to a gravity column, the resin washed with 40 mL TN-LMNG buffer, and protein eluted with 5 mL TN-LMNG-imidazole buffer (TN-LMNG also with 500 mM imidazole). The protein eluate was then desalted with a Sephadex G-25 PD-10 column into TN^low^-LMNG buffer (TN-LMNG but with 150 mM NaCl), concentrated to ∼50 μM using an Amicon Ultra 0.5 mL concentrator (10 kDa cut-off), and frozen in aliquots with liquid N_2_.

### Purification of MSP1D1

*E. coli* BL21(DE3) harboring pMSP1D1^54^ for the expression of MSP1D1 were inoculated at a starting OD_600_ of 0.05 into 1 L of LB in a Thomson Ultra Yield flask and grown at 30 ℃ with shaking (220 rpm). At an OD_600_ of 0.4 bacteria were induced with 1 mM IPTG and grown for a further 3 h. Bacteria were pelleted (5,000 x *g*, 15 min, 4 ℃), resuspended in 20 mL 4x PBS supplemented with 1% Triton X-100 and PI, and frozen in liquid N_2_. Cells were thawed on ice and lysed by sonication using a Sonopuls HD 4050 with a TS-106 probe (Bandelin) at an amplitude of 55% and a pulse time of 8s on/10s off for 30 min. Debris was removed by centrifugation (20,000 x *g*, 20 min, 4 ℃) and the clarified lysate was applied to 10 mL Ni-NTA resin in a gravity column. The resin was washed with 60 mL MSP Wash Buffer A (50 mM Tris-HCl, 300 mM NaCl, 1% Triton X-100, pH 8), followed by 6 CV MSP Wash Buffer B (50 mM Tris-HCl, 300 mM NaCl, 20 mM sodium cholate, 20 mM imidazole, pH 8), 200 mL MSP Wash Buffer C (50 mM Tris-HCl, 300 mM NaCl, 50 mM imidazole, pH 8), and the MSP1D1 eluted with 30 mL MSP Elution Buffer (50 mM Tris-HCl, 300 mM NaCl, 400 mM imidazole, pH 8). An Amicon Ultra 0.5 mL concentrator (10 kDa cut-off) was used to exchange the MSP1D1 into TN^low^ buffer (50 mM Tris-HCl, 150 mM NaCl, pH 8) and for concentration before the protein was frozen in aliquots with liquid N_2_.

### Reconstitution of ^His^TamAB490^TS^ into E. coli phospholipid nanodiscs

A chloroform solution of *E. coli* polar phospholipid extract (Avanti) was dried under a N_2_ stream for 20 min followed by drying in a vacuum desiccator overnight. Dried phospholipids were dissolved in TN^low^ buffer to a final concentration of 30 mM. Phospholipids:MSP1D1:^His^TamAB490^TS^ were mixed in a molar ratio of 70:2.8:1, added to 20 mg of Bio-Beads (Bio-Rad), and incubated at 4 ℃ with constant inversion. The Bio-Beads were changed 3 times over a 24 h period, after which the reconstitution mixture was incubated with 125 μL StrepTactin XT resin at 4 ℃ overnight with constant inversion. The resin-sample suspension was then transferred to a gravity column (all column steps mentioned hereafter were conducted at 4 °C). The resin was washed with 2.5 mL TN^low^ buffer and then the ^His^TamAB490^TS^-nanodiscs were released from the resin by adding 0.75 mL TN^low^-biotin buffer (50 mM Tris, 150 mM NaCl, 50 mM biotin, pH 8) and incubating at 4 ℃ overnight with constant inversion. The eluted protein was isolated by pipetting off the supernatant after the resin was gravity settled. ^His^TamAB490^TS^-nanodiscs were concentrated to ∼12.4 μM using an Amicon Ultra 0.5 mL concentrator (10 kDa cut-off), and frozen in aliquots with liquid N_2_.

### Cryo-EM sample preparation and imaging

All grids were prepared with an FEI Vitrobot Mark IV at 4 ℃ and 100 % humidity. 3 µL of ^His^TamAB490^TS^ LMNG (diluted to 27 µM), or undiluted ^His^TamAB490^TS^-nanodiscs, was applied to glow-discharged UltrAuFoil R1.2/1.3 300 mesh grids with a wait time of 60 s. Grids were blotted for 3 s with a blot force of 9. Datasets were recorded using a Titan Krios operated at 300 kV fitted with a Gatan BioQuantum 15 e^-^V slit energy filter and K3 camera. Movies were collected at a magnification of 105,000 and a pixel size of 0.83 Å. The ^His^TamAB490^TS^ LMNG dataset consisted of 10,106 movies and was collected at a dose rate of 9.7 e^-^/A^2^/s with an exposure time of 6 s, for a total dose of 58 e^-^/Å and 80 total frames. The ^His^TamAB490^TS^-nanodiscs dataset consisted of 20,059 movies and was collected at a dose rate of 12.0 e^-^/Å/s with an exposure time of 6 s, for a total dose of 72 e^-^/Å^2^ and 75 total frames.

### Cryo-EM image processing for ^His^TamAB490^TS^ structure in LMNG

All processing was performed with CryoSPARC^55^ v4.7.0 or v4.7.1. The following workflow is also described in Figure S4. Movies were aligned using patch motion correction, followed by patch CTF estimation^56^. Exposures were accepted using strict cut-offs of a CTF fit under 4 Å and a relative ice thickness under 1.09 leaving 3,227 exposures. Particles were picked initially using the blob picker tool and extracted at a box size of 384 pixels down sampled to 192 pixels. Several rounds of 2D classification provided distinct classes used for subsequent template picking. Exposures were denoised and both blob and template picking were run again on all denoised exposures. Because the two picking methods offered some distinct picks, these were combined yielding 3,448,506 particles after removal of duplicates. Particles were extracted at 384 pixels and binned to 128 pixels for two rounds of multi-class *ab-initio* reconstruction until a low-resolution volume consistent with the TAM complex was generated. Several ‘decoy’ volumes were generated then iterative rounds of particle sorting by heterogeneous refinement, followed by followed by using the TAM-like and decoy volumes. The particles were re-extracted to 384 pixels and a high-resolution *ab-initio* volume was generated which was then used for a non-uniform refinement^57^ to produce a consensus structure of 3.80 Å. A mask was made to ensure coverage of ^His^TamAB490^TS^ and exclude the detergent micelle which was then used for local refinement. The final volume consisted of 80,583 particles and achieved a resolution of 3.51 Å. Map sharpening was with CryoSPARC, EMReady2^58^, or Locscale2^59,60^.

### Cryo-EM image processing for ^His^TamAB490^TS^ structure in phospholipid nanodiscs

The processing for ^His^TamAB490^TS^ in phospholipid nanodiscs was similar to that in LMNG detergent, differences are highlighted below (Figure S5). After patch motion correction and CTF estimation, exposures were accepted using strict cut-offs of a CTF fit under 5 Å and a relative ice thickness under 1.07 leaving 6,290 exposures. Particles were picked initially using the blob picker tool and extracted at a box size of 384 pixels down sampled to 192 pixels. Several rounds of 2D classification provided distinct classes used for subsequent template picking. Exposures were denoised and both blob and template picking were run again on all denoised exposures. Several attempts were made to improve Topaz^61^ training for particle picking; however, the overall result was sub-optimal for the entire dataset. It was again noted that all three picking methods, blob, template, and Topaz offered some distinct picks, and so these were combined yielding 6,730,439 particles after duplicate removal. Particles were extracted at 384 pixels and binned to 128 pixels for two rounds of multi-class *ab-initio* reconstruction until a low-resolution volume consistent with the TAM complex was generated. Again, several decoy volumes were generated then three iterative rounds of particle sorting by heterogeneous refinement followed using the TAM-like and decoy volumes. The particles were re-extracted to 384 pixels following another three rounds of iterative heterogeneous refinement, after which an initial consensus map of 5.03 Å was generated by Non-Uniform refinement on 305,278 particles. Following a final round of duplicate removal and another three rounds of iterative heterogeneous refinement, a two-class high-resolution *ab-initio* volume was produced along with a mask surrounding ^His^TamAB490^TS^ excluding the nanodisc and other low-resolution features near where the TamB β-taco was truncated that obstructed alignments. Local refinement was then conducted to reach a final consensus reconstruction of 3.71 Å from 103,937 particles. The consensus volume and corresponding particles were used for 3D Variability Analysis^62^ along three component axes using a filter resolution of 6 Å. The nanodisc density from the consensus particle stack was subtracted and used to construct a 3D flex mesh consisting of two segments corresponding to the periplasmic and membrane-embedded portions of the complex (Figure S23). The 3D flex mesh was used to train a 3D Flex model for the generation of two 3D flex volume series^63^.

### Model building and refinement

The AlphaFold-predicted model for full-length TamAB was loaded into UCSF ChimeraX 1.10.1 and residues 1-836 of TamB, excluding DUF490, were deleted. A model of TamAB490 was generated using Boltz-2^64^ and TamB residues 1099-1259 of the model used to replace the corresponding residues in the truncated AlphaFold model. The model was docked into the ^His^TamAB490^TS^-nanodisc map in ISOLDE 1.7^65^ wherein bulk flexible fitting at a temperature of 20 K. The temperature of the simulation was reduced to 0 K for final correction of Ramachandran outliers, sidechain outliers, and clashes. The ISOLDE restraints for the ^His^TamAB490^TS^-nanodisc model were then used for real-space refinement in Phenix^66,67^. The ^His^TamAB490^TS^-nanodisc model was docked and refined to the ^His^TamAB490^TS^ LMNG cryo-EM map to produce the resulting model in detergent. Following refinement, both models were validated using Molprobity^68^ in Phenix. Model statistics are in Table S10.

### Western immunoblotting

The iBlotII transfer device (Life Technologies) was routinely used to transfer protein gels to nitrocellulose membranes. Immunoblotting buffer (Odyssey Blocking Buffer (Li-Cor)) supplemented with 0.01% Tween-20 was used for blocking steps and as a diluent for primary and secondary antibodies (antibodies see Table S9). Membranes were blocked for 1 hr, incubated with primary antibodies overnight [αStrepII (1:5,000 dilution), and/or αTamA, αTamB, or αBamA (1:10,000 dilution)], incubated with secondary antibodies [goat α-rabbit (Licor 926-68021) and goat α-mouse (Licor 926-32210) at 1:5,000 dilution] for 4 hr, washed (3 x 5 min with PBS containing 0.01% Tween-20 then 2 x 5 min PBS) and air dried (20 min, 37 °C). The membranes were then scanned using an Odyssey DLx infrared imager. Pixel intensities of detected proteins were measured using Fiji software.

### In vivo disulfide-bond formation assay

To observe site-specific interactions between TamA and TamB *in vivo*, disulfide-bond formation assays were conducted essentially as previously described^29,69^. *E. coli* BL21(DE3)*ΔdsbA* containing plasmids encoding TamAB subunits with cysteine substitutions at positions of interest were grown overnight from a single colony in 10 mL LB with shaking (25 °C, 250 rpm). Cultures were pelleted (2,918 x *g*, 10 min, 4 °C), washed with 10 mL LB, and resuspended in 10 mL LB before inoculating fresh 10 mL LB subcultures at OD_600_ = 0.05. After cultures were grown for 4 h (25 °C, 250 rpm) to OD_600_ ∼0.4-0.6, 0.4 mM IPTG was added to induce expression of TAM for 1 h. 1 mL samples of induced subculture were aliquoted into tubes on ice, pelleted (2 min, 4 °C, 20,817 x *g*), resuspended in 1 mL ice-cold PBS, and then bacteria were incubated with 0.2 mM of the thiol specific oxidiser 4,4’-dipyridyl disulfide (4-DPS). Mock treatment controls were instead incubated with the equivalent volume of ethanol. All samples were incubated on ice for 30 min, pelleted (10,000 x *g*, 2 min, 4 °C) and resuspended in ice-cold PBS (0.5 mL). Bacteria were lysed and proteins precipitated by adding trichloroacetic acid (TCA) and phenylmethanesulfonyl fluoride (PMSF) (final concentration of 10% v/v TCA and 4 mM PMSF) and incubated on ice for 10 min. The precipitated proteins were pelleted (20,817 x *g*, 10 min, 4 °C), washed with ice-cold acetone (0.6 mL), re-pelleted, and air dried (37 °C, 20 min). Total protein samples were resuspended in 2x SDS Protein Loading Dye (100 mM Tris-HCl pH 8, 20% (v/v) glycerol, 0.2% (w/v) bromophenol blue, 4% (v/v) SDS) in a volume normalized to an OD600 measurement recorded immediately as induced subculture samples were taken (volume in μL = 200 x OD_600_). Samples were heated to 99°C for 15 min and aliquots resolved by SDS-PAGE on 8%-16% Tris-glycine minigels (Invitrogen) (150 V, 1 hr 40 min, room temperature) before being transferred to nitrocellulose for Western immunoblot analysis as above. The immunoblot was either singly or doubly probed with appropriate antisera (Table S9).

### Heat-modifiable electrophoretic mobility shift assay

To test for BamA assembly/stability, mobility shift assays were conducted essentially as previously described^29,69^. Bacteria were grown as described in the *in vivo* disulfide-formation assay and then bacterial pellets from 1 mL aliquots were resuspended (volume, µL = OD_600_ x 100) in BugBuster Master Mix containing PI and lysed on ice for 5 min. 8 μL aliquots of lysates were mixed with 5 μL 2x SDS protein gel loading dye and were then either retained on ice or heated to 99°C for 10 min. Proteins were then resolved by cold SDS-PAGE (gel tank embedded in packed ice) and transferred to nitrocellulose for immunoblotting.

### Bacterial dilution spot plating

Single colonies of strains of interest were used to inoculate of 10 mL LB cultures which were grown overnight (37 °C, 250 rpm). Bacterial cultures were diluted to OD_600_ of 1 with LB and then serial 10-fold dilutions were prepared within a 96-well tray. A 96 pin 1 µL replicator was then used to spot bacterial dilutions onto square plates containing LB agar supplemented with treatments of interest. For assays conducted on complemented strains, IPTG was not included in the media unless otherwise indicated. Plates were incubated (37 °C, 20 h) and imaged with transillumination using an Epson Perfection V850 Pro.

### Isolation of outer membrane fractions

An overnight culture (200 mL LB) was inoculated with 10 single colonies of the strain of interest and grown overnight (20 hr, 37 °C, 250 rpm). The overnight culture was centrifuged (20,000 x *g*, 20 min, 4 °C) and the bacterial pellet was resuspended in 15 mL ice-cold PBS plus PI. Cell suspensions were subsequently lysed using a Constant Systems cell disruptor (30 kpsi, 5 °C, 3 passes). Lysates were clarified by centrifugation (15,000 x *g*, 15 min, 4 °C) and the resulting supernatants were ultracentrifuged (60,000 x *g*, 40 min, 4 °C) to obtain the whole membrane pellets. Whole membrane pellets were rinsed with 20 mL PBS (keeping pellets intact) before subsequent homogenization into 1 mL TN^low^ buffer using a Dounce homogeniser. Suspensions were prepared to an A280 of 16 and 500 µL was applied to a 30-55% sucrose density gradient. Gradients were centrifuged overnight (173,000 x *g*, 16 hr, 4 °C). 200 µL fractions were collected from immediately below the meniscus of the gradient, and frozen in liquid N_2_.

### Lipid extraction

Lipids were extracted from 50 µL of the purified OM fractions using a two-phase methyl-tert-butyl-ether (MTBE)/methanol/water protocol^70^. Membrane fractions were mixed with 850 µL MTBE and 250 µL methanol containing internal standards: 2 nmoles each of CL 14:0/14:0/14:0/14:0, PE 17:0/17:0, PG 17:0/17:0, PC 19:0/19:0; 1 nmole of PA 17:0/17:0; and 0.5 nmoles each of LPC 17:0, LysoPE 17:1, and LysoPS 17:1. Samples were sonicated for 30 min in a 4 °C water bath, followed by the addition of 162 µL milliQ water to induce phase separation. Samples were vortexed, centrifuged (2,000 x *g*, 5 min), and the upper organic phase was aliquoted to 5 mL glass tubes. The remaining aqueous phase underwent two additional extractions using 500 µL MTBE and 150 µL methanol followed by sonication for 15 min and phase separation with 125 µL water. Organic phases from the three extractions were combined and vacuum-dried overnight in a Savant SC210 SpeedVac (Thermo Fisher Scientific). Dried lipids were dissolved in 100 µL of 25% HPLC grade methanol/25% 1-butanol/50% MilliQ water containing 0.1% formic acid and 10 mM ammonium formate, centrifuged (2,000 x *g*, 20 min), and 90 µL was transferred to glass HPLC vials.

### Liquid chromatography-tandem mass spectrometry

Phospholipidomics data was acquired using a Thermo Fisher Q-Exactive HF-X mass spectrometer coupled to a Vanquish HPLC. Liquid chromatography was performed on a Waters C18 Acquity UPLC column (2.1 × 100 mm, 1.7 μm pore size) using a 25 min binary gradient for the mobile phase at 0.28 mL/min: 0 min, 80:20 A/B; 3 min, 80:20 A/B; 5.5 min, 55:45 A/B; 8 min, 35:65 A/B; 13 min, 15:85 A/B; 14 min, 0:100 A/B; 20 min, 0:100 A/B; 20.2 min: 80/20 A/B; 25 min: 80:20 A/B. Mobile phase A was 10 mM ammonium formate, 0.1% formic acid in acetonitrile:water (60:40); mobile phase B was 10 mM ammonium formate, 0.1% formic acid in isopropanol:acetonitrile (90:10). Data was acquired in full scan/data-dependent MS^2^ mode (resolution 60,000 FWHM, scan range 220–1600 *m/z*, AGC target 3e^6^, maximum integration time 50 ms) in both positive and negative mode for each sample. The injection volume was 5 µL for positive mode, and 10 µL for negative mode. The ten most abundant ions in each cycle were subjected to MS^2^ using resolution 15,000 FWHM, isolation window 1.1 *m/z*, AGC target 1e^5^, collision energy 30 eV, maximum integration time 35 ms, dynamic exclusion window 8 s. An exclusion list of background ions was based on a solvent blank. An inclusion list of the [M + H]^+^, [M + NH_4_]^+^, and [M - H]^−^ ions for all internal standards was used. No shift in lipid retention time was observed throughout the run.

### Phospholipidomic analysis

Phospholipids were annotated using LipidSearch v5.2 beta (Thermo Fisher Scientific), based on accurate precursor (5 ppm mass tolerance) and diagnostic product ions (8 ppm). Levels of phospholipids in each sample were estimated by dividing the peak area by that of the class-specific internal standard and multiplying by the amount of internal standard added. Relative abundance of phospholipids was calculated by dividing the amounts of lipid by the total lipid content estimated for each sample. Statistical analysis of relative lipid abundance was conducted using MetaboAnalyst 6.0^71^. Two sample, independent t-test was used for comparisons between two groups. One-way ANOVA followed by Tukey’s multiple comparisons was used to compare between more than two groups. P-values were corrected for false discovery rate using the Benjamini-Hochberg test. An adjusted p-value ≤ 0.05 was considered statistically significant.

### Outer membrane staining and microscopy of live bacteria

Single colonies of strains of interest were used to inoculate of 10 mL LB cultures which were grown overnight (37 °C, 250 rpm). Strains were subcultured (1:50 dilution) and grown for 3 hr (37 °C, 250 rpm). 250 μL samples of the cultures were aliquoted to 1.5 mL tubes and incubated with 33 μM FM4-64X (37 °C, 15 min, 300 rpm). Bacteria were pipetted onto custom prepared LB 1% agarose pads on glass slides, allowed to adsorb, and subsequently visualized using an Olympus BX51 fluorescence microscope with CellSens Standard imaging software (v4.4). Image processing and quantification of cell lengths was completed using the MicrobeJ plugin^72^ for Fiji ImageJ (v2.16.0).

### Data availability

Structural data supporting findings in this study have been deposited in the Protein Data Bank (PDB) and the Electron Microscopy Data Bank (EMDB). The accession codes of the cryo-EM maps and accompanying atomic models have been provided for: (1) ^His^TamAB490^TS^ in LMNG detergent (EMDB-69220, PDB: 23SQ) (2) ^His^TamAB490^TS^ in lipid nanodiscs (EMD-69219, PDB: 23SP). Phospholipidomic data sets have been deposited in Zenodo with DOI: 10.5281/zenodo.18665426. Any additional information required to reanalyze the data reported in this paper is available from the corresponding authors upon request.

## ACKNOWLEDGEMENTS

AGE and CZ are supported by University of Sydney Research Training Program Scholarships. GSKA is supported by the University of Sydney International Stipend Scholarship and the University of Sydney International Tuition Fee Scholarship. This work was supported by an Australian Research Council (ARC) Discovery Project (DP260100880) awarded to MTD, DLL, RAN, and HDB, and seed funding from both the University of Sydney Infectious Diseases Institute and the University of Sydney Centre for Drug Discovery Innovation to MTD. MTD is an affiliate member, and ABG and SHJB are members, of the ARC Industrial Transformation Training Centre for Cryo-electron Microscopy of Membrane Proteins (IC200100052). RAN is supported by an ARC DECRA (DE250101030). AGS is supported by National Health and Medical Research Council (NHMRC) grants, 2016308. The work was partially supported by NHMRC Ideas Grants 2002660 and 2028164 to ASD. CBB is an associate investigator of the ARC Centre of Excellence for Innovations in Peptide and Protein Science (CiPPS) and ARC Centre of Excellence for Synthetic Biology (CoESB). HDB is supported by the Intramural Research Program of the National Institute for Diabetes and Digestive and Kidney Diseases. We acknowledge the use of the University of Wollongong Cryogenic Electron Microscopy Facility at the Molecular Horizons. We acknowledge helpful discussions with Martina Sanderson-Smith. We thank M. Stephen Trent for providing us with *E. coli* W3110 and some reported *asmA*-family deletion mutants.

## AUTHOR CONTRIBUTIONS

The study was conceived by MTD, ABG, AGE, and LSRA, but all authors contributed to experimental design. Cryo-EM experiments, data processing, and modeling was conducted by LSRA, RAN, and ABG. Lipidomics experiments including sample collection, preparation, and analysis was conducted by AGE, CZ, and ASD. All other experiments were conducted by AGE. GSKA, RAN, DLL, HDB, AGS, CBB contributed significant materials, analysis, and intellectual support to the project. The paper was written and edited by all authors. The project was supervised by LSRA, ABG, and MTD.

