## Supplementary Information for "Mechanism of phospholipid transport to the bacterial outer membrane by TAM"

Alanah Eisenhuth *et al.*

Corresponding authors:




**This PDF file includes:**

Figure S1. Leaky expression of TAM complements but overexpression is toxic. pg 2
Figure S2. Conservation of TamB residues. pg 3
Figure S3. Predicted TAM structures across key Gram-negative species. pg 4
Figure S4. Cryo-EM reconstruction workflow for $^{His}$TamAB490$^{TS}$ in LMNG detergent. pg 5-6
Figure S5. Cryo-EM reconstruction workflow for $^{His}$TamAB490$^{TS}$ phospholipid nanodiscs. pg 7-8
Figure S6. Conservation of TamA residues. pg 9
Figure S7. Conserved intermolecular salt-bridges in the TAM hybrid-barrel. pg 10
Figure S8. The conformation of the $^{His}$TamAB490$^{TS}$-nanodisc structure is similar to the transient hybrid-barrel formed by BamA during OMP assembly. pg 11
Figure S9. Extended comparison of $^{His}$TamAB490$^{TS}$-nanodisc and TamAB$^{His}$-amphipol structures. pg 12
Figure S10. Reduced SDS-PAGE of disulfide crosslinking samples. pg 13
Figure S11. Single cysteine substitution controls for disulfide crosslinking experiments. pg 14
Figure S12. Simultaneous deletion of lipid bridge encoding genes *tamAB* and *yhdP* result in stress responses and outer membrane defects. pg 15
Figure S13. The assembly of BamA is unaffected by deletion of lipid transport genes. pg 16
Figure S14. Analysis of WT and lipid bridge deletion mutants by microscopy. pg 17
Figure S15. Isolation of *E. coli* outer membranes by sucrose density gradient. pg 18
Figure S16. Additional conditions tested for TAM derivatives with disulfide-tethered hybrid-barrel interface. pg 19
Figure S17. Density of important periplasmic segments in $^{His}$TamAB490$^{TS}$-nanodisc cryo-EM map. pg 20
Figure S18. Additional conditions tested for TAM derivatives with disulfide-tethered periplasmic segments. pg 21
Figure S19. Variability in DUF490 helices between TAM predictions. pg 22-23
Figure S20. Prediction of conserved POTRA2-DUF490 coupling-loop interaction. pg 24
Figure S21. Additional densities in the cryo-EM maps of $^{His}$TamAB490$^{TS}$. pg 25
Figure S22. TAM dysfunction due to I1102R substitution in the TamB lipid channel. pg 26
Figure S23. Analysis of the conformational dynamics of TAM subunits. pg 27
Table S1. DUF490-containing proteins.
Table S2. Statistical analyses for all experiments except phospholipidomics. pg 28
Table S3. Abundances of phospholipids relevant to Fig. 2d. pg 28-29
Table S4. Abundances of phospholipids relevant to Fig. 5c. pg 30-31
Table S5. Bacterial strains used in this study. pg 32
Table S6. Plasmids used and constructed in this study. pg 33
Table S7. Synthetic dsDNA fragments used in this study. pg 33
Table S8. ssDNA Oligonucleotides used in this study. pg 34
Table S9. Antisera used in this study. pg 35
Table S10. Model statistics. pg 35
Video Information: Videos S1-20. pg 36
Supplementary References. pg 37

### SUPPLEMENTARY FIGURES

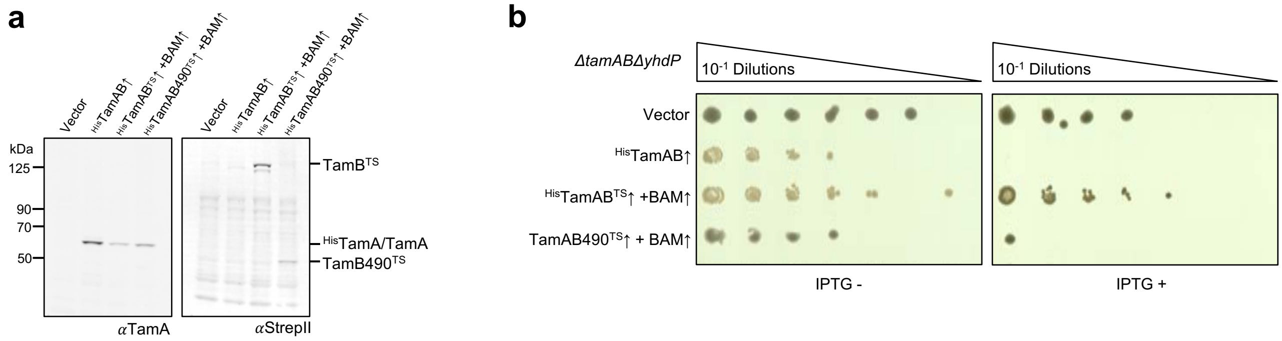

**Figure S1. Leaky expression of TAM complements but overexpression is toxic.**
(**a**) Experiment is the same as in Fig. 1b but uninduced. *E. coli* BL21 cultures expressing $^{His}$TamAB, $^{His}$TamAB$^{TS}$ and BamABCDE, $^{His}$TamAB490$^{TS}$ and BamABCDE, or possessing an empty pTrc99a plasmid were grown without IPTG at 25 °C alongside the induced samples for 1h. Total cell protein was probed by western immunoblotting using αTamA or αStrepII (n = 3). (**b**) Efficiency of plating assay. Serial dilutions of WT or mutant derivatives of *E. coli* K-12 W3110 *ΔtamABΔyhdP* complemented with empty pTrc99a or harboring genes for expression of $^{His}$TamAB, $^{His}$TamAB$^{TS}$ and BamABCDE, or $^{His}$TamAB490$^{TS}$ and BamABCDE were spotted onto plain LB agar plates or plates containing 0.4 mM IPTG, n = 3. Plates were incubated at 37 °C for 20 hr.

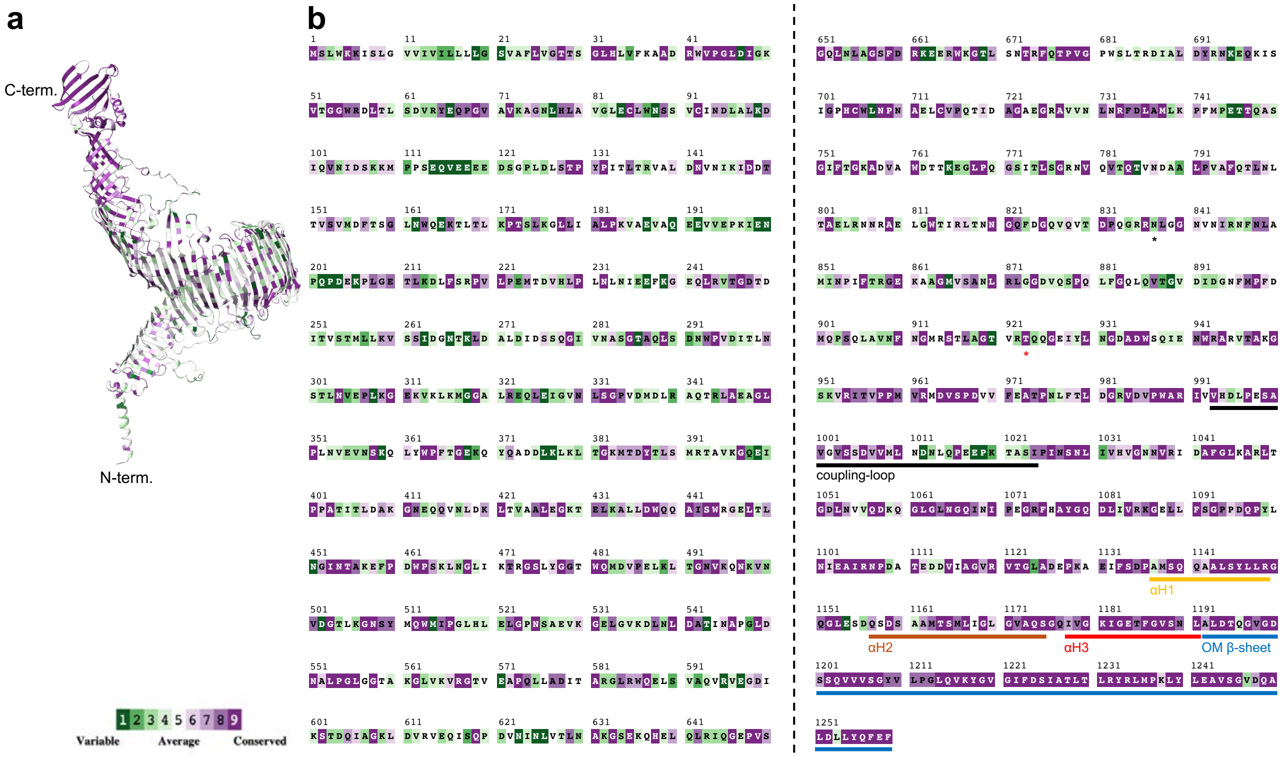

**Figure S2. Conservation of TamB residues.**
The conservation of TamB residues was calculated using ConSurf[1]. The color scale for conservation level was applied to (**a**) the AF2 predicted structure of TamB and (**b**) the primary aa sequence of TamB. Black asterisk denotes N-terminus of residues included in TamB490 construct for structural determination. Red asterisk denotes N-terminus of DUF490 (923-1259). Functional secondary structure regions are denoted based on the $^{His}$TamAB490$^{TS}$-nanodisc structure.

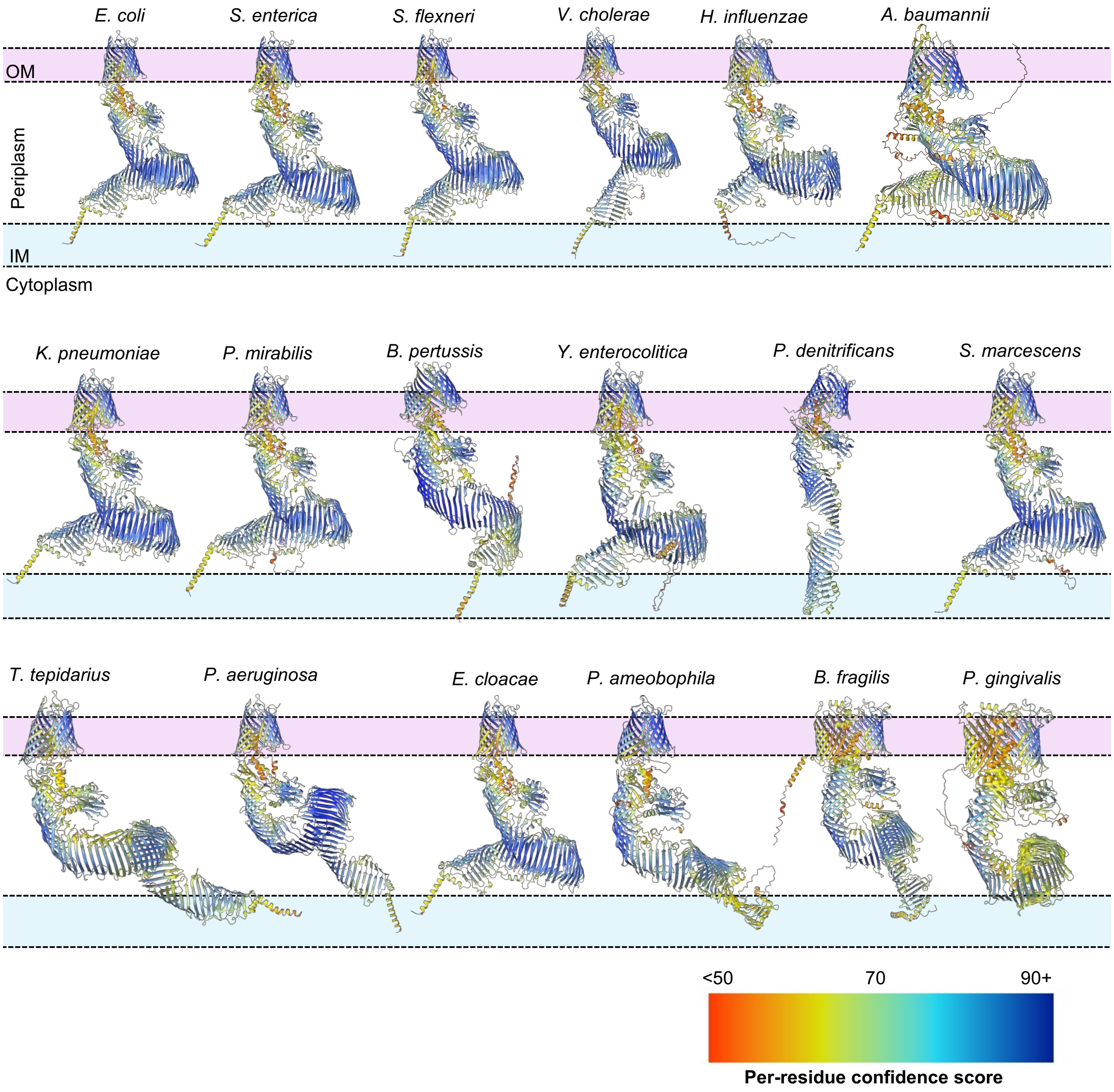

**Figure S3. Confidence of predicted TAM structures across key Gram-negative species.** AlphaFold2 was used to predict the complex structure of TamA and TamB from clinically relevant and diverse Gram-negative bacteria. The DUF490 α-helices are consistently predicted with poor confidence suggestive of intrinsic conformational dynamicity. Rank1 models are shown from five-model prediction runs (see also Fig. S19 and S20).

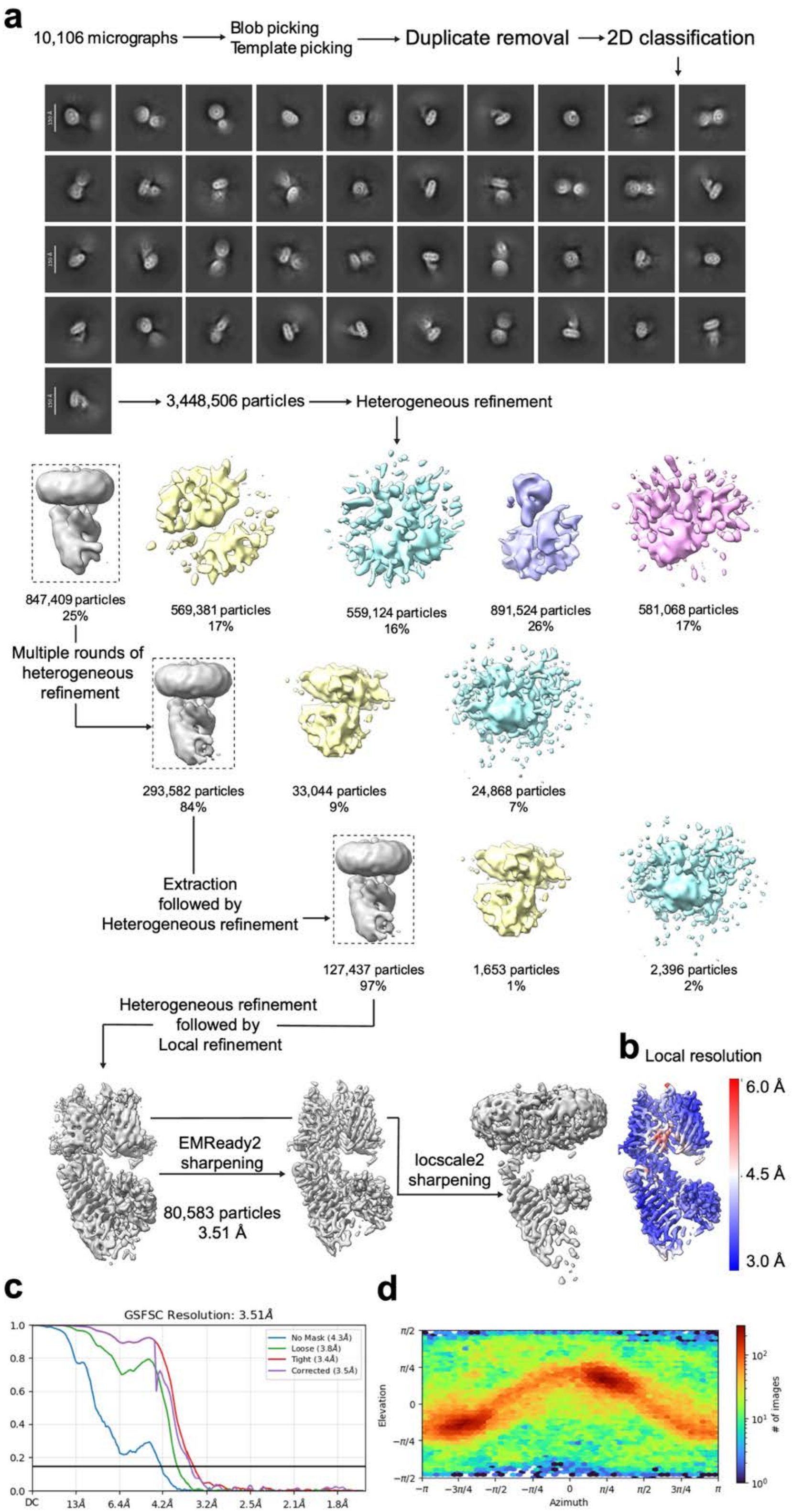

**Figure S4. Cryo-EM reconstruction workflow for $^{His}$TamAB490$^{TS}$ in LMNG detergent.**
(**a**) Workflow for processing of cryo-EM dataset collected on $^{His}$TamAB490$^{TS}$ in LMNG detergent

micelles using CryoSPARC. Classes from intermediate rounds of heterogeneous refinement and the final map from local refinement and sharpening using either EMReady2 or LocScale2. (**b**) Map of $^{His}$TamAB490$^{TS}$ in detergent colored by local resolution (CryoSPARC local resolution estimation). (**c**) Gold-standard Fourier-shell correlation (GSFSC) for $^{His}$TamAB490$^{TS}$ in detergent. (**d**) Orientation distribution for all particles used in the reconstruction of $^{His}$TamAB490$^{TS}$ in detergent.

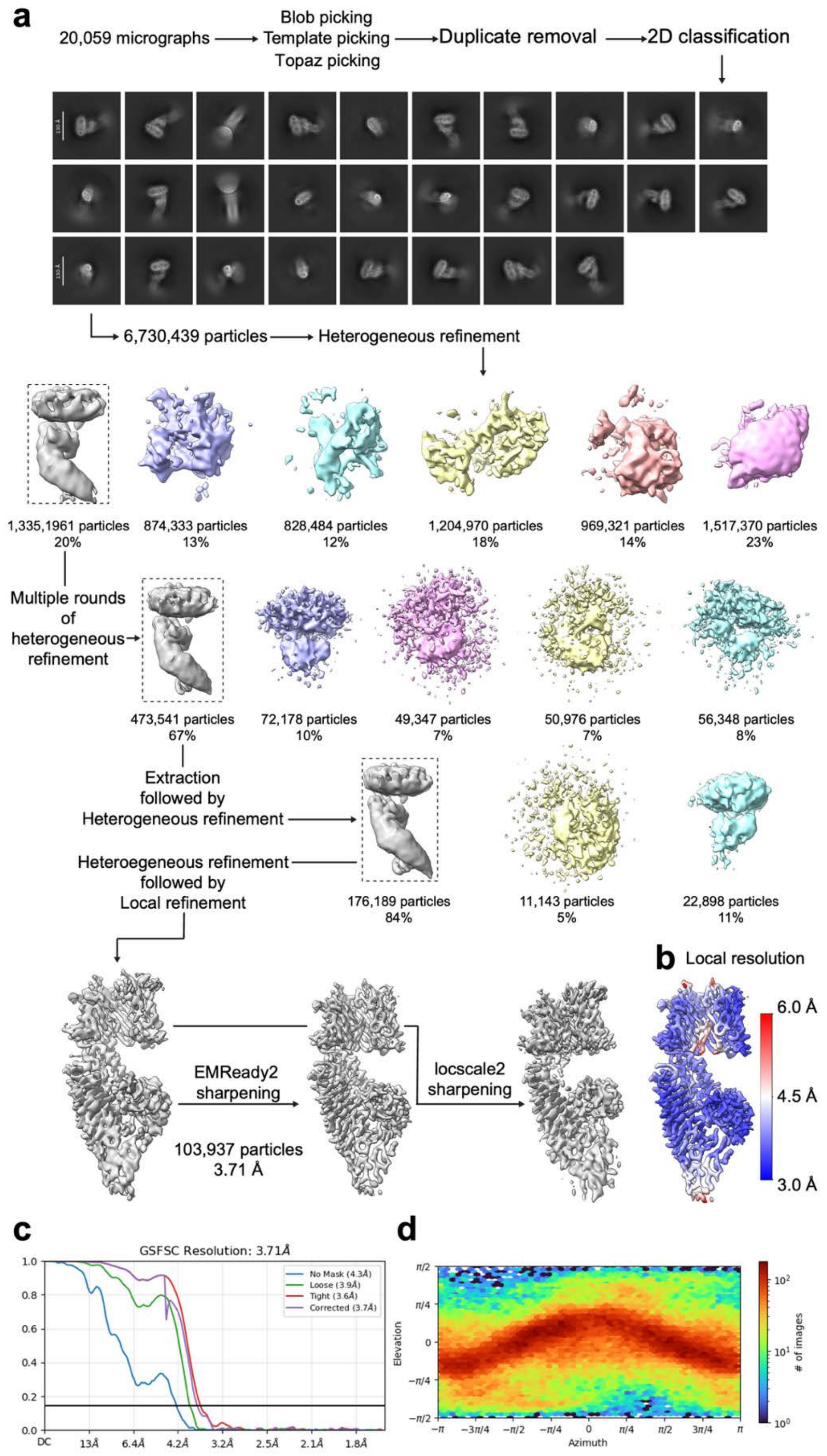

**Figure S5. Cryo-EM reconstruction workflow for $^{His}$TamAB490$^{TS}$ phospholipid nanodiscs.**
(**a**) Workflow for processing of cryo-EM dataset collected on $^{His}$TamAB490$^{TS}$ in phospholipid

nanodiscs using CryoSPARC. Classes from intermediate rounds of heterogeneous refinement and the final map from local refinement and sharpening using either EMReady2 or LocScale2. (**b**) Map of $^{His}$TamAB490$^{TS}$ in phospholipid nanodiscs colored by local resolution (CryoSPARC local resolution estimation). (**c**) Gold-standard Fourier-shell correlation (GSFSC) for $^{His}$TamAB490$^{TS}$ in phospholipid nanodiscs. (**d**) Orientation distribution for all particles used in the reconstruction of $^{His}$TamAB490$^{TS}$ in phospholipid nanodiscs.

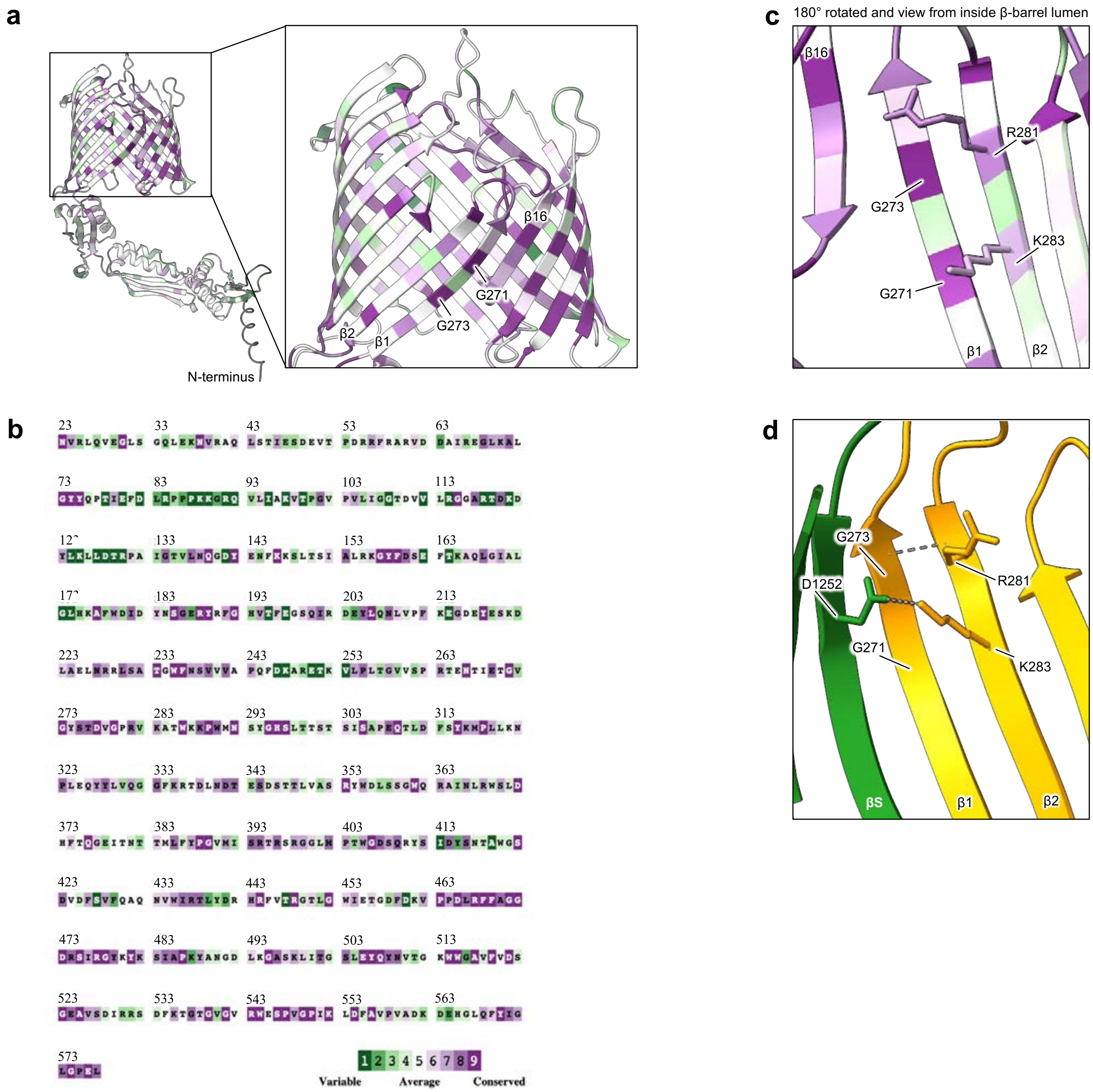

**Figure S6. Conservation of TamA residues.**
The conservation of TamA residues was calculated using ConSurf[1]. The color scale for conservation level was applied to (**a**) the structure of TamA (AF2, similar to TamA crystal structure [PDB ID: 4C00][2] but includes missing segments) and (**b**) the aa sequence of TamA. Inset: magnified view of conservation of residues in the TamA β-barrel. (**c**) β-barrel lumen view of highly conserved glycine residues in TamA β-strand 1 (β1) (G271, G273). Conserved charged residues (R281, K283) in neighboring β2 that are in-register with the β1 glycine residues. (**d**) $^{His}$TamAB490$^{TS}$-nanodisc structure with same view as in **c**. The space retained by G271 and G273 (lacking luminal R groups) might be required for the inter-strand salt-bridge that forms between TamA K283 in β2 forms a salt-bridge with TamB D1252 in the β-signal (βS) strand.

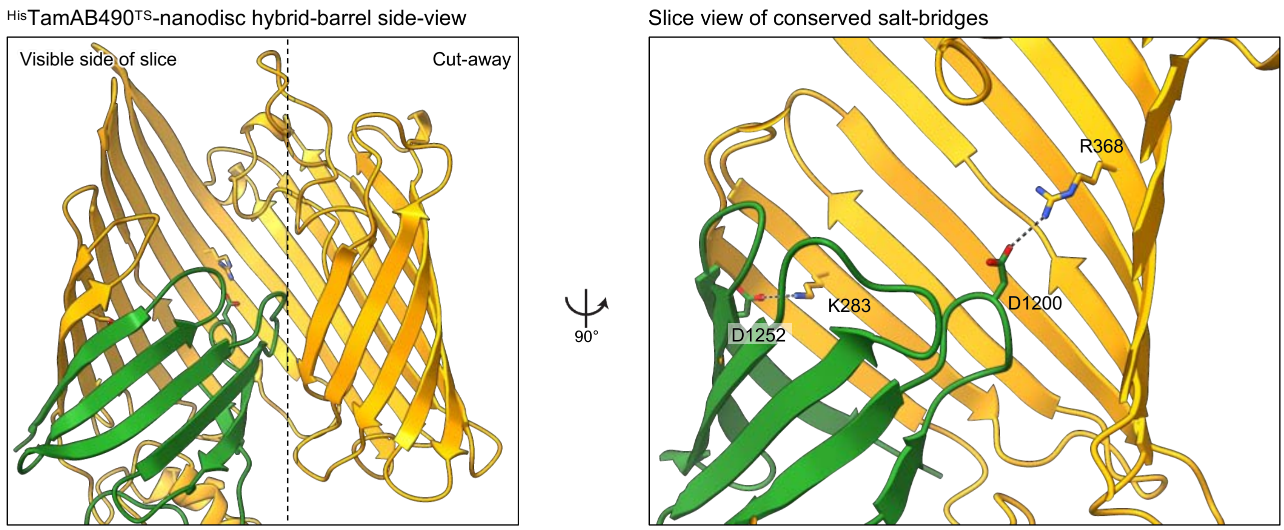

**Figure S7. Conserved intermolecular salt-bridges in the TAM hybrid-barrel .**
Left, $^{His}$TamAB40$^{TS}$-nanodisc showing a side view of the hybrid-barrel between TamA (orange) and TamB (dark-green). Right, rotated slice-view to show the location of two TamA-TamB salt-bridges formed between conserved residue pairs TamA$_{K283}$-TamB$_{D1252}$ and TamA$_{R368}$-TamB$_{D1200}$. Conservation levels are in Fig. S2 and S6.

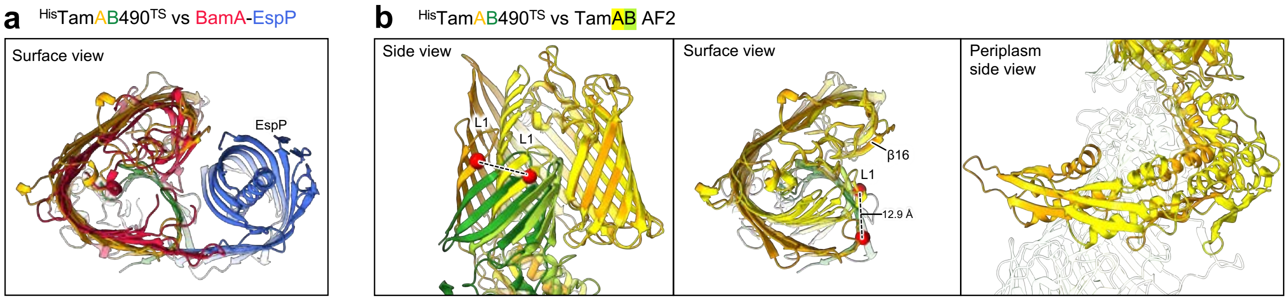

**Figure S8. The conformation of the $^{His}$TamAB490$^{TS}$-nanodisc structure is similar to the transient hybrid-barrel formed by BamA during OMP assembly.**

Model alignments as in Fig. 1f,g. (**a**) Similarity of $^{His}$TamAB490$^{TS}$-nanodisc structure to the cryo-EM structure of BAM-$^{pair1}$-EspP (PDB ID: 8BNZ)[3]. (**b**) Comparison of $^{His}$TamAB490$^{TS}$-nanodisc structure (orange, TamA; dark-green, TamB) to the AlphaFold2 (AF2) predicted structure of TAM (yellow, TamA; lime-green, TamB). The AF2 structure is in a compressed conformation similar to TamAB$^{His}$-amphipol and the TamA-alone structures.

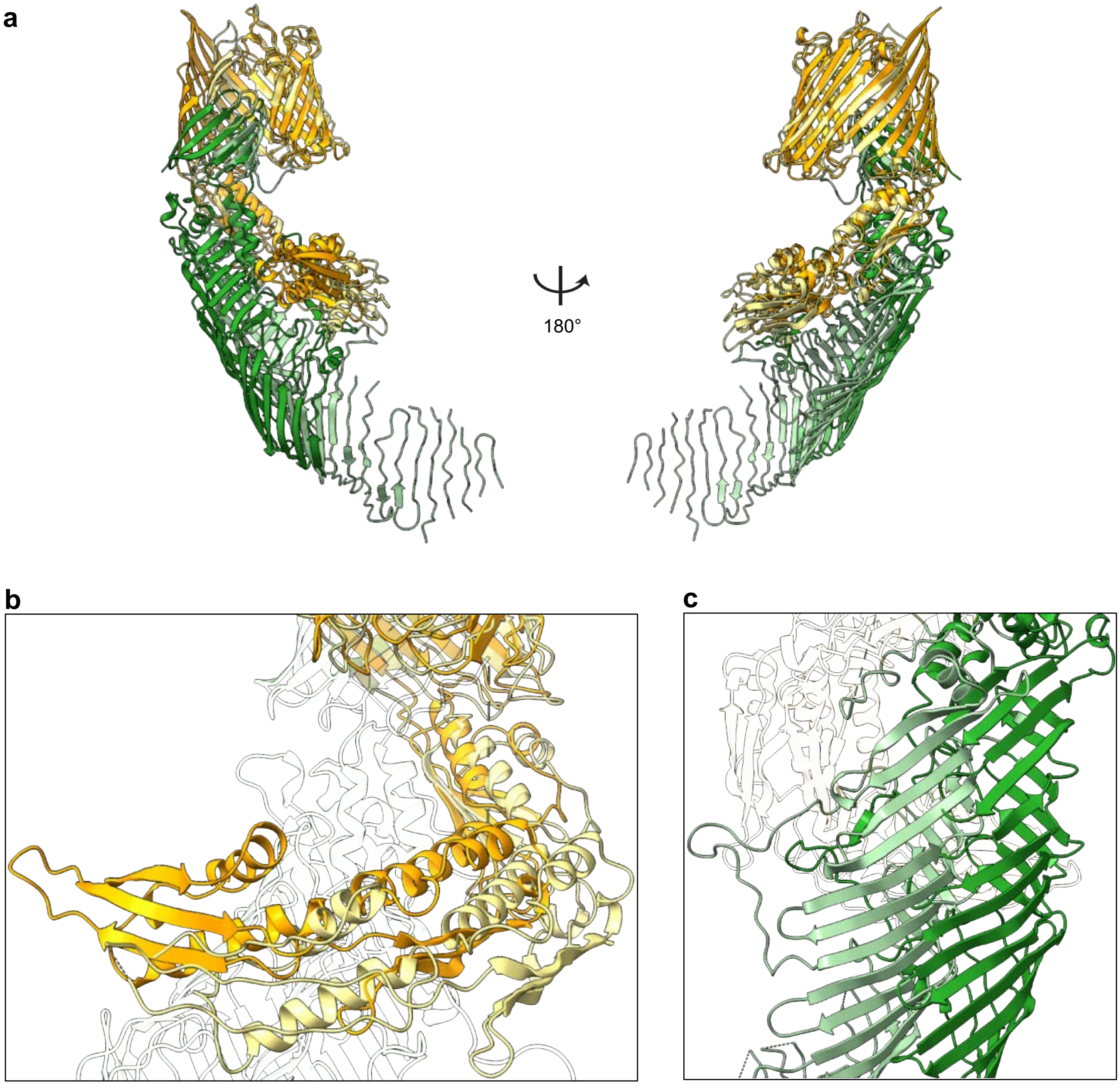

**Figure S9. Extended comparison of $^{His}$TamAB490$^{TS}$-nanodisc and TamAB$^{His}$-amphipol structures.**
(**a**) Comparison of $^{His}$TamAB490$^{TS}$-nanodisc (TamA, orange; TamB dark-green) to TamAB$^{His}$-amphipol (TamA, cream; TamB, teal-green; PDB ID: 9XDC)[4]. Models were aligned on TamA β-barrel α-carbons Y440-L577. The difference in hybrid-barrel conformation between the two structures (see Fig. 1g) corresponds to a major change in the orientation of the TamA POTRA domains (**b**) and the TamB β-taco (**c**).

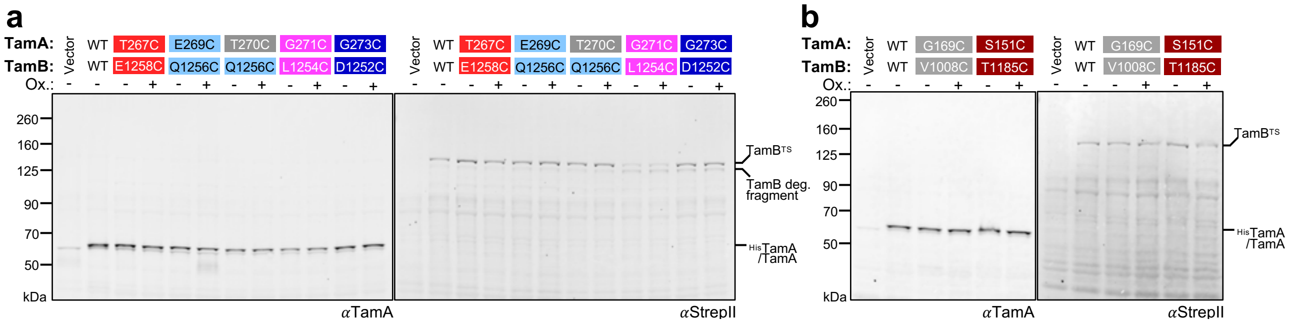

**Figure S10. Reduced SDS-PAGE of disulfide crosslinking samples.**
Aliquots of samples from Fig. 2b (**a**) and Fig. 4c (**b**) heated to 99°C with dithiothreitol (DTT; 150 mM) prior to SDS-PAGE. Western immunoblots were doubly probed with antibodies against TamA (αTamA) and the TS-tag in TamB (αStrepII).

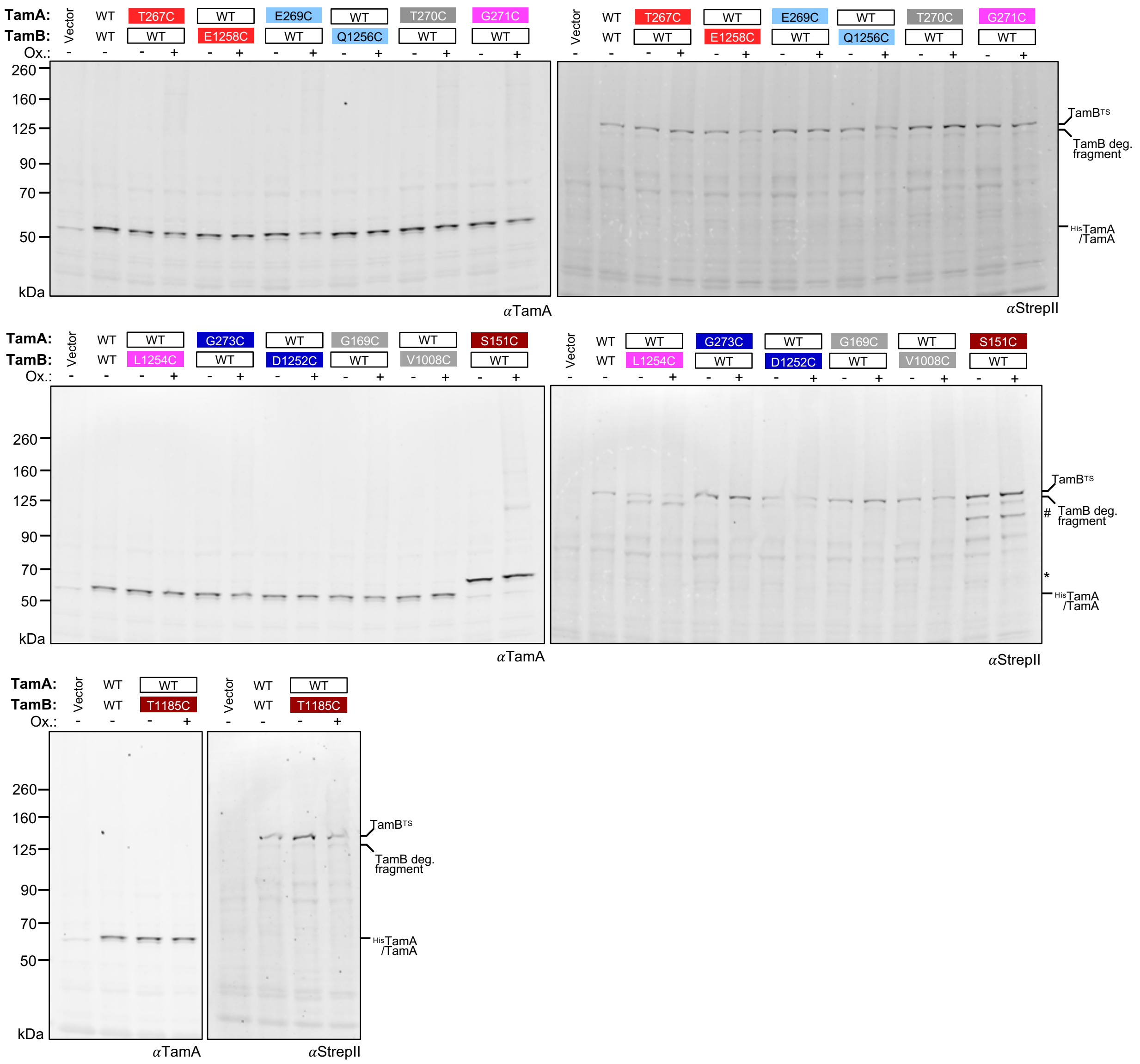

**Figure S11. Single cysteine substitution controls for disulfide crosslinking experiments.** Experiment conducted as in Fig. 2b except using *E. coli* expressing wild-type (WT) $^{His}$TamAB$^{TS}$, or derivatives with single cysteine substitutions as controls. Western immunoblots were doubly probed with antibodies against TamA (αTamA) and the TS-tag in TamB (αStrepII). #, additional degradation products. *, putative alternative adduct in the absence of single cysteine.

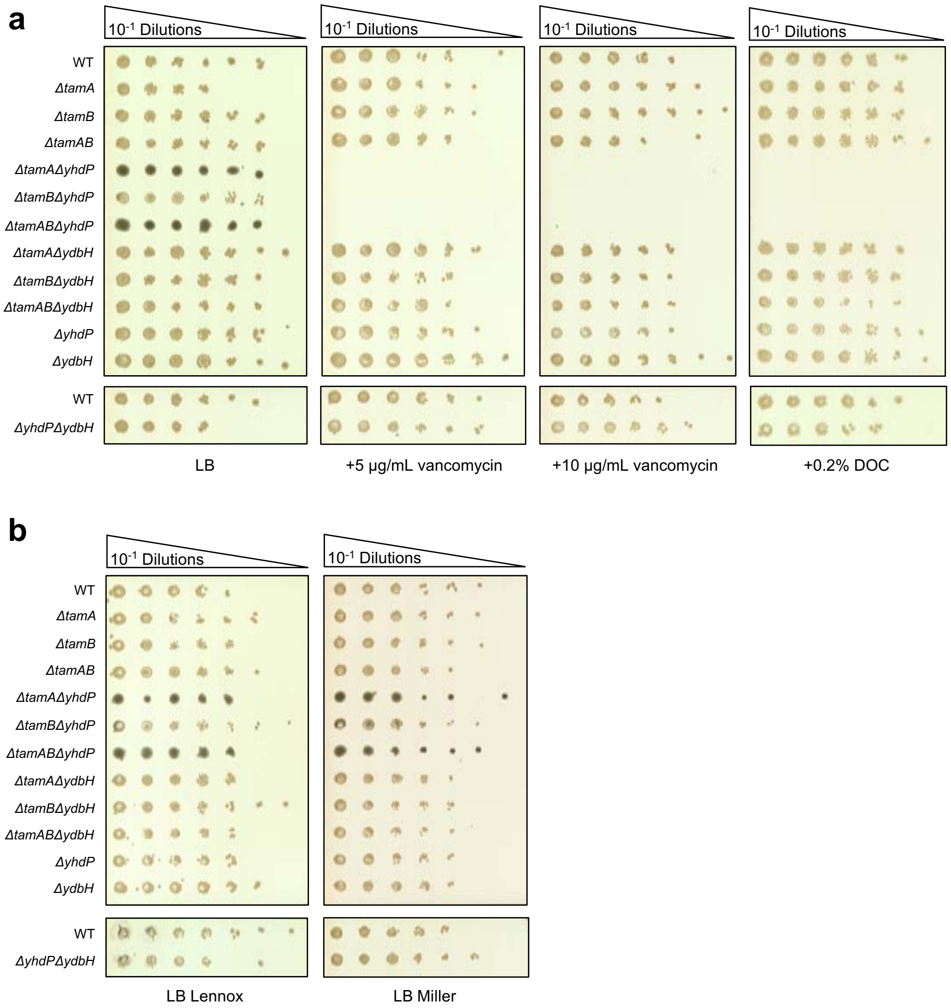

**Figure S12. Simultaneous deletion of lipid bridge encoding genes *tamAB* and *yhdP* result in stress responses and outer membrane defects.**

(**a**) Efficiency of plating assay as in Fig. 1c. Serial dilutions of WT or mutant derivatives of *E. coli* K-12 W3110 were spotted onto plain LB agar plates or plates containing 5 or 10 µg/mL vancomycin, or 0.2% deoxycholate (DOC), n = 3. Plates were incubated at 37 °C for 20 hr. Deletions of *tam* genes together with *yhdP* result in a strong mucoid stress response that is apparent as dark colonies on transillumination. (**b**) For all experiments in the current work we have used the Lennox formulation of LB. Here, an efficiency of plating assay was conducted to compare phenotypes of the strains in **a** between grown on LB Lennox or LB Miller (double NaCl concentration) noting a mild increase in mucoidy on LB Miller.

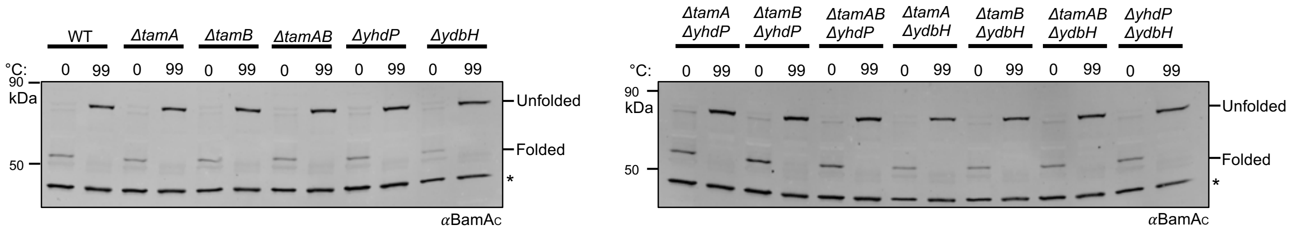

**Figure S13. The assembly of BamA is unaffected by deletion of lipid transport genes.** Heat-modifiable electrophoretic shift assay. Wild-type (WT) *E. coli* K-12 W3110 or gene deletion mutant derivatives thereof where cultured in liquid LB at 37 °C until log phase and lysates where either unheated (0 °C) or heated to 99 °C and proteins were resolved by cold SDS-PAGE. Folded states of endogenous BamA were detected by Western immunoblotting with an antibody against the C-terminus of BamA (αBamA$_C$).

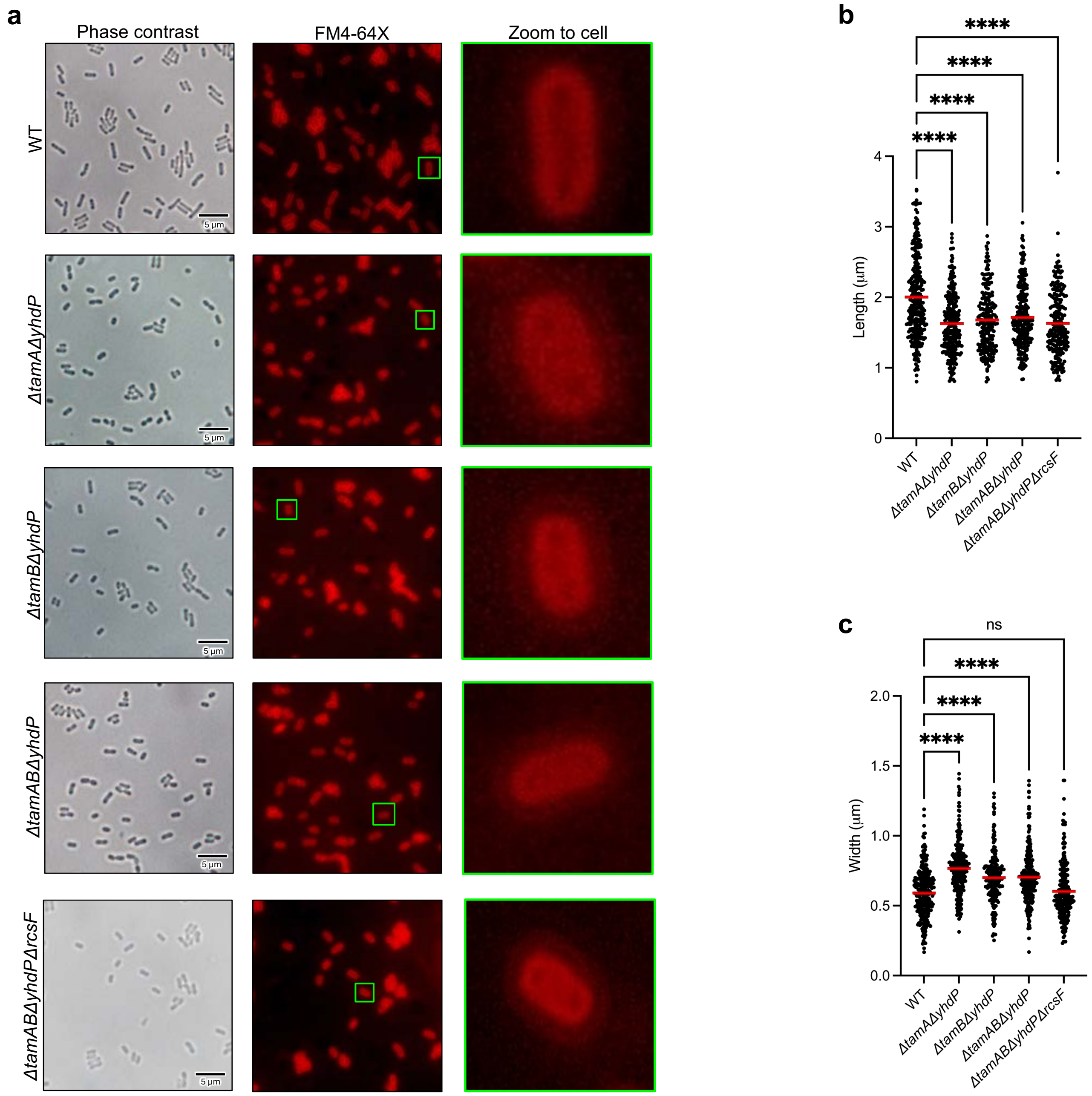

**Figure S14. Analysis of WT and lipid bridge deletion mutants by microscopy.**
(**a**) WT or mutant derivatives of *E. coli* K-12 W3110 were grown in LB at 37 °C overnight and incubated with FM4-64X, a stain that preferentially labels the outer membrane[5], n = 3. Live stained bacteria where imaged on LB agarose with an Olympus BX51 fluorescence microscope. Cell lengths (**b**) and widths (**c**) were quantified using MicrobeJ[6]. Red lines show mean value.

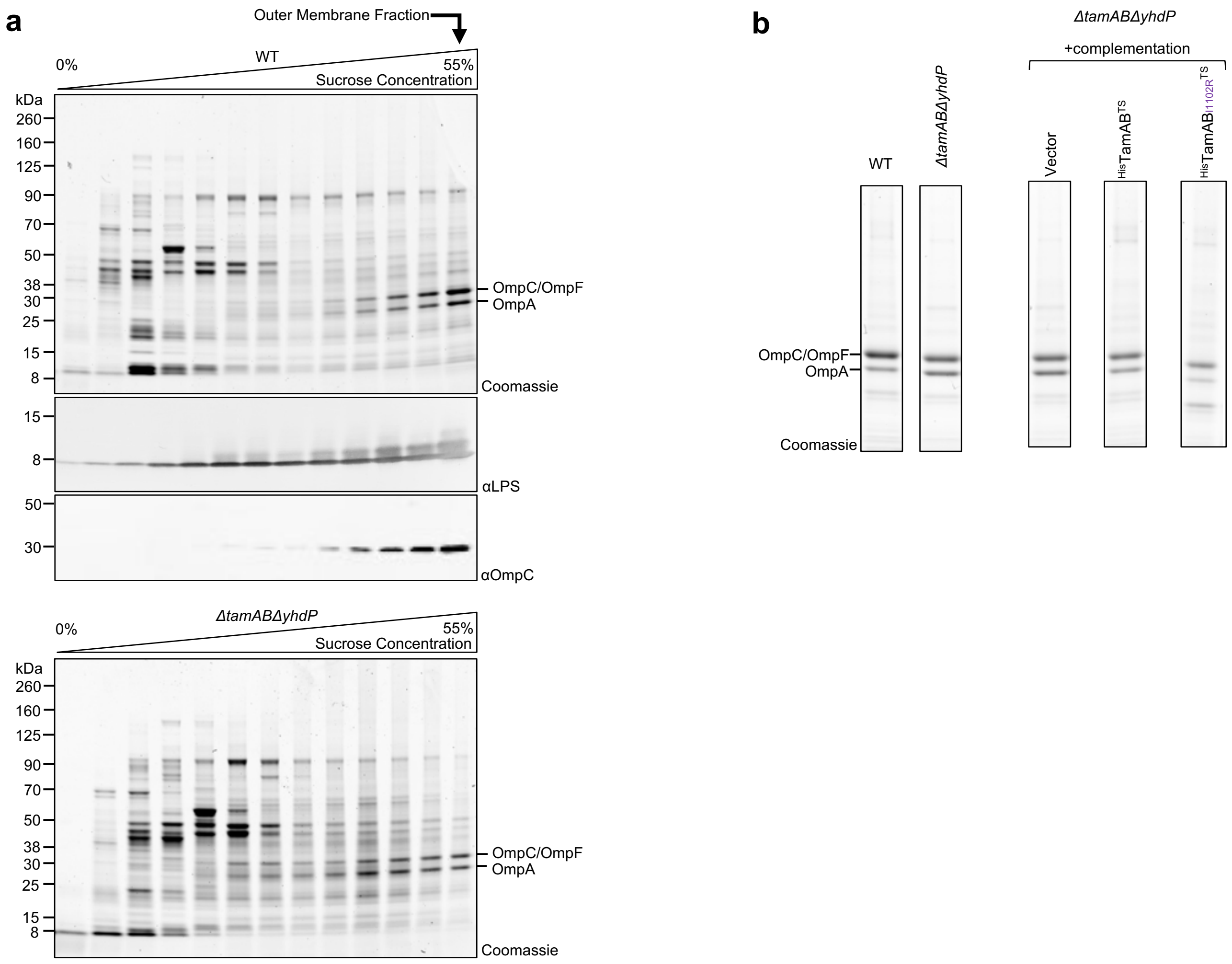

**Figure S15. Isolation of *E. coli* outer membranes by sucrose density gradient.**

(**a**) WT *E. coli* K-12 W3110 and *ΔtamABΔyhdP* were grown in LB at 37 °C overnight, lysed, and membrane fractions resolved by ultracentrifugation through a 0-55% sucrose density gradient. Fractions were separated by SDS-PAGE and analyzed by Coomassie staining and western immunoblotting with antibodies for the detection of lipopolysaccharide (αLPS) or OmpC (αOmpC). The final high-density fraction in which major outer membrane proteins OmpC/F and OmpA, and LPS were accumulated was designated the outer membrane fraction and retained for analysis. (**b**) Examples of outer membrane fractions obtained for strains of interest for phospholipidomics analysis, relevant to Fig 2d and 5c.

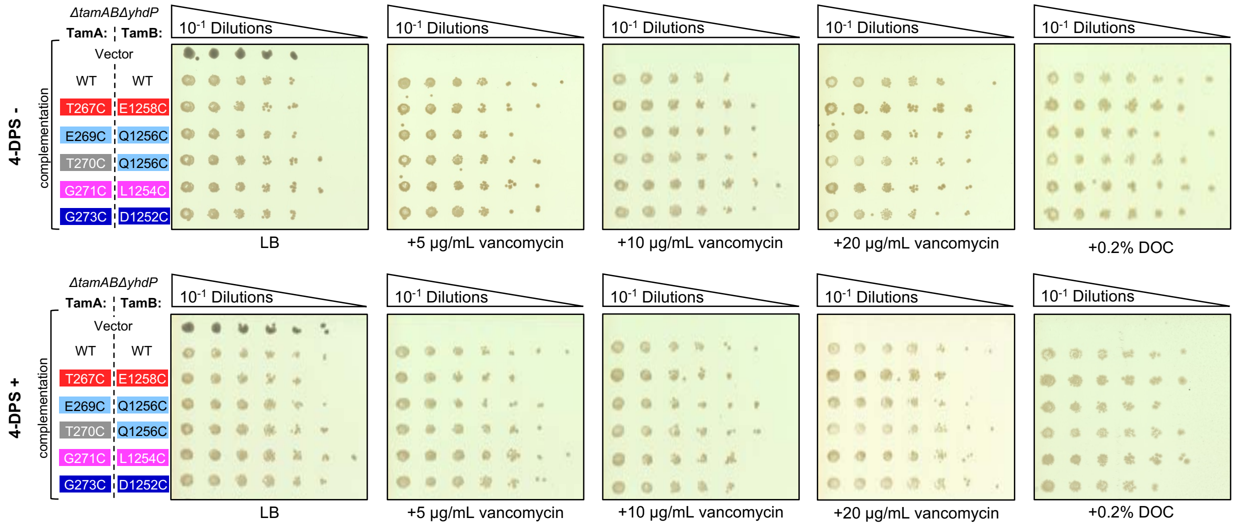

**Figure S16. Additional conditions tested for TAM derivatives with disulfide-tethered hybrid-barrel interface.**

Additional conditions tested in experiment in Fig. 2f wherein strains were plated in the presence or absence of vancomycin (5 µg/mL, 10 µg/mL, 20 µg/mL), sodium deoxycholate (DOC, 0.2%) with or without the thiol specific oxidizer 4-DPS (50 µM), n = 3.

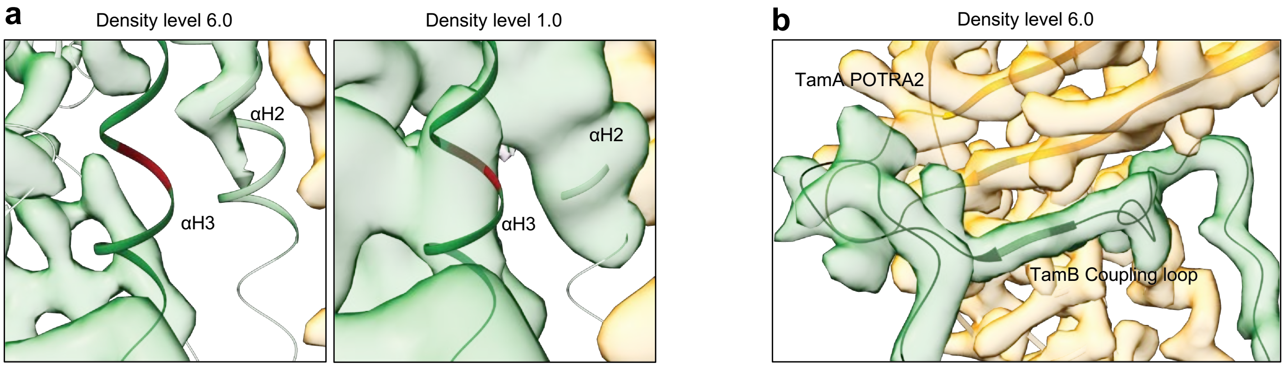

**Figure S17. Density of important periplasmic segments in $^{His}$TamAB490$^{TS}$-nanodisc cryo-EM map.**
(**a**) Variable density corresponding to α-helix 3 (αH3) in TamB can be observed at lower map thresholds. See also Video S13-15 for 3D variability analysis relevant to this region. (**b**) Strong density corresponding to a stable interaction between TamA POTRA2 and the TamB coupling-loop.

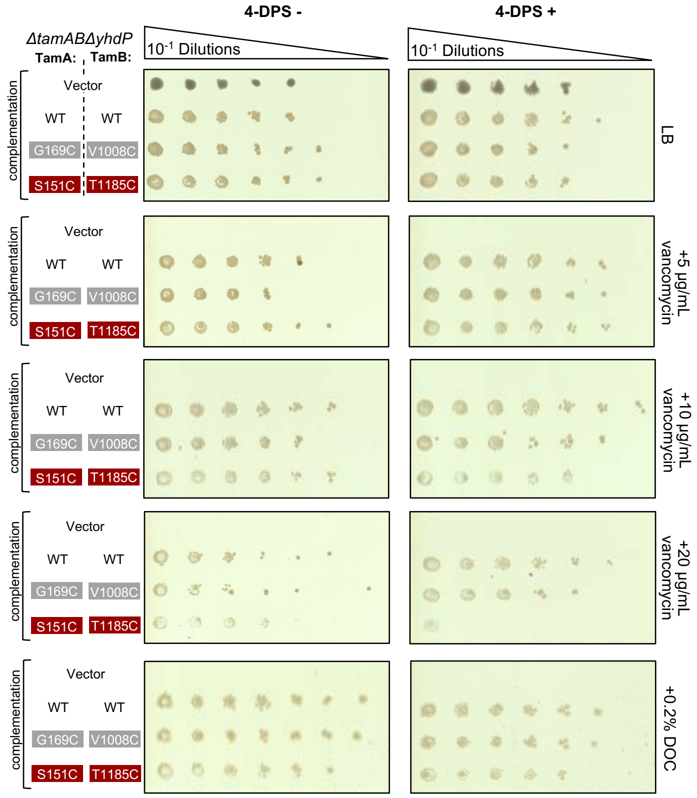

**Figure S18. Additional conditions tested for TAM derivatives with disulfide-tethered periplasmic segments.**

Additional conditions tested in experiment in Fig. 4d wherein strains were plated in the presence or absence of vancomycin (5 µg/mL, 10 µg/mL, 20 µg/mL), sodium deoxycholate (DOC, 0.2%), with or without the thiol specific oxidizer 4-DPS (50 µM), n = 3.

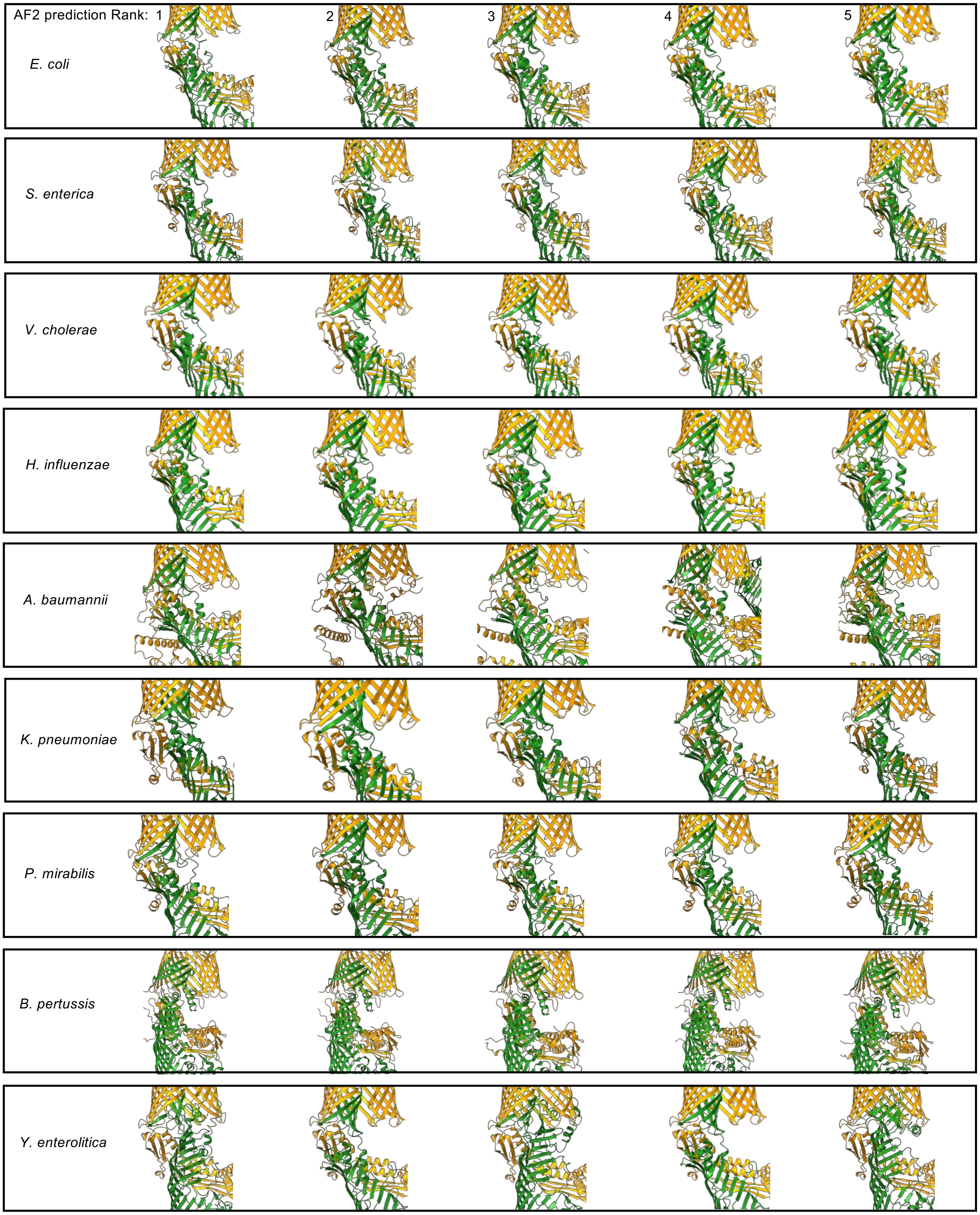

**Figure S19. Variability in DUF490 helices between TAM predictions. (cont. next pg.)**

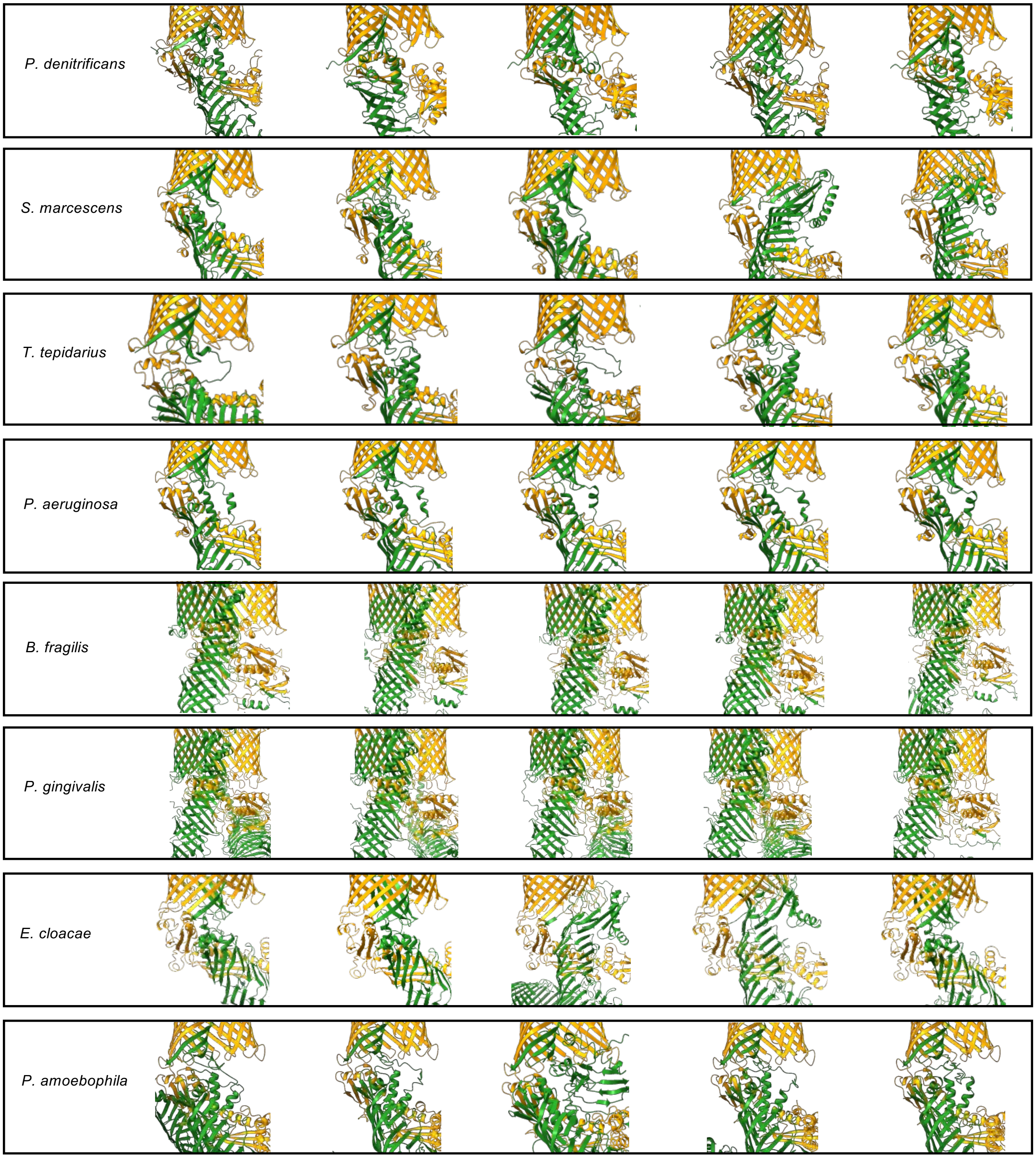

**Figure S19. Variability in DUF490 helices between TAM predictions.**
Comparisons of AlphaFold2 (AF2) ranked models for each species predicted TAM and region of DUF480 helices and C-terminal membrane embedded hybrid-barrel strands magnified. A trend of uncertainty in helix position and structure between models is evident.

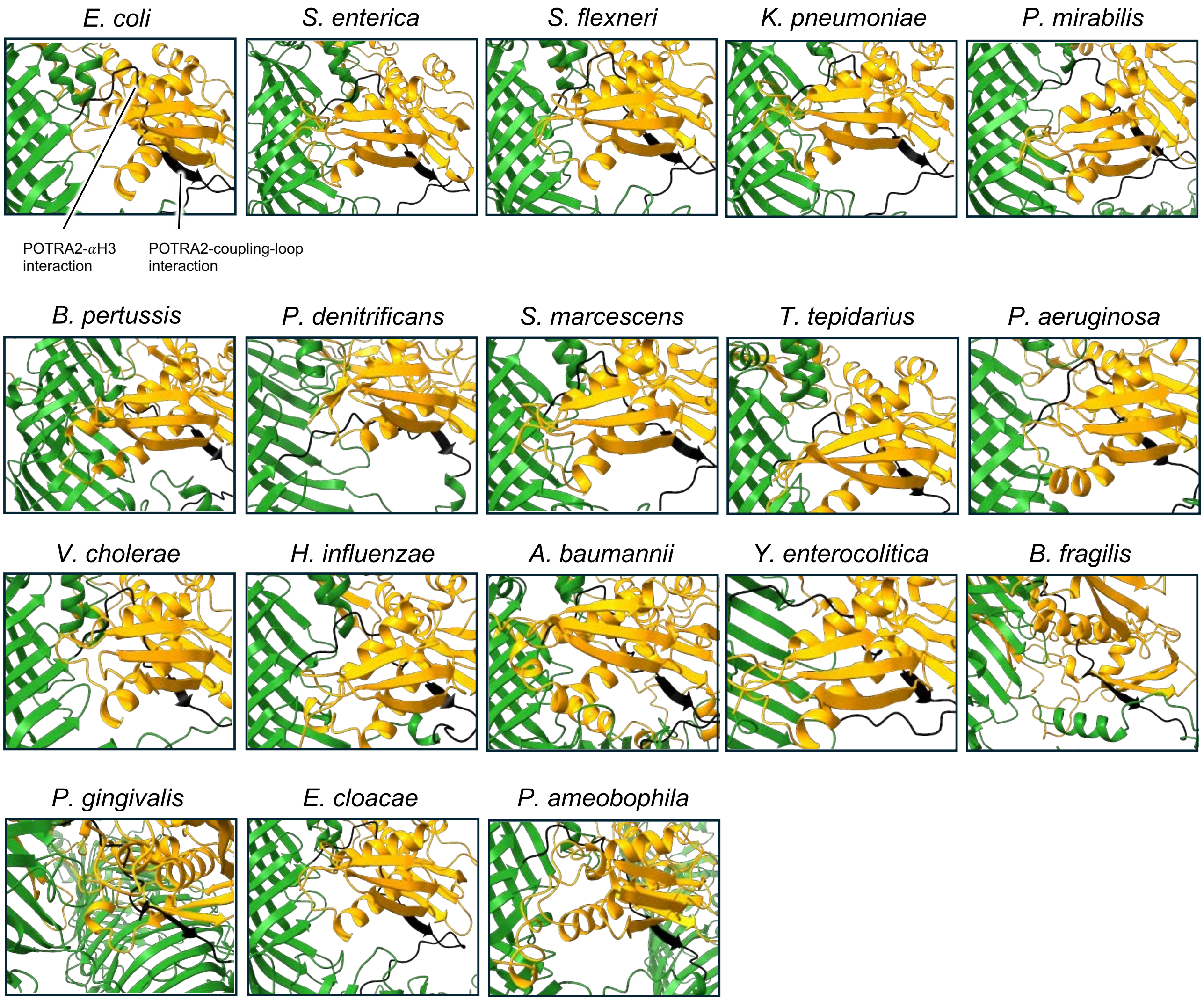

**Figure S20. Prediction of conserved POTRA2-DUF490 coupling-loop interaction.** Comparisons of AlphaFold2 for each species predicted TAM with the interaction between TamA POTRA2 (orange) and the TamB DUF490 coupling-loop (black) magnified. The interaction is consistently predicted between species and with high confidence and suggests that this is a stable interaction wherein POTRA2 induces a β-strand conformation in the coupling-loop via β-augmentation.

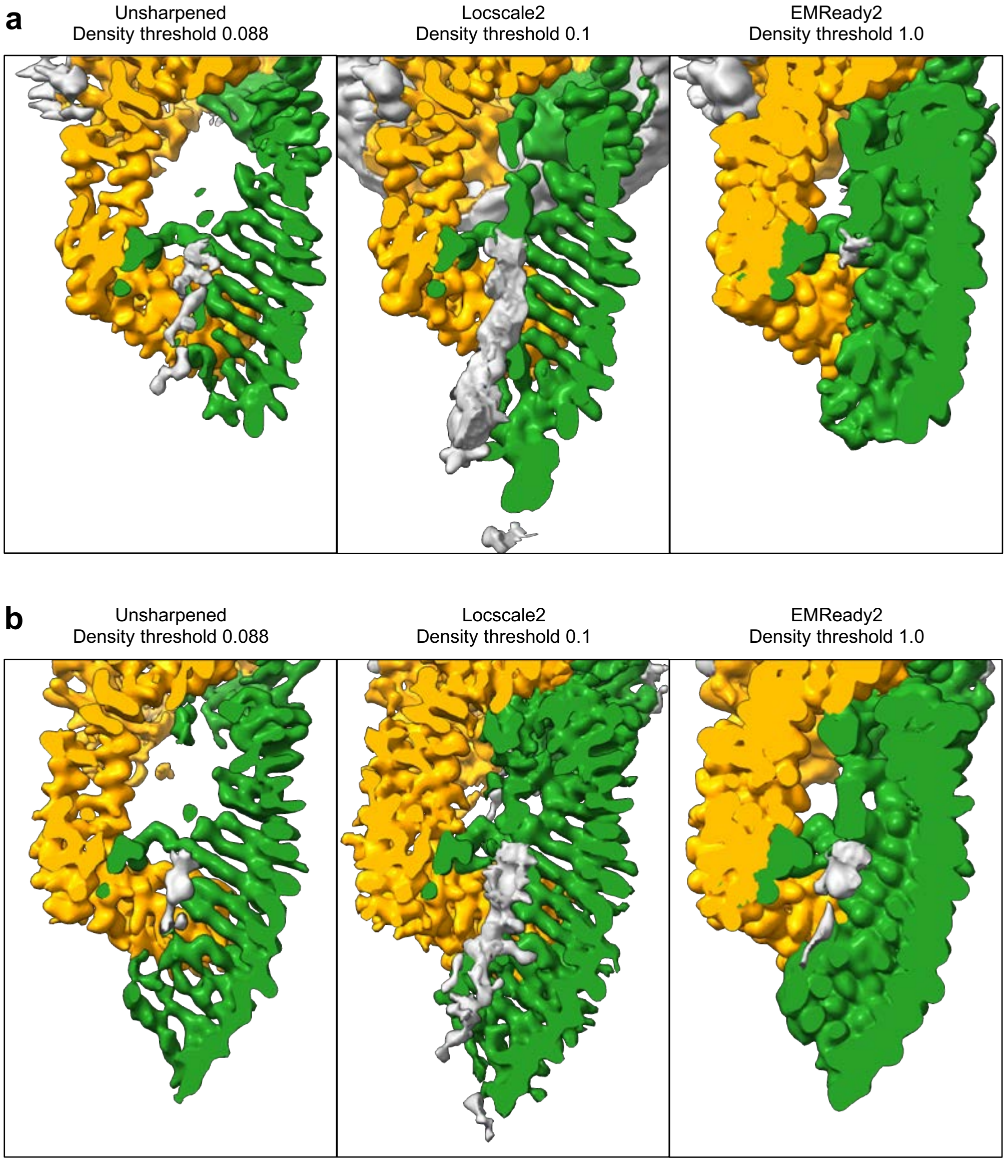

**Figure S21. Additional densities in the cryo-EM maps of $^{His}$TamAB490$^{TS}$.**
$^{His}$TamAB490$^{TS}$-LMNG detergent (**a**) or $^{His}$TamAB490$^{TS}$-nanodisc (**b**) cryo-EM maps shown as a vertical slice across the lipophilic TamB β-taco as in Fig. 5a. TamA (orange), TamB (green), and density not assigned to either subunit (grey). Unsharpened maps from cryoSPARC refinement are shown alongside results from sharpening algorithms Locscale2[7,8] and EMReady2[9].

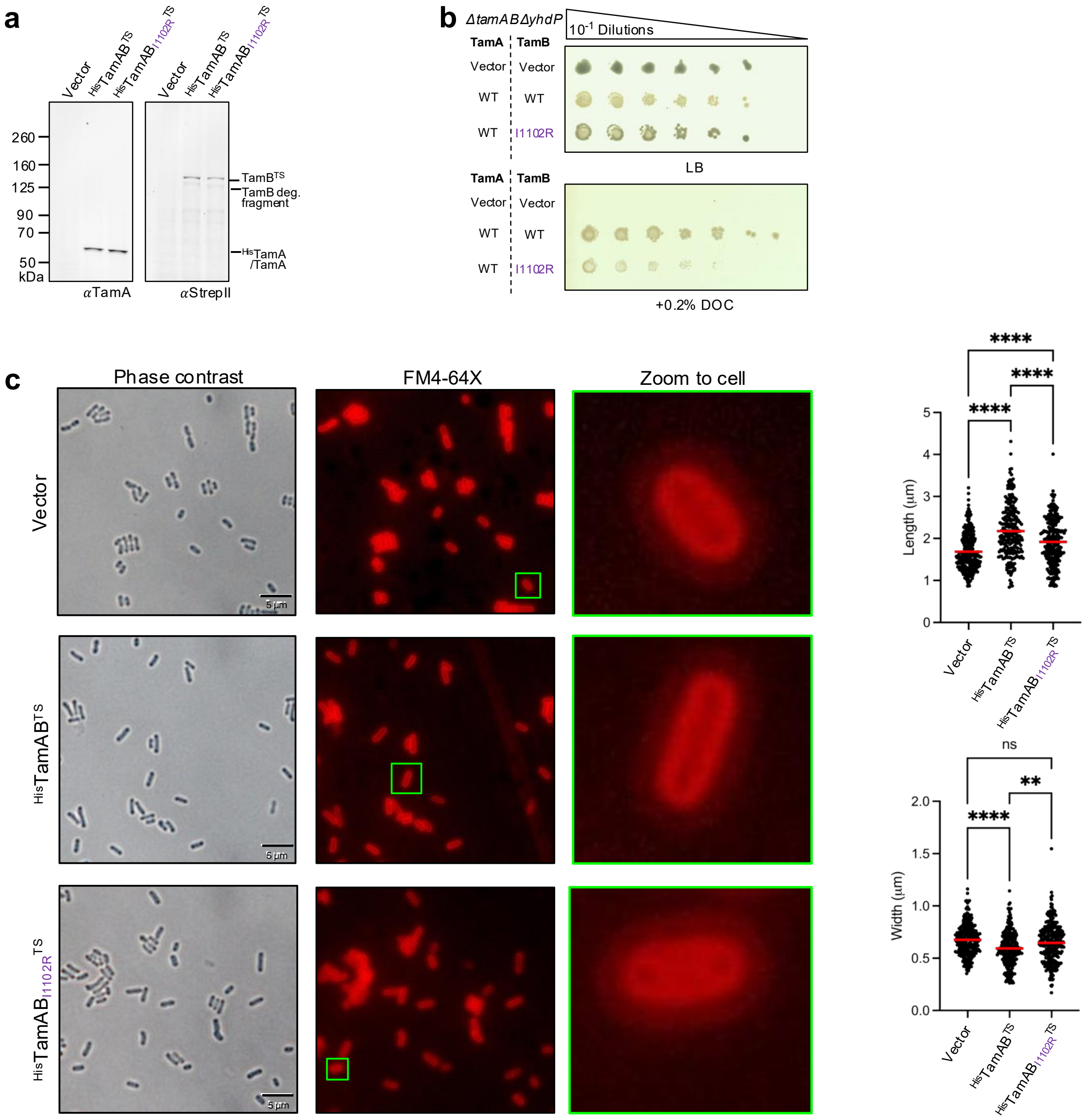

**Figure S22. TAM dysfunction due to I1102R substitution in the TamB lipid channel.**
(**a**) *E. coli* K-12 W3110 *ΔtamABΔyhdP* strain complemented with empty pTrc99a or harboring genes for expression of $^{His}$TamAB$^{TS}$, or $^{His}$TamAB$_{I1102R}^{TS}$ were grown in LB and overnight at 37 °C and total protein extracts were probed by immunoblotting with either of αTamA and the TS-tag in TamB (αStrepII), n = 3. (**b**) Experiment as in Fig. 4b with additional growth of strains on deoxycholate (DOC, 0.2%), n = 3. (**c**) Strains were grown as in **c** and then microscopy of live bacteria conducted as in Fig. S14, n = 3. Left, example micrographs. Right, Length and widths quantified.

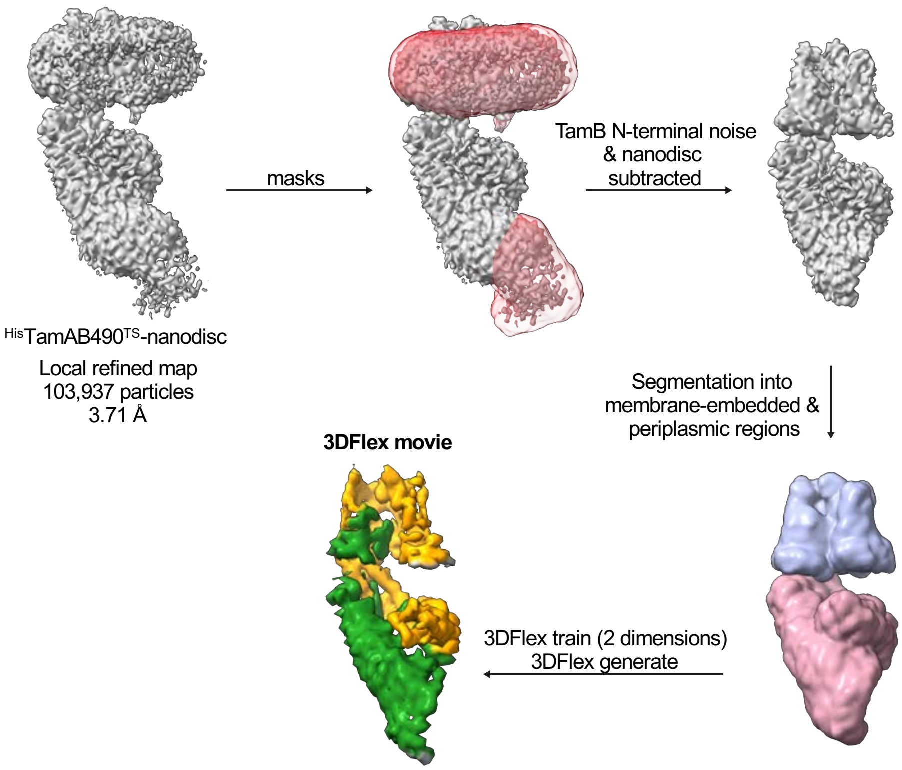

**Figure S23. Analysis of the conformational dynamics of TAM subunits.**
Processing workflow using cryoSPARC 3DFlex analysis on the particles used for $^{His}$TamAB490$^{TS}$-nanodisc reconstruction. See Video S19 and S20.

### SUPPLEMENTARY TABLES

**Table S1. DUF490-containing proteins.**
See supplemental file "Table S1.xlsx". Proteins within the UniProKB containing a DUF490 were identified using the IPR007452/PF04357 HHM, redundancy reduced by UniRef50_P39321, and pruned to contain proteins of at least 800aa.

**Table S2. Statistical analyses for all experiments except phospholipidomics.**
See supplemental file "Table S2.xlsx".

**Table S3. Abundances of phospholipids relevant to Fig. 2d.**

| | ABUNDANCE (% of Total) | | | | Red = p.value between 0.05-0.01. | | | |
|---|---|---|---|---|---|---|---|---|
| | WT | | ΔtamABΔyhdP | | Sorted based on WT means. | | | |
| Lipid | Mean | SEM | Mean | SEM | p.value | -LOG10(p) | FDR | Alias |
| CL(15:0_16:0_16:1_17:1)-H | 1.2301 | 0.0588 | 0.5340 | 0.0836 | 0.0005 | 3.3088 | 0.0118 | j |
| CL(16:0_16:0_16:1_18:1)-H | 1.1657 | 0.0718 | 0.7518 | 0.1522 | 0.0491 | 1.3086 | 0.1443 | |
| CL(16:0_16:0_16:1_17:1)-H | 1.0650 | 0.0587 | 0.7600 | 0.1282 | 0.0737 | 1.1323 | 0.1948 | |
| CL(16:0_16:0_16:0_16:1)-H | 0.5924 | 0.0370 | 0.3053 | 0.0703 | 0.0111 | 1.9532 | 0.0501 | |
| CL(14:0_16:0_16:0_16:1)-H | 0.4603 | 0.0214 | 0.2528 | 0.0440 | 0.0054 | 2.2657 | 0.0314 | k |
| CL(16:0_16:0_17:1_18:1)-H | 0.4436 | 0.0314 | 0.4307 | 0.0935 | 0.9000 | 0.0458 | 0.9529 | |
| CL(15:0_16:0_16:0_16:1)-H | 0.3123 | 0.0143 | 0.2676 | 0.0481 | 0.4082 | 0.3891 | 0.5763 | |
| CL(14:0_16:0_16:1_17:1)-H | 0.2606 | 0.0113 | 0.2154 | 0.0290 | 0.1973 | 0.7050 | 0.3839 | |
| CL(16:0_16:0_16:0_17:1)-H | 0.2383 | 0.0250 | 0.2173 | 0.0373 | 0.6575 | 0.1821 | 0.7890 | |
| CL(16:0_16:0_18:1_18:1)-H | 0.2247 | 0.0132 | 0.1958 | 0.0418 | 0.5339 | 0.2725 | 0.6865 | |
| CL(14:0_16:0_16:1_16:1)-H | 0.2005 | 0.0091 | 0.1207 | 0.0110 | 0.0014 | 2.8562 | 0.0125 | l |
| CL(16:0_16:0_16:0_18:1)-H | 0.1990 | 0.0238 | 0.1364 | 0.0294 | 0.1485 | 0.8284 | 0.3239 | |
| CL(48:2)-H | 0.1881 | 0.0086 | 0.0931 | 0.0204 | 0.0052 | 2.2871 | 0.0314 | m |
| CL(16:0_16:1_16:1_18:1)-H | 0.1785 | 0.0085 | 0.0938 | 0.0115 | 0.0010 | 2.9825 | 0.0125 | n |
| CL(16:0_16:1_18:1)-H | 0.1614 | 0.0048 | 0.1348 | 0.0270 | 0.3700 | 0.4318 | 0.5671 | |
| **TOTAL CL** | **6.9204** | | **4.5094** | | | | | |
| PE(16:0_17:1)-H | 23.9317 | 0.7954 | 25.8846 | 1.2823 | 0.2432 | 0.6141 | 0.4490 | |
| PE(16:0_16:1)-H | 17.3141 | 1.0886 | 14.5736 | 1.1109 | 0.1285 | 0.8910 | 0.2986 | |
| PE(16:0_18:1)-H | 8.5455 | 0.1935 | 8.7177 | 0.1295 | 0.4873 | 0.3122 | 0.6497 | |
| PE(14:0_16:0)-H | 6.3279 | 0.2342 | 5.7825 | 0.2295 | 0.1473 | 0.8317 | 0.3239 | |
| PE(16:0_16:0)-H | 4.9365 | 0.3065 | 2.9976 | 0.0789 | 0.0009 | 3.0630 | 0.0125 | a |
| PE(15:0_16:0)-H | 3.8249 | 0.4272 | 3.2751 | 0.1538 | 0.2715 | 0.5663 | 0.4546 | |
| PE(16:0_14:1)-H | 2.4807 | 0.1527 | 2.9195 | 0.1362 | 0.0757 | 1.1207 | 0.1948 | |
| PE(15:0_16:1)-H | 2.2209 | 0.5888 | 3.9817 | 0.9453 | 0.1649 | 0.7827 | 0.3392 | |
| PE(16:1_18:1)-H | 1.3791 | 0.0926 | 1.8302 | 0.0696 | 0.0080 | 2.0951 | 0.0412 | b |
| PE(16:0_19:1)-H | 1.1955 | 0.0251 | 2.0121 | 0.1057 | 0.0003 | 3.5411 | 0.0104 | c |
| PE(16:1_17:1)-H | 1.1180 | 0.0544 | 1.4475 | 0.0565 | 0.0057 | 2.2458 | 0.0314 | d |
| PE(17:1_18:1)-H | 0.9306 | 0.0285 | 1.1977 | 0.0373 | 0.0013 | 2.8977 | 0.0125 | e |
| PE(17:1_17:1)-H | 0.9344 | 0.0624 | 1.0354 | 0.0504 | 0.2546 | 0.5942 | 0.4546 | |
| PE(14:0_15:0)-H | 0.8821 | 0.0807 | 1.3272 | 0.1277 | 0.0257 | 1.5892 | 0.0843 | |
| PE(14:0_14:0)-H | 0.6522 | 0.0496 | 0.7217 | 0.0554 | 0.3859 | 0.4135 | 0.5671 | |
| **TOTAL PE** | **76.6741** | | **77.7040** | | | | | |
| PG(16:0_16:1)-H | 4.0666 | 0.1830 | 4.2526 | 0.3387 | 0.6461 | 0.1897 | 0.7885 | |
| PG(16:0_18:1)-H | 1.7104 | 0.0921 | 2.3362 | 0.1649 | 0.0161 | 1.7919 | 0.0684 | |
| PG(16:0_17:1)-H | 1.6249 | 0.0329 | 2.6460 | 0.0322 | 0.0000 | 6.2608 | 0.0000 | f |
| PG(32:0)-H | 1.4822 | 0.0896 | 0.8934 | 0.0405 | 0.0010 | 3.0116 | 0.0125 | g |
| PG(14:0_16:0)-H | 0.7144 | 0.0433 | 0.7115 | 0.0551 | 0.9686 | 0.0139 | 0.9822 | |
| PG(17:0_16:1)-H | 0.6524 | 0.2291 | 0.7673 | 0.1146 | 0.6696 | 0.1742 | 0.7903 | |
| PG(15:0_16:0)-H | 0.6468 | 0.0836 | 0.6470 | 0.0938 | 0.9991 | 0.0004 | 0.9991 | |
| PG(15:0_16:1)-H | 0.3248 | 0.0314 | 0.6628 | 0.1048 | 0.0214 | 1.6692 | 0.0804 | |
| PG(16:0_14:1)-H | 0.2681 | 0.0253 | 0.3533 | 0.0386 | 0.1143 | 0.9419 | 0.2743 | |
| PG(16:0_19:1)-H | 0.2560 | 0.0318 | 0.3466 | 0.0190 | 0.0501 | 1.3002 | 0.1443 | |
| PG(34:2)-H | 0.2091 | 0.0168 | 0.3589 | 0.0575 | 0.0465 | 1.3325 | 0.1443 | |
| PG(33:0)-H | 0.1770 | 0.0298 | 0.1405 | 0.0160 | 0.3230 | 0.4908 | 0.5168 | |

| | | | | | | | | |
|---|---|---|---|---|---|---|---|---|
| PG(36:2)-H | 0.1311 | 0.0136 | 0.2610 | 0.0262 | 0.0045 | 2.3425 | 0.0314 | h |
| PG(36:0)-H | 0.1068 | 0.0360 | 0.1167 | 0.0299 | 0.8393 | 0.0761 | 0.9156 | |
| PG(13:0_15:0)-H | 0.0908 | 0.0092 | 0.1078 | 0.0057 | 0.1696 | 0.7706 | 0.3392 | |
| **TOTAL PG** | **12.4614** | | **14.6017** | | | | | |
| PC(34:0)+H | 0.0229 | 0.0027 | 0.0372 | 0.0103 | 0.2257 | 0.6464 | 0.4277 | |
| PC(32:0)+H | 0.0100 | 0.0038 | 0.0530 | 0.0260 | 0.1538 | 0.8131 | 0.3256 | |
| PC(28:0)+H | 0.1382 | 0.0300 | 0.1616 | 0.0496 | 0.7007 | 0.1545 | 0.8137 | |
| PC(30:0)+H | 0.0120 | 0.0018 | 0.0083 | 0.0025 | 0.2698 | 0.5689 | 0.4546 | |
| PC(36:1)+H | 0.0049 | 0.0012 | 0.0085 | 0.0046 | 0.4742 | 0.3241 | 0.6497 | |
| PC(34:1)+H | 0.0504 | 0.0204 | 0.0617 | 0.0322 | 0.7768 | 0.1097 | 0.8605 | |
| PC(36:2)+H | 0.0448 | 0.0310 | 0.0834 | 0.0640 | 0.6067 | 0.2170 | 0.7532 | |
| PC(34:2)+H | 0.0341 | 0.0083 | 0.0456 | 0.0189 | 0.5963 | 0.2246 | 0.7532 | |
| **TOTAL PC** | **0.3172** | | **0.4592** | | | | | |
| PA(16:0_16:1)-H | 0.1456 | 0.0200 | 0.0769 | 0.0109 | 0.0234 | 1.6300 | 0.0804 | |
| PA(16:1_18:1)-H | 0.0198 | 0.0034 | 0.0179 | 0.0039 | 0.7194 | 0.1431 | 0.8221 | |
| PA(14:0_14:0)-H | 0.0035 | 0.0007 | 0.0030 | 0.0003 | 0.5100 | 0.2924 | 0.6677 | |
| PA(14:0_16:1)-H | 0.0110 | 0.0022 | 0.0111 | 0.0021 | 0.9609 | 0.0173 | 0.9822 | |
| **TOTAL PA** | **0.1799** | | **0.1088** | | | | | |
| LPE(16:1)-H | 1.0162 | 0.0945 | 0.5838 | 0.0613 | 0.0086 | 2.0668 | 0.0412 | q |
| LPE(16:0)-H | 0.7699 | 0.0660 | 0.3747 | 0.0313 | 0.0016 | 2.7838 | 0.0132 | r |
| LPE(18:1)-H | 0.4364 | 0.0383 | 0.3087 | 0.0414 | 0.0640 | 1.1936 | 0.1773 | |
| LPE(14:0)-H | 0.1795 | 0.0210 | 0.1100 | 0.0080 | 0.0212 | 1.6728 | 0.0804 | |
| LPE(17:0)-H | 0.0619 | 0.0083 | 0.0742 | 0.0100 | 0.3791 | 0.4213 | 0.5671 | |
| LPE(15:0)-H | 0.0645 | 0.0042 | 0.0874 | 0.0115 | 0.1103 | 0.9573 | 0.2739 | |
| LPE(19:1)-H | 0.0516 | 0.0030 | 0.0590 | 0.0071 | 0.3768 | 0.4239 | 0.5671 | |
| LPE(18:0)-H | 0.0119 | 0.0010 | 0.0117 | 0.0017 | 0.9374 | 0.0281 | 0.9782 | |
| LPE(15:1)-H | 0.0111 | 0.0014 | 0.0234 | 0.0038 | 0.0226 | 1.6455 | 0.0804 | |
| **TOTAL LPE** | **2.6030** | | **1.6329** | | | | | |
| LPC(19:0)+H | 0.1460 | 0.0385 | 0.1651 | 0.0466 | 0.7627 | 0.1177 | 0.8580 | |
| LPC(16:0)+H | 0.0165 | 0.0026 | 0.0313 | 0.0116 | 0.2597 | 0.5855 | 0.4546 | |
| LPC(18:0)+H | 0.0066 | 0.0019 | 0.0143 | 0.0083 | 0.4011 | 0.3968 | 0.5763 | |
| LPC(18:1)+H | 0.0045 | 0.0011 | 0.0055 | 0.0008 | 0.4788 | 0.3199 | 0.6497 | |
| **TOTAL LPC** | **0.1736** | | **0.2162** | | | | | |
| LPA(17:1)-H | 0.6432 | 0.1412 | 0.6920 | 0.2247 | 0.8602 | 0.0654 | 0.9243 | |
| LPA(18:0)-H | 0.0273 | 0.0062 | 0.0759 | 0.0420 | 0.2967 | 0.5277 | 0.4854 | |
| **TOTAL LPA** | **0.6705** | | **0.7678** | | | | | |
| TOTAL | 100.0000 | | 100.0000 | | | | | |

***Table S3 end***

### Table S4. Abundances of phospholipids relevant to Fig. 4c.

| | ABUNDANCE (% of Total) | | | | | | Red = p.value between 0.05-0.01. | | | |
|---|---|---|---|---|---|---|---|---|---|---|
| | vector | | TamAB | | TamAB$_{I1102R}$ | | Sorted based on TamAB means. | | | |
| Lipid | Mean | SEM | Mean | SEM | Mean | SEM | p.value | -LOG10(p) | FDR | Alias |
| CL(15:0_16:0_16:1_17:1)-H | 0.6052 | 0.0707 | 1.5603 | 0.1168 | 1.0469 | 0.0981 | 0.0002 | 3.6243 | 0.0120 | j |
| CL(16:0_16:0_16:1_18:1)-H | 0.8719 | 0.0816 | 1.3807 | 0.0528 | 1.2682 | 0.1586 | 0.0203 | 1.6919 | 0.0915 | |
| CL(16:0_16:0_16:1_17:1)-H | 0.7627 | 0.0504 | 1.1019 | 0.0362 | 1.1617 | 0.1131 | 0.0089 | 2.0500 | 0.0494 | o |
| CL(16:0_16:0_16:0_16:1)-H | 0.2841 | 0.0224 | 0.6534 | 0.0508 | 0.4909 | 0.0502 | 0.0007 | 3.1731 | 0.0120 | p |
| CL(14:0_16:0_16:0_16:1)-H | 0.1850 | 0.0243 | 0.6559 | 0.0956 | 0.3615 | 0.0343 | 0.0012 | 2.9223 | 0.0123 | k |
| CL(16:0_16:0_17:1_18:1)-H | 0.4813 | 0.0410 | 0.4588 | 0.0329 | 0.6153 | 0.0721 | 0.1208 | 0.9179 | 0.2559 | |
| CL(15:0_16:0_16:0_16:1)-H | 0.2048 | 0.0187 | 0.4413 | 0.0882 | 0.3359 | 0.0308 | 0.0412 | 1.3856 | 0.1399 | |
| CL(14:0_16:0_16:1_17:1)-H | 0.2164 | 0.0188 | 0.2971 | 0.1248 | 0.3086 | 0.0593 | 0.6881 | 0.1624 | 0.8397 | |
| CL(16:0_16:0_16:0_17:1)-H | 0.1872 | 0.0131 | 0.2598 | 0.0258 | 0.2898 | 0.0188 | 0.0145 | 1.8394 | 0.0695 | |
| CL(16:0_16:0_18:1_18:1)-H | 0.2766 | 0.0338 | 0.2831 | 0.0217 | 0.3178 | 0.0394 | 0.6426 | 0.1921 | 0.8261 | |
| CL(14:0_16:0_16:1_16:1)-H | 0.1058 | 0.0061 | 0.2474 | 0.0248 | 0.1788 | 0.0136 | 0.0007 | 3.1381 | 0.0120 | l |
| CL(16:0_16:0_16:0_18:1)-H | 0.1425 | 0.0121 | 0.2063 | 0.0129 | 0.2108 | 0.0254 | 0.0428 | 1.3690 | 0.1399 | |
| CL(48:2)-H | 0.0428 | 0.0010 | 0.1419 | 0.0270 | 0.0548 | 0.0149 | 0.0067 | 2.1750 | 0.0438 | m |
| CL(16:0_16:1_16:1_18:1)-H | 0.1438 | 0.0213 | 0.2309 | 0.0175 | 0.1828 | 0.0181 | 0.0305 | 1.5164 | 0.1154 | |
| CL(16:0_16:1_18:1)-H | 0.0624 | 0.0016 | 0.1246 | 0.0235 | 0.0666 | 0.0199 | 0.0634 | 1.1982 | 0.1684 | |
| **TOTAL CL** | **4.5724** | | **8.0434** | | **6.8905** | | | | | |
| PE(16:0_17:1)-H | 23.7786 | 0.7168 | 22.7750 | 0.7077 | 23.6354 | 1.3653 | 0.7430 | 0.1290 | 0.8665 | |
| PE(16:0_16:1)-H | 16.7467 | 1.1339 | 19.9893 | 0.6434 | 16.0208 | 1.4247 | 0.0713 | 1.1468 | 0.1684 | |
| PE(16:0_18:1)-H | 9.8418 | 0.1364 | 9.1273 | 0.3103 | 8.5959 | 0.4323 | 0.0607 | 1.2166 | 0.1684 | |
| PE(14:0_16:0)-H | 4.0849 | 0.4561 | 5.3678 | 0.2880 | 4.4811 | 0.2691 | 0.0725 | 1.1398 | 0.1684 | |
| PE(16:0_16:0)-H | 2.5294 | 0.1219 | 3.4597 | 0.1467 | 2.8863 | 0.1632 | 0.0045 | 2.3516 | 0.0320 | a |
| PE(15:0_16:0)-H | 3.1914 | 0.1528 | 2.6261 | 0.5772 | 2.6573 | 0.6027 | 0.6687 | 0.1747 | 0.8302 | |
| PE(16:0_14:1)-H | 2.4141 | 0.1341 | 2.6159 | 0.3113 | 2.4829 | 0.1326 | 0.7928 | 0.1009 | 0.8781 | |
| PE(15:0_16:1)-H | 5.2587 | 0.9469 | 2.1863 | 0.4015 | 4.1536 | 1.1237 | 0.0929 | 1.0321 | 0.2090 | |
| PE(16:1_18:1)-H | 2.3336 | 0.1583 | 1.8487 | 0.1056 | 1.8908 | 0.1227 | 0.0509 | 1.2937 | 0.1592 | |
| PE(16:0_19:1)-H | 1.6336 | 0.2047 | 1.3844 | 0.1803 | 1.6694 | 0.3032 | 0.6603 | 0.1803 | 0.8302 | |
| PE(16:1_17:1)-H | 1.5853 | 0.0958 | 1.3256 | 0.0734 | 1.4042 | 0.0840 | 0.1403 | 0.8531 | 0.2885 | |
| PE(17:1_18:1)-H | 1.3439 | 0.0130 | 0.9389 | 0.0458 | 1.1221 | 0.0750 | 0.0012 | 2.9277 | 0.0123 | e |
| PE(17:1_17:1)-H | 0.8101 | 0.0872 | 0.6860 | 0.1170 | 0.7364 | 0.1316 | 0.7461 | 0.1272 | 0.8665 | |
| PE(14:0_15:0)-H | 0.9847 | 0.1154 | 0.7016 | 0.0509 | 0.7693 | 0.1159 | 0.1631 | 0.7875 | 0.3174 | |
| PE(14:0_14:0)-H | 0.4136 | 0.0871 | 0.5978 | 0.1283 | 0.5353 | 0.1253 | 0.5392 | 0.2683 | 0.7325 | |
| **TOTAL PE** | **76.9505** | | **75.6304** | | **73.0409** | | | | | |
| PG(16:0_16:1)-H | 4.9949 | 0.1445 | 4.7698 | 0.2262 | 6.0834 | 0.5592 | 0.0620 | 1.2074 | 0.1684 | |
| PG(16:0_18:1)-H | 2.9357 | 0.1233 | 2.0769 | 0.1222 | 2.8088 | 0.2882 | 0.0252 | 1.5985 | 0.1068 | |
| PG(16:0_17:1)-H | 1.9872 | 0.2057 | 1.7657 | 0.1638 | 2.8265 | 0.4117 | 0.0591 | 1.2282 | 0.1684 | |
| PG(32:0)-H | 0.8498 | 0.0905 | 0.9800 | 0.1040 | 1.0987 | 0.1839 | 0.4478 | 0.3489 | 0.6593 | |
| PG(14:0_16:0)-H | 0.6082 | 0.1208 | 0.8638 | 0.1087 | 0.7772 | 0.1011 | 0.2992 | 0.5240 | 0.4938 | |
| PG(17:0_16:1)-H | 1.2101 | 0.1147 | 0.5238 | 0.0979 | 0.9710 | 0.2769 | 0.0685 | 1.1641 | 0.1684 | |
| PG(15:0_16:0)-H | 0.6502 | 0.0501 | 0.7581 | 0.2084 | 0.7816 | 0.1912 | 0.8395 | 0.0760 | 0.8901 | |
| PG(15:0_16:1)-H | 0.7381 | 0.0223 | 0.5899 | 0.1440 | 0.8501 | 0.2235 | 0.5137 | 0.2893 | 0.7113 | |
| PG(16:0_14:1)-H | 0.3145 | 0.0320 | 0.3260 | 0.0438 | 0.4167 | 0.0620 | 0.2987 | 0.5248 | 0.4938 | |
| PG(16:0_19:1)-H | 0.3782 | 0.0521 | 0.2413 | 0.0225 | 0.3935 | 0.0756 | 0.1501 | 0.8235 | 0.3003 | |
| PG(34:2)-H | 0.4771 | 0.0242 | 0.2774 | 0.0265 | 0.4268 | 0.0307 | 0.0015 | 2.8170 | 0.0137 | i |
| PG(33:0)-H | 0.1952 | 0.0149 | 0.1185 | 0.0309 | 0.1619 | 0.0407 | 0.2612 | 0.5830 | 0.4702 | |
| PG(36:2)-H | 0.4168 | 0.0342 | 0.2007 | 0.0150 | 0.3226 | 0.0252 | 0.0008 | 3.0874 | 0.0120 | h |
| PG(36:0)-H | 0.0994 | 0.0146 | 0.0896 | 0.0206 | 0.0854 | 0.0181 | 0.8530 | 0.0691 | 0.8901 | |
| PG(13:0_15:0)-H | 0.0882 | 0.0127 | 0.1060 | 0.0128 | 0.1055 | 0.0302 | 0.7852 | 0.1050 | 0.8781 | |
| **TOTAL PG** | **15.9438** | | **13.6874** | | **18.1097** | | | | | |
| PC(34:0)+H | 0.0282 | 0.0017 | 0.0240 | 0.0062 | 0.0276 | 0.0072 | 0.8486 | 0.0713 | 0.8901 | |
| PC(32:0)+H | 0.0226 | 0.0083 | 0.0212 | 0.0185 | 0.0293 | 0.0255 | 0.9482 | 0.0231 | 0.9482 | |
| PC(28:0)+H | 0.1371 | 0.0091 | 0.0979 | 0.0154 | 0.1049 | 0.0210 | 0.2319 | 0.6348 | 0.4280 | |
| PC(30:0)+H | 0.0086 | 0.0021 | 0.0108 | 0.0039 | 0.0046 | 0.0015 | 0.3086 | 0.5106 | 0.4938 | |
| PC(36:1)+H | 0.0086 | 0.0040 | 0.0047 | 0.0021 | 0.0026 | 0.0001 | 0.3058 | 0.5146 | 0.4938 | |
| PC(34:1)+H | 0.0606 | 0.0258 | 0.0361 | 0.0198 | 0.0164 | 0.0019 | 0.3004 | 0.5223 | 0.4938 | |
| PC(36:2)+H | 0.0275 | 0.0152 | 0.0277 | 0.0214 | 0.0088 | 0.0025 | 0.6168 | 0.2098 | 0.8075 | |
| PC(34:2)+H | 0.0433 | 0.0148 | 0.0436 | 0.0261 | 0.0153 | 0.0014 | 0.4487 | 0.3481 | 0.6593 | |
| **TOTAL PC** | **0.3364** | | **0.2659** | | **0.2096** | | | | | |

| | | | | | | | | | | |
|---|---|---|---|---|---|---|---|---|---|---|
| PA(16:0_16:1)-H | 0.1057 | 0.0149 | 0.1416 | 0.0170 | 0.1661 | 0.0313 | 0.2112 | 0.6753 | 0.4002 | |
| PA(16:1_18:1)-H | 0.0243 | 0.0052 | 0.0212 | 0.0024 | 0.0257 | 0.0064 | 0.8088 | 0.0922 | 0.8823 | |
| PA(14:0_14:0)-H | 0.0037 | 0.0008 | 0.0144 | 0.0121 | 0.0076 | 0.0024 | 0.5840 | 0.2336 | 0.7787 | |
| PA(14:0_16:1)-H | 0.0120 | 0.0019 | 0.0098 | 0.0013 | 0.0237 | 0.0053 | 0.0355 | 1.4497 | 0.1278 | |
| **TOTAL PA** | **0.1458** | | **0.1870** | **0.0328** | **0.2231** | | | | | |
| LPE(16:1)-H | 0.5364 | 0.0383 | 0.6975 | 0.0690 | 0.3700 | 0.0548 | 0.0078 | 2.1058 | 0.0470 | q |
| LPE(16:0)-H | 0.2549 | 0.0104 | 0.3869 | 0.0296 | 0.2153 | 0.0490 | 0.0138 | 1.8598 | 0.0695 | |
| LPE(18:1)-H | 0.2128 | 0.0084 | 0.2180 | 0.0241 | 0.1355 | 0.0333 | 0.0691 | 1.1606 | 0.1684 | |
| LPE(14:0)-H | 0.0554 | 0.0056 | 0.0888 | 0.0110 | 0.0469 | 0.0111 | 0.0296 | 1.5288 | 0.1154 | |
| LPE(17:0)-H | 0.0620 | 0.0080 | 0.0411 | 0.0083 | 0.0380 | 0.0069 | 0.1106 | 0.9561 | 0.2414 | |
| LPE(15:0)-H | 0.0565 | 0.0024 | 0.0324 | 0.0060 | 0.0257 | 0.0040 | 0.0019 | 2.7164 | 0.0154 | s |
| LPE(19:1)-H | 0.0302 | 0.0042 | 0.0275 | 0.0052 | 0.0203 | 0.0063 | 0.4336 | 0.3629 | 0.6593 | |
| LPE(18:0)-H | 0.0087 | 0.0016 | 0.0101 | 0.0030 | 0.0057 | 0.0013 | 0.3502 | 0.4557 | 0.5481 | |
| LPE(15:1)-H | 0.0181 | 0.0023 | 0.0070 | 0.0005 | 0.0076 | 0.0011 | 0.0008 | 3.0806 | 0.0120 | t |
| **TOTAL LPE** | **1.2351** | | **1.5093** | | **0.8649** | | | | | |
| LPC(19:0)+H | 0.1063 | 0.0179 | 0.0886 | 0.0141 | 0.1010 | 0.0209 | 0.7769 | 0.1097 | 0.8781 | |
| LPC(16:0)+H | 0.0157 | 0.0017 | 0.0147 | 0.0025 | 0.0132 | 0.0022 | 0.7140 | 0.1463 | 0.8568 | |
| LPC(18:0)+H | 0.0039 | 0.0012 | 0.0028 | 0.0011 | 0.0047 | 0.0008 | 0.4790 | 0.3197 | 0.6802 | |
| LPC(18:1)+H | 0.0037 | 0.0005 | 0.0035 | 0.0002 | 0.0033 | 0.0008 | 0.8857 | 0.0527 | 0.9110 | |
| **TOTAL LPC** | **0.1297** | | **0.1096** | | **0.1222** | | | | | |
| LPA(17:1)-H | 0.6406 | 0.0656 | 0.5293 | 0.1166 | 0.4890 | 0.0738 | 0.4818 | 0.3171 | 0.6802 | |
| LPA(18:0)-H | 0.0458 | 0.0168 | 0.0376 | 0.0238 | 0.0501 | 0.0265 | 0.9254 | 0.0337 | 0.9384 | |
| **TOTAL LPA** | **0.6864** | | **0.5669** | | **0.5391** | | | | | |
| TOTAL | 100.0000 | | 100.0000 | | 100.0000 | | | | | |

***Table S4 end***

**Table S5. Bacterial strains used in this study.**

| Strain Details | Source |
|---|---|
| *E. coli* K-12 NEB5α | New England Biolabs (cat#C2987H) |
| *E. coli* K-12 XL1 Blue | Agilent (cat#200236) |
| *E. coli* K-12 XL10 Gold | Agilent (cat#200315) |
| *E. coli* BL21(DE3) | Invitrogen (cat#C600003) |
| *E. coli* BL21(DE3)*ΔdsbA* | This study |
| *E. coli* K-12 W3110 wild-type (WT) | Provided by M Stephen Trent |
| W3110*ΔyhdP* | Douglass et al., 2022[10] |
| W3110*ΔtamB* | Douglass et al., 2022[10] |
| W3110*ΔtamBΔyhdP* | Douglass et al., 2022[10] |
| W3110*ΔtamA* | This study |
| W3110*ΔtamAB* | This study |
| W3110*ΔydbH* | This study |
| W3110*ΔtamAΔydbH* | This study |
| W3110*ΔtamBΔydbH* | This study |
| W3110*ΔtamABΔydbH* | This study |
| W3110*ΔydbHΔyhdP* | This study |
| W3110*ΔtamABΔyhdP* | This study |
| W3110*ΔtamAΔyhdP* | This study |
| W3110*ΔtamABΔyhdPΔrcsF*::$Km^R$ | This study |

**Table S6. Plasmids used and constructed in this study.**

| Plasmid | Description | Source |
|---|---|---|
| pTrc99a | IPTG-inducible plasmid, ampicillin resistance | Novopro |
| pJH114 | pTrc99a::BamABCDE$^{His}$ | Roman-Hernandez et al., 2014[11] |
| pXW47 | pTrc99a::$^{His}$TamAB | Wang et al., 2024[12] |
| pMSP1D1 | pET28a::$^{His\text{-}TEV}$MSP1D1 | Denisov et al., 2004[13] |
| pMTD1590 | pTrc99a::BamABCDE-$^{His}$TamAB | This study |
| pMTDS80 | pTrc99a::BamABCDE-$^{His}$TamA-TamB$^{TS}$<br>(TS-tag between 836-837 codons) | This study |
| pMTDS93 | pTrc99a::BamABCDE$^{His}$TamA-TamB490$^{TS}$<br>(Encodes TamB aa 837-1259 with N-terminal TS-tag) | This study |
| pMTDS116 | pTrc99a::BamABCDE-$^{His}$TamA-TamB$_{E1258C}^{TS}$ | This study |
| pMTDS206 | pTrc99a::BamABCDE-$^{His}$TamA-TamB$_{Q1256C}^{TS}$ | This study |
| pMTDS343 | pTrc99a::BamABCDE-$^{His}$TamA-TamB$_{L1254C}^{TS}$ | This study |
| pMTDS653 | pTrc99a::BamABCDE-$^{His}$TamA$_{T267C}$-TamB$_{E1258C}^{TS}$ | This study |
| pMTDS662 | pTrc99a::BamABCDE-$^{His}$TamA$_{G273C}$-TamB$^{TS}$ | This study |
| pMTDS663 | pTrc99a::BamABCDE-$^{His}$TamA$_{T267C}$-TamB$^{TS}$ | This study |
| pMTDS665 | pTrc99a::BamABCDE-$^{His}$TamA-TamB$_{T1185C}^{TS}$ | This study |
| pMTDS684 | pTrc99a::BamABCDE-$^{His}$TamA$_{E269C}$-TamB$^{TS}$ | This study |
| pMTDS685 | pTrc99a::BamABCDE-$^{His}$TamA$_{G169C}$-TamB$^{TS}$ | This study |
| pMTDS686 | pTrc99a::BamABCDE-$^{His}$TamA$_{G271C}$-TamB$^{TS}$ | This study |
| pMTDS688 | pTrc99a::BamABCDE-$^{His}$TamA$_{T270C}$-TamB$^{TS}$ | This study |
| pMTDS689 | pTrc99a::BamABCDE-$^{His}$TamA$_{S151C}$-TamB$^{TS}$ | This study |
| pMTDS691 | pTrc99a::BamABCDE-$^{His}$TamA-TamB$_{D1252C}^{TS}$ | This study |
| pMTDS692 | pTrc99a::BamABCDE-$^{His}$TamA$_{E269C}$-TamB$_{Q1256C}^{TS}$ | This study |
| pMTDS693 | pTrc99a::BamABCDE-$^{His}$TamA$_{S151C}$-TamB$_{T1185C}^{TS}$ | This study |
| pMTDS710 | pTrc99a::BamABCDE-$^{His}$TamA$_{G169C}$-TamB$_{V1008C}^{TS}$ | This study |
| pMTDS711 | pTrc99a::BamABCDE-$^{His}$TamA-TamB$_{V1008C}^{TS}$ | This study |
| pMTDS713 | pTrc99a::BamABCDE-$^{His}$TamA$_{G271C}$-TamB$_{L1254C}^{TS}$ | This study |
| pMTDS787 | pTrc99a::BamABCDE-$^{His}$TamA$_{T270C}$-TamB$_{Q1256C}^{TS}$ | This study |
| pMTDS842 | pTrc99a::BamABCDE-$^{His}$TamA$_{G273C}$-TamB$_{D1252C}^{TS}$ | This study |
| pMTDS918 | pTrc99a::BamABCDE-$^{His}$TamA-TamB$_{I1102R}^{TS}$ | This study |

**Table S7. Synthetic dsDNA fragments used in this study.**

| | | |
|---|---|---|
| mtd36 | GTTTTACATCGGTCTGGGGCCAGAATTATAACAGGATCCCGAAAAGGAATCAA<br>GAATTCATGGCGATGAAAAAACTGCTGATTGCGAGCCTGCTGTTTAGCAGCGC<br>GACCGTGTATGGCGCGAGCGCGTGGTCTCACCCGCAGTTCGAAAAGGGTGGCG<br>GGAGCGGTGGCGGTAGCGGTGGCTCCGCGTGGAGCCATCCGCAGTTTGAAAAG<br>GGTGGCTCCAATCTTGGTGGCAACGTCAATATCCGTAAC | To insert TS-tag into pMTD1590 to make pMTDS93 |
| mtd37 | GTGCAGGTGACCGATCCGCAAGGCCGCCGTGGTGGTGGTTCTGCATGGTCTCA<br>CCCGCAGTTCGAAAAGGGTGGCGGGAGCGGTGGCGGTAGCGGTGGCTCCGCGT<br>GGAGCCATCCGCAGTTTGAAAAGGGTGAATTCGGCTCCGGTGGCGGGAGCAAT<br>CTTGGTGGCAACGTCAATATCCGTAAC | To insert TS-tag into pMTD1590 to make pMTDS80 |

**Table S8. ssDNA Oligonucleotides used in this study.**

| DNA | Sequence | Notes |
|---|---|---|
| 375bamF | GATTTGCTCTATCAGTTCGAGTTTTAGGATCCTCTAGAGTCGACCTGC | pJH114, F |
| 376bamR | GCGCACATTTTCTCCTGAATATCCTTAACCACCGTTACCACTCAGCG | pJH114, R |
| 377tamF | GGATATTCAGGAGAAAATGTGCGC | pXW47, F |
| 378tamR | CTAAAACTCGAACTGATAGAGCAAATC | pXW47, R |
| mtd1 | TAATCATCCGGCTCGTATAATGTG | pTrc99a, F |
| mtd2 | GGCATGGGGTCAGGTGG | pTrc99a, F |
| mtd20 | GCGTAAAGGTTATTTCGATAGCG | TamA primer, F |
| mtd27 | CGATCTGCCGGAAAGCGC | TamB primer, F |
| mtd28 | AACAGAAAACTGCATCGAAACCGGGGTCGGTTACTCTACG | TamA T267C, F |
| mtd29 | CGCGGCGAAACCACGCCC | TamA T267C, R |
| mtd30 | CTATCAGTTCTGCTTTTAGGATCCTCTAGAGTCGACC | TamB E1258C, F |
| mtd31 | AGCAAATCCAGTGCCTGGTCTAC | TamB E1258C, R |
| mtd33 | TTATAATTCTGGCCCCAGACCGATGTAAAAC | End *tamA*, R |
| mtd34 | AATCTTGGTGGCAACGTCAATATCCGTAAC | TamB 837 codon, F |
| mtd35 | ACGGCGGCCTTGCGGATCGGTC | TamB 836 codon, R |
| mtd393 | AAAGTCCTTAACCTGCATTGCGTTGCGTAAAGG | TamA S151C, F |
| mtd394 | ACGCAACGCAATGCAGGTTAAGGACTTTTTGAAATTTTCATAATCGCCC | TamA S151C, R |
| mtd395 | CAAAGCGCAGCTGTGCATTGCGCTCGGCCTGC | TamA G169C, F |
| mtd396 | CGAGCGCAATGCACAGCTGCGCTTTGGTAAATTCGCT | TamA G169C, R |
| mtd401 | CGCGAACAGAAAACTGCATCGAAACCGGGGTCG | TamA T267C, F |
| mtd402 | CCGGTTTCGATGCAGTTTTCTGTTCGCGGCGAAACC | TamA T267C, R |
| mtd403 | AGAAAACACCATCTGCACCGGGGTCGGTTACT | TamA E269C, F |
| mtd404 | CGACCCCGGTGCAGATGGTGTTTTCTGTTCGCGGCG | TamA E269C, R |
| mtd405 | AAACACCATCGAATGCGGGGTCGGTTACTCTACGG | TamA T270C, F |
| mtd406 | TAACCGACCCCGCATTCGATGGTGTTTTCTGTTCGCGGCG | TamA T270C, R |
| mtd407 | ACACCATCGAAACCTGCGTCGGTTACTCTACGG | TamA G271C, F |
| mtd408 | AGTAACCGACGCAGGTTTCGATGGTGTTTTCTGTTCGC | TamA G271C, R |
| mtd409 | CGAAACCGGGGTCTGCTACTCTACGGACGTGGGACC | TamA G273C, F |
| mtd410 | CCGTAGAGTAGCAGACCCCGGTTTCGATGGTGT | TamA G273C, R |
| mtd411 | CGGGGTCGGTTACTGCACGGACGTGGGACCGC | TamA S275C, F |
| mtd412 | CCACGTCCGTGCAGTAACCGACCCCGGTTTCGA | TamA S275C, R |
| mtd413 | CTCCAGCGATGTGTGCATGCTTAACGATAACCTGCAACCGG | TamB V1008C, F |
| mtd414 | CGTTAAGCATGCACACATCGCTGGAGACGCCT | TamB V1008C, R |
| mtd419 | GGTAAAATCGGCGAGTGCTTTGGCGTAAGCAATTTAGCGCTCG | TamB T1185C, F |
| mtd420 | GCTTACGCCAAAGCACTCGCCGATTTTACCCACAATCTGG | TamB T1185C, R |
| mtd423 | GCTATATCTGGAATGCGTGTCTGGTGTAGACCAGGCA | TamB A1243C, F |
| mtd424 | ACACCAGACACGCATTCCAGATATAGCTTAGGCATCAGGCG | TamB A1243C, R |
| mtd427 | AGACCAGGCACTGTGCTTGCTCTATCAGTTCGAGTTTTAGG | TamB D1252C, F |
| mtd428 | TGATAGAGCAAGCACAGTGCCTGGTCTACACCAGAC | TamB D1252C, R |
| mtd467 | TCAACCGTATCTTAATCGTGAAGCTATTCGTAACCCGGATGC | TamB I1102R, F |
| mtd468 | ACGAATAGCTTCACGATTAAGATACGGTTGATCTGGCGGACCA | TamB I1102R, R |

F = forward primer, R = reverse primer.

**Table S9. Antisera used in this study.**

| Antibodies | Notes | Source |
|---|---|---|
| Rabbit αBamA$_C$ | Raised against peptide CQPFKKYDGDKAEQFQFNIGKT | Pavlova et al., 2013[14] |
| Rabbit αTamA | Raised against peptide CPVADKDEHGLQFYIGLGPE | Wang et al., 2024[12] |
| Mouse αStrepII | Monoclonal antibody (#34850) | Qiagen |
| Rabbit αTamB | Raised against residues 963-1138 | Chris Stubenrauch |

**Table S10. Model statistics.**

| | $^{His}$TamAB490$^{TS}$ LMNG Detergent | $^{His}$TamAB490$^{TS}$ Lipid Nanodisc |
|---|---|---|
| **Data collection & processing** | | |
| Magnification | 105,000 | 105,000 |
| Voltage (kV) | 300 | 300 |
| Electron exposure ($e^-/Å^2$) | 58 | 72 |
| Defocus range (μm) | -0.1 to -1.7 | -0.2 to -2.3 |
| Pixel size (Å) | 0.83 | 0.83 |
| Symmetry imposed | C1 | C1 |
| Initial particle images (no.) | 3,448,506 | 6,730,439 |
| Final particle images (no.) | 80,583 | 103,937 |
| Map resolution (Å) | 3.51 | 3.71 |
| FSC threshold | 0.143 | 0.143 |
| **Refinement** | | |
| Initial model used (PDB code) | n/a | n/a |
| Model resolution (Å) | 3.65 | 3.80 |
| FSC threshold | 0.5 | 0.5 |
| Map-model CC | 0.7449 | 0.7283 |
| Model composition | | |
| Non-hydrogen atoms | 7627 | 7627 |
| Protein residues | 978 | 978 |
| R.M.S. deviations | | |
| Bond length (Å) | 0.005 | 0.007 |
| Bond angle (°) | 0.999 | 1.070 |
| Validation | | |
| MolProbity score | 1.17 | 1.23 |
| Clashscore | 0.99 | 1.98 |
| Rotamer outliers (%) | 0 | 0 |
| Ramachandran plot | | |
| Favoured (%) | 94.46 | 96.1 |
| Allowed (%) | 5.54 | 3.9 |
| Disfavoured (%) | 0.0 | 0.0 |

### VIDEO INFORMATION:

***3D variability analysis (3DVA) on particles in LMNG micelles:***

**Video S1-3. $^{His}$TamAB490$^{TS}$ micelle (full).**
Videos correspond to variability component 0, 1, and 2. TamA (orange) and TamB (dark-green).

**Video S4-6. $^{His}$TamAB490$^{TS}$ micelle (focus on α-helix 3).**
Videos correspond to variability component 0, 1, and 2. TamA (orange), TamB (dark-green), and TamB α-helix 3 (red). The model is also overlaid in this video.

**Video S7-9. $^{His}$TamAB490$^{TS}$ micelle (TamB β-taco).**
Videos correspond to variability component 0, 1, and 2. TamA is hidden. TamB (dark-green), surface corresponding to L1102 (purple), putative detergent densities (grey).

***3D variability analysis (3DVA) on particles in phospholipid nanodiscs:***

**Video S10-12. $^{His}$TamAB490$^{TS}$-nanodiscs (full).**
Videos correspond to variability component 0, 1, and 2. TamA (orange) and TamB (dark-green).

**Video S13-15. $^{His}$TamAB490$^{TS}$-nanodiscs (focus on α-helix 3).**
Videos correspond to variability component 0, 1, and 2. TamA (orange), TamB (dark-green), and TamB α-helix 3 (red). The model is also overlaid in this video.

**Video S16-18. $^{His}$TamAB490$^{TS}$-nanodiscs (TamB β-taco).**
Videos correspond to variability component 0, 1, and 2. TamA is hidden. TamB (dark-green), surface corresponding to L1102 (purple), putative phospholipid densities (grey).

***3DFlex analysis on particles in phospholipid nanodiscs:***

**Video S19-20. $^{His}$TamAB490$^{TS}$-nanodiscs (full).**
3DFlex conformational variability analysis videos corresponding to variability component 0 and 1. TamA (orange) and TamB (dark-green).
